# Müllerian mimicry of a quantitative trait despite contrasting levels of genomic divergence and selection

**DOI:** 10.1101/842708

**Authors:** Emma V. Curran, Sean Stankowski, Carolina Pardo-Diaz, Camilo Salazar, Mauricio Linares, Nicola J. Nadeau

## Abstract

Hybrid zones, where distinct populations meet and interbreed, give insight into how differences between populations are maintained despite gene flow. Studying clines in genetic loci and adaptive traits across hybrid zones is a powerful method for understanding how selection drives differentiation within a single species, but can also be used to compare parallel divergence in different species responding to a common selective pressure. Here, we study parallel divergence of wing colouration in the butterflies *Heliconius erato* and *H. melpomene*, which are distantly related Müllerian mimics that show parallel geographic variation in both discrete variation in pigmentation, and quantitative variation in structural colour. Using geographic cline analysis, we show that clines in these traits are positioned in the roughly the same geographic region for both species, which is consistent with direct selection for mimicry. However, the width of the clines varies markedly between species. This difference is explained in part by variation in the strength of selection acting on colour traits within each species, but may also be influenced by differences in the dispersal rate and total strength of selection against hybrids between the species. Genotyping-by-sequencing also revealed weaker population structure in *H. melpomene*, suggesting the hybrid zones may have evolved differently in each species; which may also contribute to the patterns of phenotypic divergence in this system Overall, we conclude that multiple factors are needed to explain patterns of clinal variation within and between these species, although mimicry has probably played a central role.

## Introduction

Hybrid zones, where genetically differentiated populations are in contact and interbreed, have long been a valuable resource in understanding the evolutionary processes shaping taxonomic boundaries (Barton & Gale, 1993; Endler, 1977). Hybrid zones can form in continuously distributed populations, where different alleles are favoured at either end of an ecological gradient, a process called primary intergradation (Endler, 1977). Alternatively, they can form when previously isolated populations, which have become genetically differentiated in allopatry, come into secondary contact (Endler, 1977). Both scenarios can lead to the formation of sharp geographic clines in quantitative traits and the loci that underlie them. These clines reflect the balance between gene flow and divergent selection (Barton & Hewitt, 1985), and their study can therefore provide deep insight into potential targets of natural selection.

Cline theory provides a powerful framework for studying patterns of variation across hybrid zones, enabling key biological parameters, including the strength and nature of selection shaping variation, to be estimated (Barton & Hewitt, 1985). By fitting geographic cline models to many loci or quantitative traits, it is possible to understand how the nature and the relative strength of selection varies among them. For example, assuming selection is acting across a sharp environmental gradient, the cline centre is indicative of the geographic location where the direction of divergent selection switches. In this case, if clines are centred at the same location (henceforth referred to as cline coincidence), this indicates that a suite of traits and loci are all affected by a common selective agent, or multiple agents that coincide geographically (Barton and Hewitt 1985). Variation in cline width can be used to make inferences about the strength of selection acting on a locus or trait, with narrower clines indicating stronger selection, all else being equal. The overall shape of clines is also informative about the nature of selection shaping the cline. For example, if variation at a trait or locus is shaped only by direct selection, the cline is predicted to have a sigmoidal shape (Barton and Hewitt 1985; Barton and Gale 1993). However, if the strength of direct selection on each locus is outweighed by indirect selection from many loci in linkage disequilibrium (LD), the total selection affecting each locus in LD will be approximately equal (Kruuk, Baird, Gale, & Barton, 1999; J M Szymura & Barton, 1991). This can result in many clines with similar centres and widths, and with steeper centres than would be expected from direct selection alone, referred to as “stepped” clines (Barton & Hewitt, 1985; Vines et al., 2016)

Despite being primarily used to study patterns of trait and marker variation within a single species, cline analysis may also be used to understand how closely related species are shaped by the same extrinsic selection pressure (e.g. Mallet et al., 1990). For example, if multiple closely related and ecologically similar species are distributed across the same habitat transition, local adaptation may cause similar traits to diverge in concert. Although examples of parallel adaptation can demonstrate striking convergence, the extent of trait divergence within each species, and extent of parallelism between them, may vary depending on a host of factors. For example, at White Sands, New Mexico, three lizard species show strong divergence in their dorsal colour, an adaptation that improves crypsis on different soil types (Rosenblum & Harmon, 2011). However, the extent of colour divergence varies between the species for reasons that are not entirely clear (Rosenblum & Harmon 2011). Cline analysis can be used to precisely quantify differences in the geographical distribution and variation of putative adaptive traits between species. When combined with genome-wide data, this can provide insight into factors influencing the degree of parallelism. Differences between species in intrinsic factors, such as their dispersal rate, population density, variation in their past demographic histories, and the genetic architecture of traits, could alter patterns of clinal variation in adaptive traits between species that are otherwise subject to the same extrinsic selection pressures.

Here, we studied a case of parallel divergence in the Müllerian co-mimics *Heliconius erato* and *Heliconius melpomene*. We examined clinal variation in two colour pattern traits: the yellow hindwing bar and iridescence. Where the pair co-occur, they converge on almost identical patterns to share the cost of educating predators of their distastefulness (Brown, 1981). Both species comprise many parapatric colour pattern races, or subspecies, connected by hybrid zones (Mallet, 1993; Rosser, Dasmahapatra, & Mallet, 2014). When different subspecies hybridise, their offspring can display novel or heterozygous phenotypes (Arias et al., 2008; Mallet, 1989; Mallet, 1986). Predators are less likely to learn to avoid rare phenotypes, causing frequency-dependent selection on colour patterns (Langham, 2004; Mallet & Barton, 1989). This maintains stable hybrid zones (Mallet, 1986; Rosser et al., 2014). The diverse colouration seen in the *Heliconius* genus has been extensively studied, the vast majority of which is determined by a genetic ‘tool kit’ of five major-effect loci (S W Baxter, Johnston, & Jiggins, 2009; Kronforst et al., 2006; A. Martin et al., 2012; Nadeau, 2016; Nadeau et al., 2016; Reed et al., 2011; Westerman et al., 2018). Previous studies have found low levels of genetic differentiation between parapatric colour races, with a few diverged loci, mainly controlling colour pattern differences (Martin et al., 2013; Nadeau et al., 2014; Supple et al., 2013).

Near the Panamanian-Colombian border, there are co-occurring hybrid zones between subspecies of *H. erato* and *H. melpomene*, which differ in the presence of a yellow hindwing bar and in iridescent blue colouration (Mallet, 1986; Figure 1). Iridescence is produced by nano-structural ridges on the surface of wing scales, which are layered to produce constructive interference of blue light (Parnell et al., 2018). In a system so well-studied, little is known regarding selection on structural colour (Sweeney, Jiggins, & Johnsen, 2003). Divergence in this trait has not been previously studied. While the yellow bar is controlled by a single major-effect gene (Mallet, 1986; Nadeau, 2016), iridescence segregates as continuous variation, with conservative estimates suggesting it is controlled by around five additive genetic loci (Brien et al., 2018). While differences in pigment colouration across hybrid zones seem to be maintained by strong divergent selection, despite gene flow across the rest of the genome (Nadeau et al., 2014), it is unclear whether we would expect to see this in a more complex trait such as iridescence. Polygenic local adaptation may only require small allele frequency changes, but can also involve greater levels of covariance between loci (Le Corre & Kremer, 2012). The combined action of divergent selection and the build-up of statistical associations between loci can reduce effective migration rates across the genome (Flaxman, Wacholder, Feder, & Nosil, 2014; Kruuk et al., 1999). Therefore, an increased level of overall genome-wide differentiation, and population level genetic structure may be expected across hybrid zones over which quantitative variation is maintained, particularly if the trait is highly polygenic.

**Figure 1.**
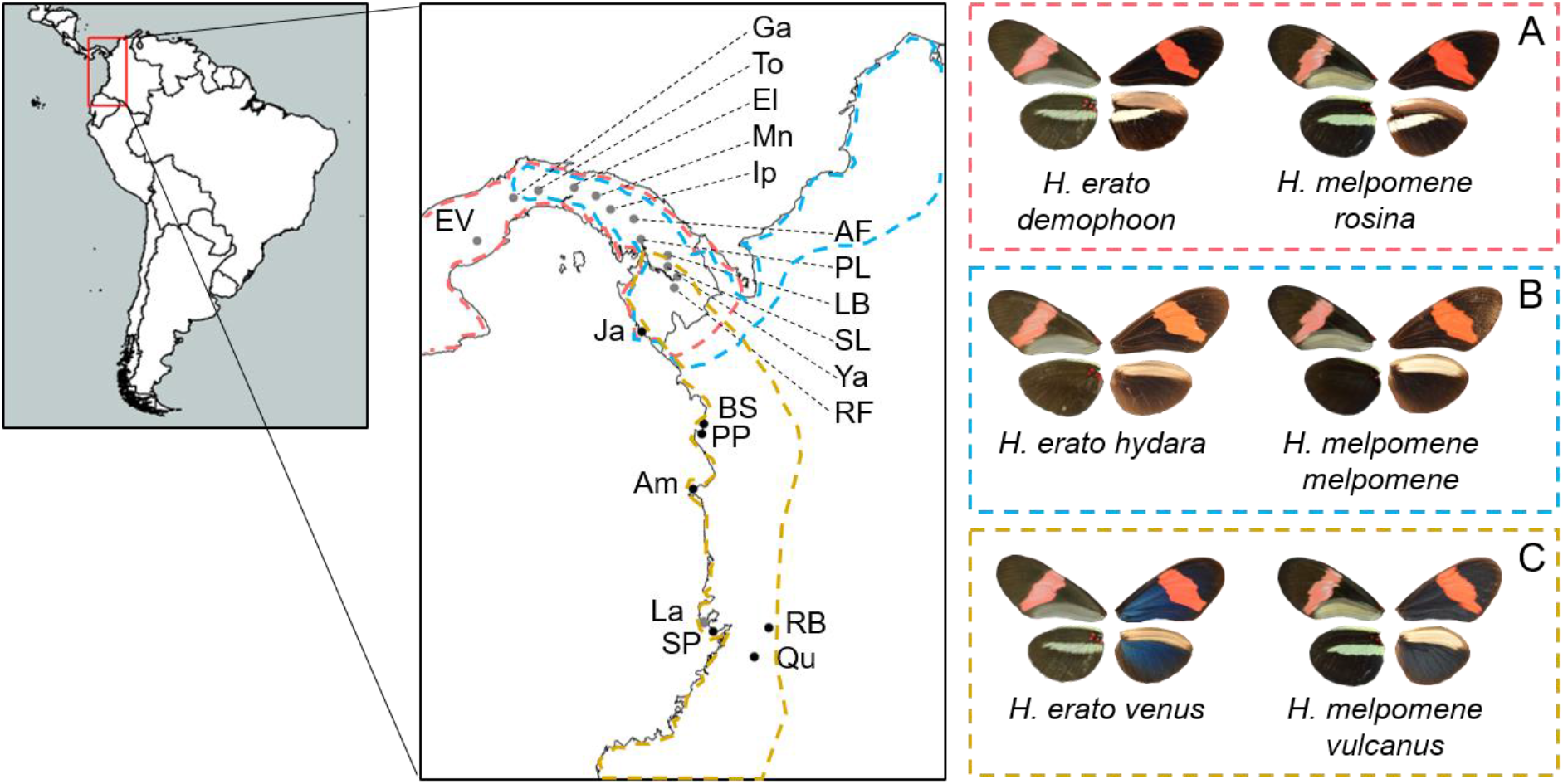
Sampled populations in Colombia and Panama. Sites are labelled with abbreviations (further information about sites and collections are in Table S1). Photographs show the phenotypes of mimetic races of *H. erato* and *H. melpomene* from Central America (**A**), North Colombia (**B**), and Western Colombia (**C**). For each pictured phenotype, the wings on the left-hand side show the ventral wing pattern, and the wings on the right-hand side show the dorsal wing pattern. Approximate ranges for the mimetic race pairs are outlined with dashed lines (Rosser, Phillimore, Huertas, Willmott, & Mallet, 2012). Populations that are included in the phenotypic analysis only are shown in grey, populations that are included in both the phenotypic and genetic analysis are shown in black.

Here, we use geographic cline analysis to examine the selection regimes impacting variation and convergence of iridescence and other traits, both within and between the co-mimics *H. erato* and *H. melpomene*. Within species, we are primarily interested in understanding how the genetic basis of these traits has influenced their divergence across the hybrid zone. Because iridescence is polygenic, it may be more difficult for direct selection to maintain strong trait divergence along the cline compared with the yellow bar trait, which has a simple genetic basis and is highly visible to selection. Between the species, our aim is to compare the extent of parallel divergence in the co-mimics. Due to the strong existing evidence that mimicry drives colour pattern convergence between *H. erato* and *H. melpomene*, our null expectation would be that clines in colour traits should be very similar in position and shape. Any deviations from this expectation would suggest that either selection is acting differently on iridescence in each species, or that some other factor has affected the extent of divergence within each species.

## Methods

### Butterfly specimens

*Heliconius melpomene* and *Heliconius erato* individuals were collected from several sites in the Chocó-Darien ecoregion between the Andes and the Pacific in Colombia, and part way across the isthmus of Panama (Figure 1, SI Table S1). Wings were removed and stored in envelopes. Bodies were preserved in NaCl saturated 20% dimethyl sulfoxide (DMSO) 0.25M EDTA.

### Sequencing data

Restriction-associated DNA (RAD) sequence data were generated for 265 *H. erato* (SI Table S2), and whole genome re-sequencing was carried out on 36 *H. melpomene* individuals (SI Table S3). Genomic DNA was extracted from each individual using DNeasy Blood and Tissue Kits (Qiagen). Library preparation and sequencing was carried out by Edinburgh Genomics (University of Edinburgh).

Single-digest RAD libraries were prepared using the *Pst1* restriction enzyme, with eight base-pair barcodes and sequenced on the Illumina HiSeq 2500 platform (v4 chemistry), generating an average of 554,826 125 base paired-end reads per individual (see SI Table S2 for coverage and accession information). We demultiplexed the pooled reads using the RADpools program in the RADtools package version 1.2.4 (Baxter et al., 2011).

For the whole-genome sequencing, TruSeq Nano, gel-free libraries were prepared from genomic DNA samples of 36 *H. melpomene* individuals and sequenced on Illumina HiSeq 2500 platform (v4 chemistry), generating an average of 31,484,363 125 base paired-end reads per individual (see SI Table S3 for coverage and accession information).

### Data processing and variant calling

We checked the quality of all the raw sequencing reads using FastQC (v 0.11.5) and removed any remaining adapters using Trim Galore (v 0.4.1). We aligned the sequence data of all individuals, both RAD sequenced and WGS, to their corresponding reference genomes, either *Heliconius melpomene* version 2 (Davey et al., 2016) or *Heliconius erato* (Van Belleghem et al., 2017), obtained from lepbase (Challis, Kumar, Dasmahapatra, Jiggins, & Blaxter, 2016), using bowtie2 (v 2.3.2), with the local alignment option, and the very-sensitive pre-set parameter options to improve accuracy of the alignment. We used samtools (v 1.3.1) to sort and index the alignment files. We removed any duplicates that may have arisen during library preparation using the MarkDuplicates program in Picard tools (v 1.92).

Single nucleotide polymorphism (SNP) datasets were generated using samtools mpileup (v 1.5) to compute genotype likelihoods and bcftools (v 1.5) for variant calling. For a site to be a variant, the probability that it was homozygous for the reference allele across all samples was required to be less than 0.05. Multiallelic sites, insertions and deletions were ignored. For *H. melpomene* we identified 30,027,707 SNPs and for *H. erato* we identified 5,088,449 SNPs. We removed SNPs with a phred quality score lower than 30, that lacked sequence data in 50% or more of the individuals, that had a minor allele frequency lower than 0.05 or that were private variants. We pruned based on linkage disequilibrium, discarding SNPs within a 20kb window with *r*^*2*^ > 0.8, using the bcftools plugin ‘+prune’. This reduced the initial number of called SNPs down to 9,336,937 in *H. melpomene* and 159,405 in *H. erato*.

### Population structure

To examine population structure, we estimated the ancestry of each individual using the software NGSadmix (Skotte, Korneliussen, & Albrechtsen, 2013), which estimates the proportion of each genome that can be attributed to predefined number of populations (*k*) using genotype likelihoods. For each species, NGSadmix was run for a range of values of *k*, one to ten, each being replicated ten times with a random seed. The value of *k* best describing the population structure was determined using the ∆*k* criterion (Evanno, Regnaut, & Goudet, 2005), implemented in CLUMPAK (Kopelman, Mayzel, Jakobsson, Rosenberg, & Mayrose, 2015).

We carried out a principal components analysis (PCA) using PCAngsd (Meisner & Albrechtsen, 2018), which estimates a covariance matrix based on genotype likelihoods. We used eigenvector decomposition to retrieve the principal components of genetic structure.

### Population differentiation

To test the extent of genetic differentiation between the iridescent and non-iridescent subspecies, we measured *F*_*ST*_ between all individuals from iridescent populations south of the hybrid zone, and all non-iridescent individuals north of the hybrid zone, excluding the sampling site Jaqué, which was in the centre of the hybrid zone in both species. In each species, the two non-iridescent colour pattern races were collapsed into a single “non-iridescent” group, north of the hybrid zone, since our results show there is no genetic structure between them based on race. SNP datasets were generated for each species, using samtools mpileup and bcftools (v 1.5). In each species Hudson’s *F*_*ST*_ estimator was calculated among populations (Hudson, Slatkin, & Maddison, 1992):

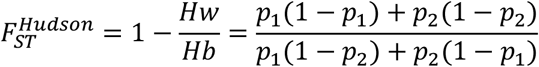

Where *Hw* is the within-population heterozygosity, *Hb* is the between-population heterozygosity, and *p*_*1*_ and *p*_*2*_ represent the allele frequencies in each population. This was calculated in *R* for every SNP with a custom script. Average genome-wide *F*_*ST*_ was calculated as a ratio of averages, by averaging the variance components, *Hw* and *Hb*, separately, as recommended by Bhatia et al. (2013) We also estimated average genome-wide *F*_*ST*_ between all pairs of populations, including those in the hybrid zone, for each species, and plotted pairwise *F*_*ST*_ against pairwise geographic distance.

### Phenotypic measurements

Digital images of butterfly wings were taken with a Nikon D7000 DSLR camera fitted with an AF-S DX Micro NIKKOR 40 mm f/2.8G lens (Nikon UK Ltd., Surrey, UK), mounted on an adjustable platform. Standardised lighting conditions were achieved using two external natural daylight fluorescent lights, mounted to illuminate at 45 degrees from incident, to maximise brightness of observed iridescent colour. Photographs were taken with a shutter speed of 1/60 sec and an aperture of f/10. Each sample was photographed with an X-Rite colorchecker passport (X-Rite, Inc., MI, USA) in shot. The Nikon raw (.NEF) image files were converted to standard raw files (.DNG) using Adobe DNG converter (Adobe Systems Inc., USA). The RGB channels in the images were then linearized using the neutral grey scale on the colorchecker using GNU Image Manipulation Program, v2.8.

The mean RGB values from regions in the discal cell on the right forewing and the Cu_2_ cell on the right hindwing were measured (SI Figure S1A). If the wings on the right-hand side showed damage, wings on the left-hand side were used. Wing regions were selected using the polygon selection tool in ImageJ, version 1.50b (Abràmoff, Magalhães, & Ram, 2004), and mean RGB scores were measured using the Color Histogram plugin. To minimise variation in blue colour due to age and wing wear, we excluded individuals with extensive wing wear or damage.

We tested for repeatability (Whitlock & Schluter, 2009) of the RGB values on 26 individuals photographed a second time under the same conditions on a different day, with a second set of RGB measurements taken. These individuals were selected from regions in which varying levels of iridescence is seen (20 individuals from Valle del Cauca, Colombia, and 6 individuals from Darién, Panama). Variance among individuals was calculated by taking the difference between the group mean square and the error mean square, and dividing it by the number of replicates. These components of variance were extracted from a general linear model in R v3.2.3 (R Core Team, 2015). The fraction of total variance that is due to true differences between individuals was then calculated by dividing the variance among individuals by the total variance.

A measure of relative blue reflectance (blue score) was determined for each individual by taking the mean blue channel value (B) and the mean red channel value (R) for both wing regions and calculating:

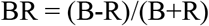

This gives a standardised score of how blue an individual is, with BR = 1 being the ‘bluest’, and BR = −1 being the ‘reddest’ (Figure S1 B, C).

### Estimation of ‘yellow bar’ allele frequencies

Allele frequencies for the yellow hindwing bar were estimated based on phenotype for both species. This was done for all sampling sites in Colombia and Panama with five or more individuals. The ‘yellow bar’ phenotype was scored categorically according to Mallet (1986), who showed that this phenotype segregates in the same way for both *Heliconius erato* and *H. melpomene*. Variation in the yellow bar across this hybrid zone is controlled by three alleles: The North Colombian yellow bar allele (Y), the West Colombian yellow bar allele (y_wc_) and the Central American yellow bar allele (y_ca_). Individuals of both species with a yellow bar on both sides of the wing (Figure 1A) have genotype y_ca_y_ca_. Individuals lacking a yellow bar (Figure 1B) have genotype YY. Individuals with the “shadow bar” phenotype, where the outline of the bar can be seen on the underside of the hindwing without any yellow pigment, and without a bar on the upper side of the hindwing, have genotype Yy_wc_ or Yy_ca_. Individuals with a yellow bar on the underside of the hindwing (Figure 1C) have genotype y_wc_y_ca_ or y_wc_y_wc_. As two of the four phenotypes can be produced by two different allele combinations we inferred the allele frequencies at each locality for each species assuming Hardy-Weinberg equilibrium for the three alleles. The frequency of Y could be directly observed from both its heterozygous and homozygous phenotypes. The frequency of y_ca_ could be inferred from the frequency of its homozygous phenotype, allowing us to infer the frequency of y_wc_. We focus on the y_wc_ allele for the remainder of the paper, as this underlies the yellow bar phenotype seen in the iridescent forms of both species, and appears to be lost across the same hybrid zone over which iridescence is lost. This provides us with the opportunity to directly compare clines in Mendelian and polygenic traits.

### Geographic cline analysis

We used geographic cline analysis to model patterns of clinal variation in (*i*) the mean iridescence score, (ii) frequency of the yellow bar allele (y_wc_), and (iii) mean admixture, estimated using NGSadmix, across both hybrid zones. We assumed two parental populations here, as our analyses of population structure reveal two genetic clusters in each species, despite there being three overlapping colour pattern races. Specifically, we fitted three alternative geographic cline models (Szymura & Barton, 1991; Szymura & Barton, 1986) using ANALYSE v1.30 (Barton & Baird, 2002). Sampling sites with fewer than five individuals were excluded from the cline analyses, leaving 529 *H. erato* and 126 *H. melpomene*. Blue scores were normalised to a new range of 0 to 1 (Fig S1 B, C) as required by the software. Distances between sampling sites were estimated using the great circle distance, calculated using the ‘hzar.map.greatCircleDistance’ function in the R package HZAR (Derryberry, Derryberry, Maley, & Brumfield, 2014).

ANALYSE fits cline models to marker loci and/or quantitative trait data, and can be used to compare the fit of three alternative cline models to either population means (used for iridescence and admixture scores) or frequency data (used for the yellow bar allele). The simplest model is a sigmoid cline described by a hyperbolic tangent (Szymura & Barton, 1986). The other two more complex models are ‘stepped’ clines. They consist of a central sigmoid step flanked by two exponential tails that describe the pattern of introgression from the centre into the foreign genepool; θ is the rate of decay, and the strength of the barrier to gene flow, *B*, can be estimated as the ratio between the difference in the allele frequency and the initial gradient in allele frequency with distance *x* at the edges of the central segment. In the symmetrical ‘Sstep’ model, θ and *B* are equal on both sides. In the asymmetrical ‘Astep’ model, the pattern of introgression is different on the left and right side.

ANALYSE uses the Metropolis algorithm to search the likelihood surface to find the ML solution to the model. To ensure that the likelihood surface was thoroughly explored, independent runs were conducted using a range of initial parameter estimates. After obtaining maximum likelihood (ML) solutions for the three cline models, the most likely model was identified using Likelihood Ratio Tests. As the minimum and maximum mean allele frequency or trait values (p(*z*)_min_, p(*z*)_max_) were allowed to vary in the tails of the cline, the sigmoid, Sstep and Astep models were described by 2 (*c*, *w*), 4 (*c*, *w*, θ, *B*) and 6 parameters (*c*, *w*, θ_0_, θ_1_, *B*_0_, *B*_1_), respectively.

After model selection, support limits were estimated for each parameter in the ML model. Starting with the optimum fit, and constraining the values of all other parameters, the likelihood surface for individual parameters were explored by making 10,000 random changes of their value. The range of estimates that was within 2 log-likelihood units of the maximum estimate was taken as the support limit for that parameter, and is approximately equivalent to a 95% confidence interval.

Coincidence of cline centres (*c*) and concordance of cline widths (*w*) were tested using the composite likelihood method (Kawakami, Butlin, Adams, Paull, & Cooper, 2009; Phillips, Baird, & Moritz, 2004). The method involves obtaining a composite ML score for a given parameter (ML_comp_) and comparing it with the sum of the ML estimates obtained for each profile (ML_sum_). ML_comp_ was obtained by constructing a log-likelihood profile (10 km intervals for *c* and *w*, between 0 km and 1000 km) with all other parameters allowed to vary, summing the profiles, and obtaining the ML estimate; ML_sum_ was obtained by summing the ML estimates from each profile. If clines are not coincident or concordant, ML_sum_ is significantly smaller than ML_comp_, as determined by a chi-squared test (α = 0.05) with n-1 degrees of freedom, where n is the number of traits. One complication with this method for comparing cline parameters is that the profiles for each trait must be built using the same model. Although the more complex Sstep and Astep models are a significantly better fit than the sigmoid model, the parameters estimates for the cline centre and cline width were similar regardless of the model fit (see results).

Therefore, all likelihood profiling was conducted with the sigmoid model.

To estimate the strength of selection acting on y_wc_, the following equation was used from Barton and Gale (1993):

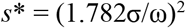

Where *s*\* is the difference in mean fitness between populations at the edge of the cline, and populations at the centre. This demonstrates the mean strength of effective selection on loci underlying a trait required to maintain a cline of width (*w*), given the dispersal distance per generation (σ). Dispersal estimates were taken from Mallet et al. (1990) and Blum (2002).

### Tests for concordance of clines using regression analysis

In addition to geographic cline analysis, we also used the regression procedure outlined in Nürnberger et al. (1995) as a method for testing for the concordance of clines within and between the species. Concordance of clines is predicted to result in a linear regression of population means or allele frequencies between characters *i* and *j*. Alternatively, non-concordant clines should show a deviation from linearity that can be described by a quadratic polynomial. We compared the fits of linear and quadratic models to each pair of characters, including the mean admixture score, frequency of the y_wc_ alleles and the mean iridescence score within a species using custom script in R. Because data were collected for each species in the same location, we could also use this analysis to compare clines in the same traits between *H. erato* and *H. melpomene*, with the exception of the admixture score because few sites included genetic data for both species. Significance of the deviation from linearity was determined by comparing the F-ratios of the quadratic and linear fits.

## Results

### Population Structure

We investigated population structure using genome-wide SNP data in the programs NGSadmix, to estimate ancestry proportions from a varying number of genetic clusters (*K*), and PCAngsd to confirm population clustering by principal components (PCA). This revealed different patterns of population structure between the co-mimics. In *H. erato*, NGSadmix supported *K*=2 (SI Figure S3B), representing a “Panama-like” and a “Colombia-like” genetic background (Figure 2B), with individuals of consistently mixed ancestry found in the site closest to the centre of the iridescence cline. Introgression from Panamanian populations could be detected in northern Colombian populations. The PCA supported this, with three clusters separated by geography along the first axis of variation, representing the Colombian populations, the Panamanian populations, and individuals with mixed ancestry and intermediate levels of iridescence clustered between them (Figure 2C). PC1 explained 5.84% of genetic variation in *H. erato*, with all subsequent eigenvectors explaining 0.7% or less of the genetic variation (SI Figure S4).

**Figure 2.**
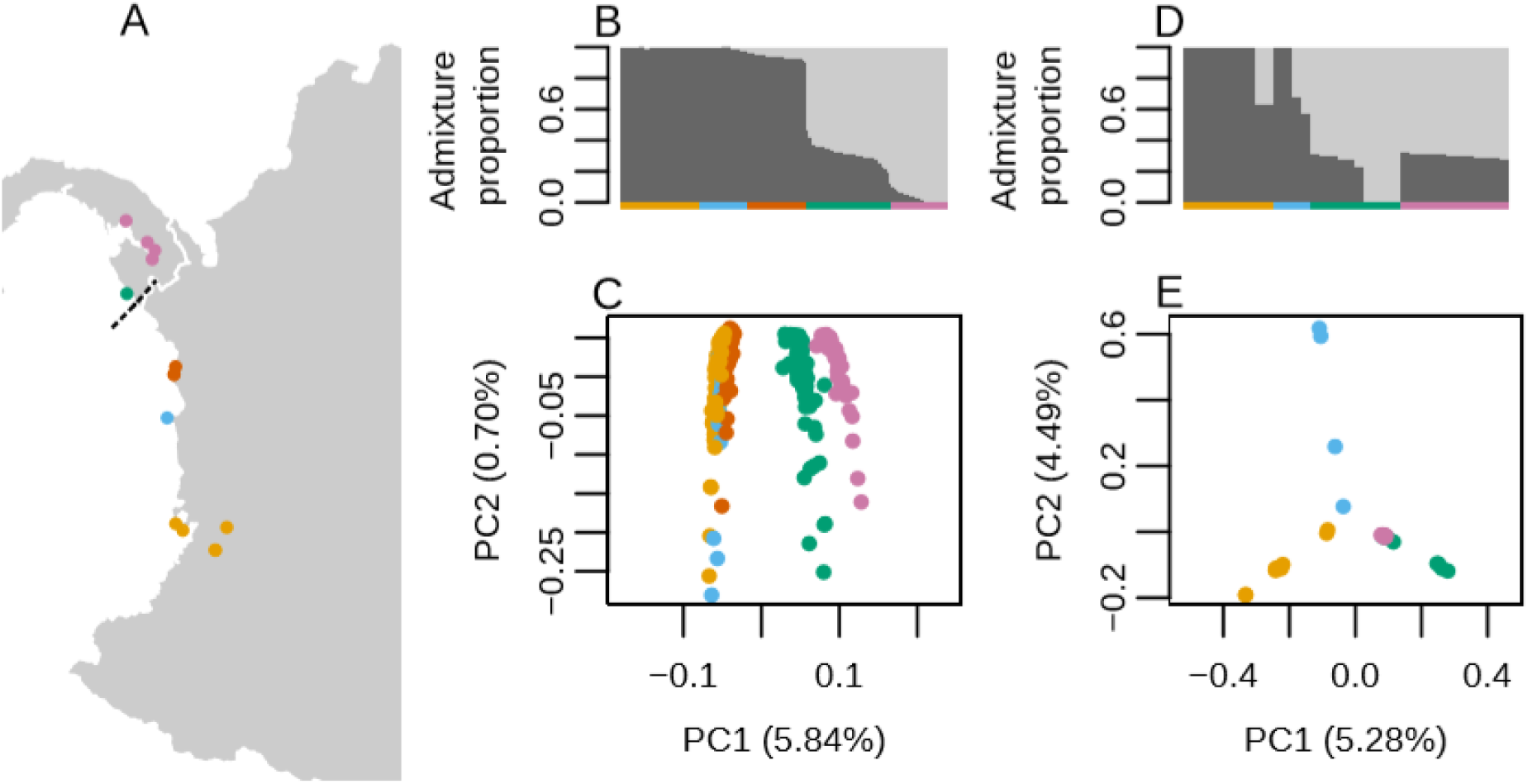
Population structure across the hybrid zones in *H. erato* (**B,C**) and in *H. melpomene* (**D, E**). Sampling locations across the hybrid zone. Approximate centre of the iridescence cline in *H. erato* indicated by a dashed line.(**A**). Individual admixture proportions estimated using NGSadmix, with *k* = 2 (**B, D**). Each vertical bar represents an individual, bar colour represents the estimated proportion of ancestry derived from population 1 (dark grey) or population 2 (light grey). Horizontal bars indicate the population of origin, colours match those on the map. Principal components analysis (**C, E**). Colour of points indicate the population of origin, as shown on the map.

NGSadmix also supported *K*=2 for *H. melpomene* (although *K*=1 cannot be tested, SI Figure S3D), but revealed a less straightforward population structure. While a “Colombia-like” genetic background could be seen, Panamanian individuals showed mixed ancestry, with the exception of four individuals from the site closest to the centre of the iridescence cline (Figure 2D). This is supported by the PCA. PC1 explained 5.28% of genetic variation, separating Colombian and Panamanian individuals. Individuals with intermediate levels of iridescence do not form an intermediate cluster between Panamanian and Colombian individuals, as is seen in *H. erato* (Figure 2E). The percent of genetic variation explained by PC1 and subsequent principal components show a more uniform distribution than in *H. erato* (SI Figure S4) consistent with weaker population structure.

Given the support for two genetic clusters, we compared the levels of differentiation between these populations using SNPs from individuals either side of the hybrid zone in southern Panama. Genome-wide average Hudson’s *F*_*ST*_ was estimated for each species, using the ratio of averages approach. This revealed that genome-wide divergence across the hybrid zone is greater in *H. erato* (*F*_*ST*_ =0.188), compared to *H. melpomene* (*F*_*ST*_=0.0739). The difference in genetic structure is also apparent in the plots of the pairwise genetic distance between sampling locations, plotted against their geographic distance. In *H. erato*, within-race comparisons that span distances of 195 – 325 km show a range of *F*_ST_ values between 0.063 – 0.129. However, between-race comparisons made over a similar range of distances (188 – 345 km) have substantially higher *F*_*ST*_ (0.226 – 0.271), suggesting that the genetic structure is much stronger than would be expected based on geography alone (Figure 3). The pattern in *H. melpomene* is very different, as the between-race comparisons span a similar range of *F*_*ST*_ values to the within-race comparisons.

**Figure 3.**
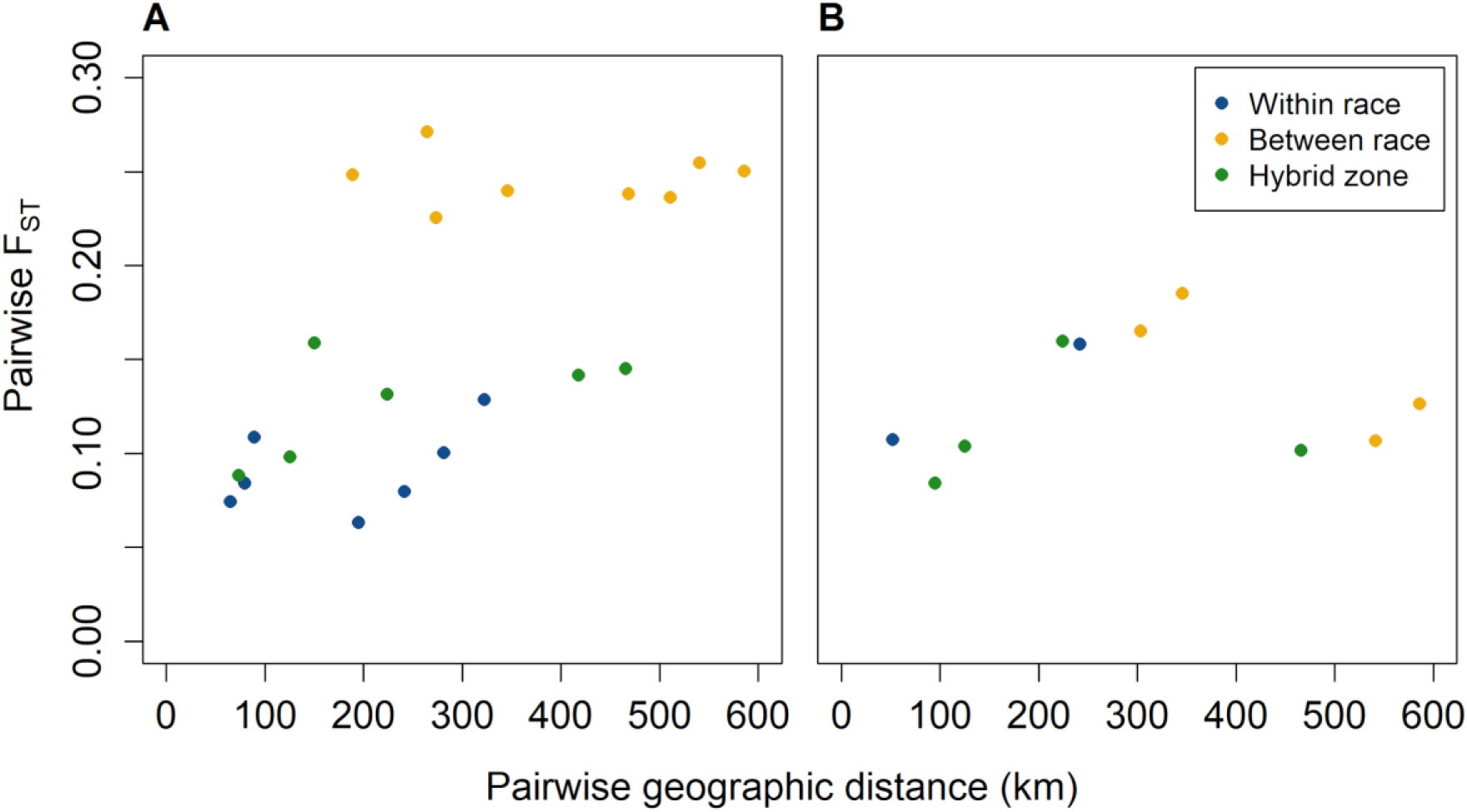
Relationship between geographic distance and genetic differentiation (genome-wide average *F*_*ST*_) between sampling sites in *H. erato* (**A**) and *H. melpomene* (**B**). Pairwise comparisons are colour-coded to indicate comparisons between populations of the same colour pattern race (blue), between populations of different colour pattern races (yellow), and comparisons where one population is from the hybrid zone centre (green).

### Phenotypic Variation

Strong phenotypic variation was observed across our range of sampling sites, with some difference apparent between *H. erato* and *H. melpomene* (SI Figure S2). The West Colombian yellow bar allele (y_wc_) was fixed in all Colombian sampling sites, apart from at some of the northernmost Colombian sampling sites near Bahía Solano (BS; SI Figure S2 C, D.). In *H. melpomene*, the frequency of y_wc_ gradually decreased, and persisted at comparable frequencies to the North Colombian yellow bar allele (Y) for ~200 km, before the Central American yellow bar allele (y_ca_) became predominant (SI Figure S2D). In contrast, in *H. erato* Y became the predominant allele, with y_ca_ approaching fixation towards the end of the transect (SI Figure S2C).

In both species the blue score, used as a proxy measure for iridescence, decreased across the transect (SI Figure S2 A,B). The colour measurements used to calculate the blue score were highly repeatable (p<0.001 for both red and blue values in both wing patches measured, Table S5). The bluest *H. melpomene* individuals were less blue than the bluest *H. erato* (SI Figure S2), which is consistent with reflectance spectrometry data from *H. erato* cyrbia and *H. melpomene* cythera (Parnell et al., 2018)

### Clinal variation within species

Cline fitting revealed that an asymmetrical stepped cline best described the variation in iridescence in *H. erato*, with a steeper right tail, which continually declines away from the cline centre. Neither stepped model was a significantly better fit than sigmoidal clines for the yellow bar in *H. erato*, and both colour traits in *H. melpomene* (Table 1; SI Table S6). For the admixture proportion, an asymmetrical stepped cline model was the best fit in *H. erato* (Table 1, SI Table S6), with a steeper right tail, similar to the iridescence cline, whereas the sigmoid model was the best fit for *H. melpomene*.

**Table 1.**
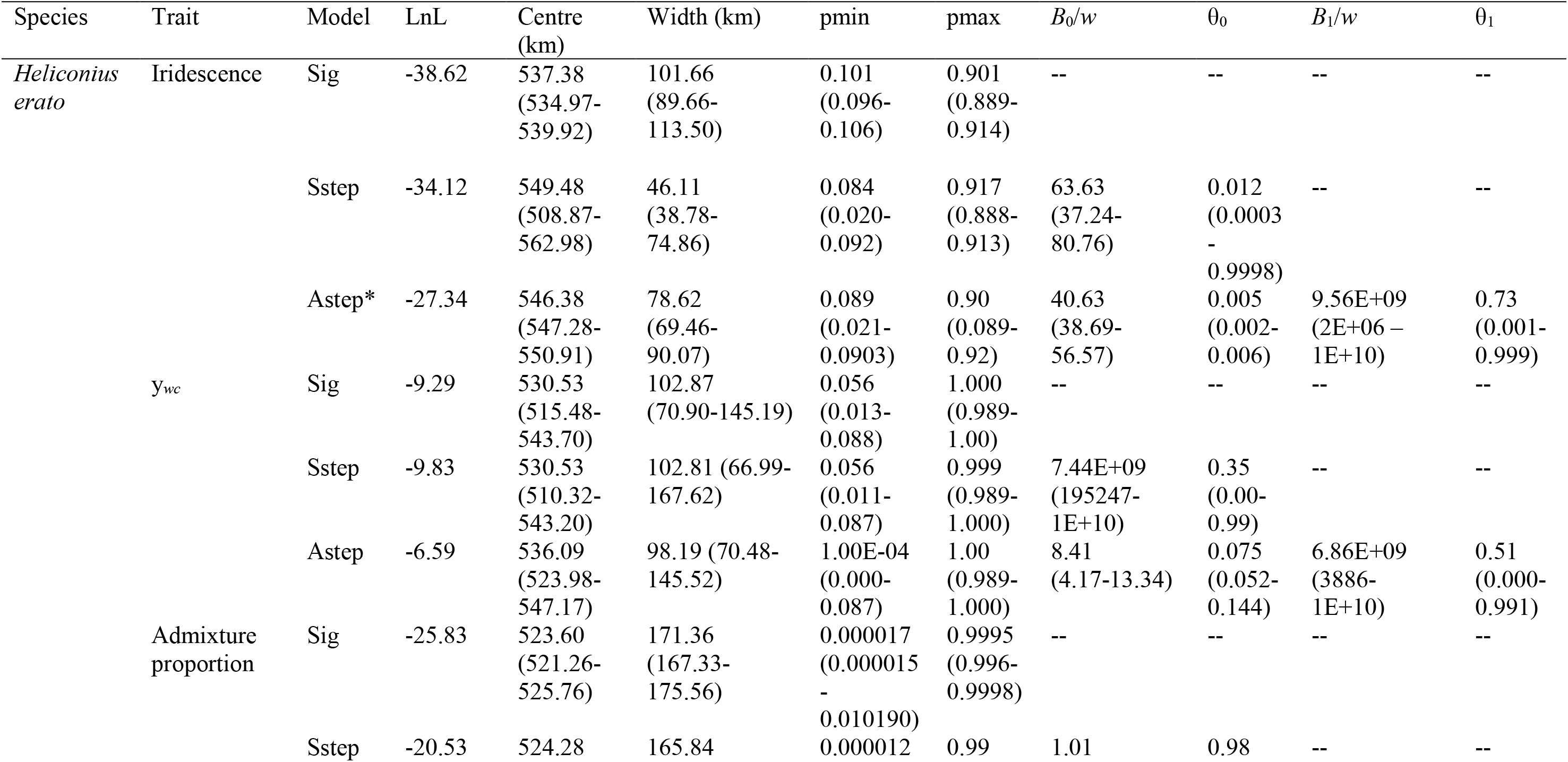
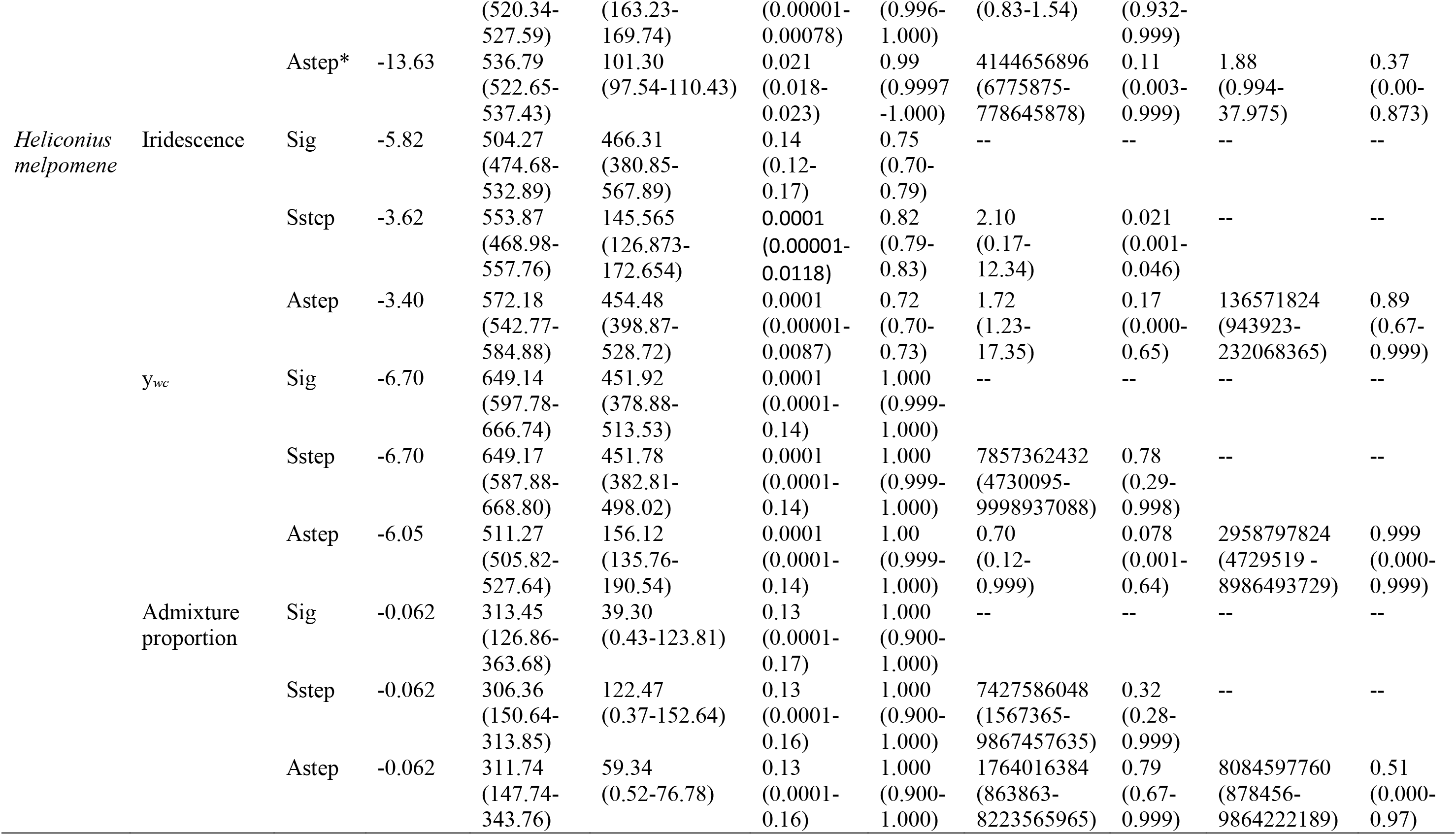
Cline parameter estimates for variation in iridescence, *y*_*wc*_ allele frequency, and admixture proportion across transects for *H. erato* and *H. melpomene*, which begin at the Queremal (Qu) locality. ML estimates for sigmoid models (Sig), symmetrical stepped models (Sstep), and asymmetrical stepped models (Astep) were estimated for each trait. If a model is a significantly better fit as determined by a likelihood ratio test (details in Table S6) it is denoted with *. Parameters are log-likelihood (LnL) cline centre (*c*), width (*w*), barrier strength for either side of stepped models (*B*_*0*_/*w*, *B*_*1*_/*w*), the rate of exponential decay for either tail (θ_0_, θ_1_).

Overall, our analysis indicates that the clines in iridescence, yellow bar and the admixture were highly similar within both of the species. Likelihood profiling revealed that we could not reject the null hypothesis that both iridescence and the yellow bar clines had coincident centres and concordant widths within both species. In *H. erato*, the parameters for the cline in the admixture score were different from the clines in iridescence (both centre and width) and yellow bar (centre only) (Table 2), though the differences were quite subtle. For *H. melpomene*, likelihood profiling indicated that neither the width nor centre of the admixture cline differed from the phenotypic clines; inspection of the likelihood surfaces indicates that power to reject the hypothesis of coincidence and concordance were low, as the profiles were flat across a broad range of the parameter space (Figures 3, S5). This could be due to weaker/non-clinal population structure, or the sparse sampling of genomic data within *H*. *melpomene*.

**Table 2.** Likelihood ratio tests for coincidence (*c*) and concordance (*w*) of iridescence, yellow bar, and admixture proportion clines. ∆ML is the test statistic, d.f. is degrees of freedom. The combination of clines being compared is noted under Trait(s).

| Trait(s) | *c* |  |  | *w* |  |  |
| --- | --- | --- | --- | --- | --- | --- |
| | ΔML | d.f. | *P* | ΔML | d.f. | *P* |
| *H. erato* |  |  |  |  |  |  |
| **iridescence, yellow bar** | 0.75 | 1 | 0.39 | 0 | 1 | 1.00 |
| **iridescence, admixture proportion** | 11.22 | 1 | < 0.001 | 16.85 | 1 | < 0.001 |
| **iridescence, yellow bar, admixture proportion** | 11.93 | 2 | 0.003 | 20.47 | 2 | < 0.001 |
| *H. melpomene* |  |  |  |  |  |  |
| **iridescence, yellow bar** | 0.80 | 1 | 0.37 | 0.004 | 1 | 0.95 |
| **iridescence, admixture proportion** | 0.22 | 1 | 0.64 | 0.008 | 1 | 0.93 |
| **iridescence, yellow bar, admixture proportion** | 1.02 | 2 | 0.60 | 0.01 | 2 | 0.99 |
| Both species |  |  |  |  |  |  |
| **iridescence** | 0.62 | 1 | 0.43 | 6.42 | 1 | 0.01 |
| **yellow bar** | 0.59 | 1 | 0.44 | 1.70 | 1 | 0.19 |
| **admixture proportion** | 0.22 | 1 | 0.64 | 0 | 1 | 1.00 |

The similarity of clines within the two species was supported by pairwise regression analysis, as the linear model was always a better description of the data than the quadratic polynomial (Figure S6). For *H. erato*, the linear model explained between 98% and 99% percent of the variation relationship between for all pairs of characters, and had a higher F-ratio than the polynomial quadratic, which explained a similar amount of the variation in the data (Table S7). The results were the same for *H. melpomene*, except that the quadratic fit was often a much poorer fit than the linear fit (Table S7).

### Comparison of clines between H. *melpomene* and *H. erato*

In contrast with the similar patterns of clinal variation within species, our profiling and regression analyses revealed striking differences in the cline shape between the species. For both iridescence, and yellow bar, the ML estimates of the cline width were roughly four times wider in *H*. *melpomene* than in *H. erato* (Table 1, Figure 4). For both traits, the peaks of the likelihood profiles did not overlap (Figure 5), with the difference being significant for iridescence (*p* = 0.01). Although the difference was not significant for yellow bar, because the change in likelihood was not as dramatic across the profile for that trait (*p* = 0.19), the regression analysis indicated that clines were not concordant as the quadratic model was a far better fit to the regression of the frequency of y_wc_ between *H*. *melpomene* and *H. erato* (Figure S6, Table S7). Both likelihood profile analysis and regression analysis indicated that the admixture clines were concordant (Figures 3, S6), but again had low power to detect any difference due to the coarse geographic sampling in both species.

**Figure 4.**
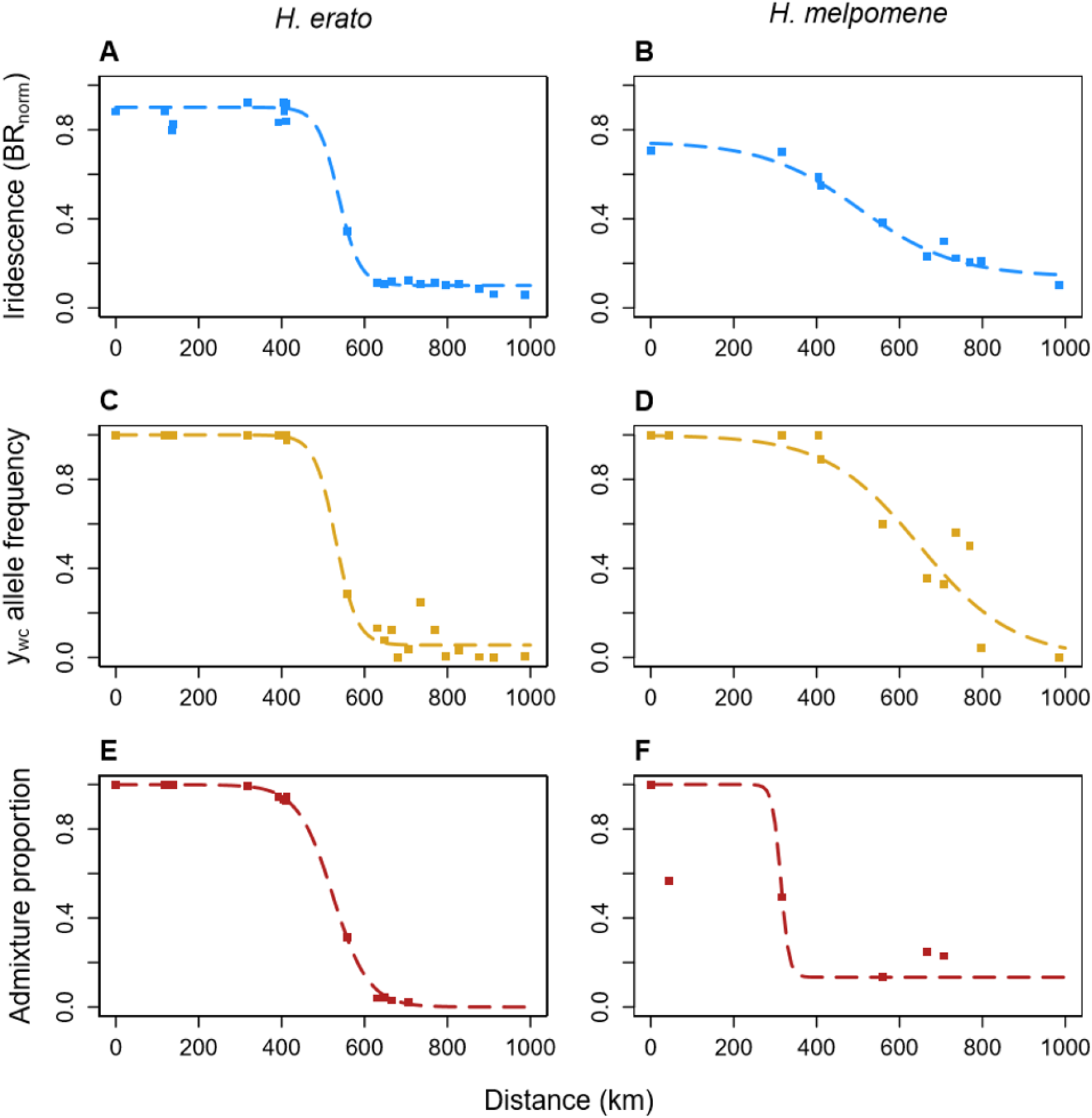
The best fitting geographic clines (dashed lines) of iridescence (**A, B**; blue), the West Colombian yellow bar allele frequency (y_wc_; **C, D**; yellow), and admixture proportions (**E, F**; red), across a transect of sampling sites (points) for *Heliconius melpomene* (**A, C, E**) and *Heliconius erato* (**B, D, F**). The transect begins (at 0 km) in the Queremal (Qu) locality, in the Cauca Valley region of Colombia.

**Figure 5.**
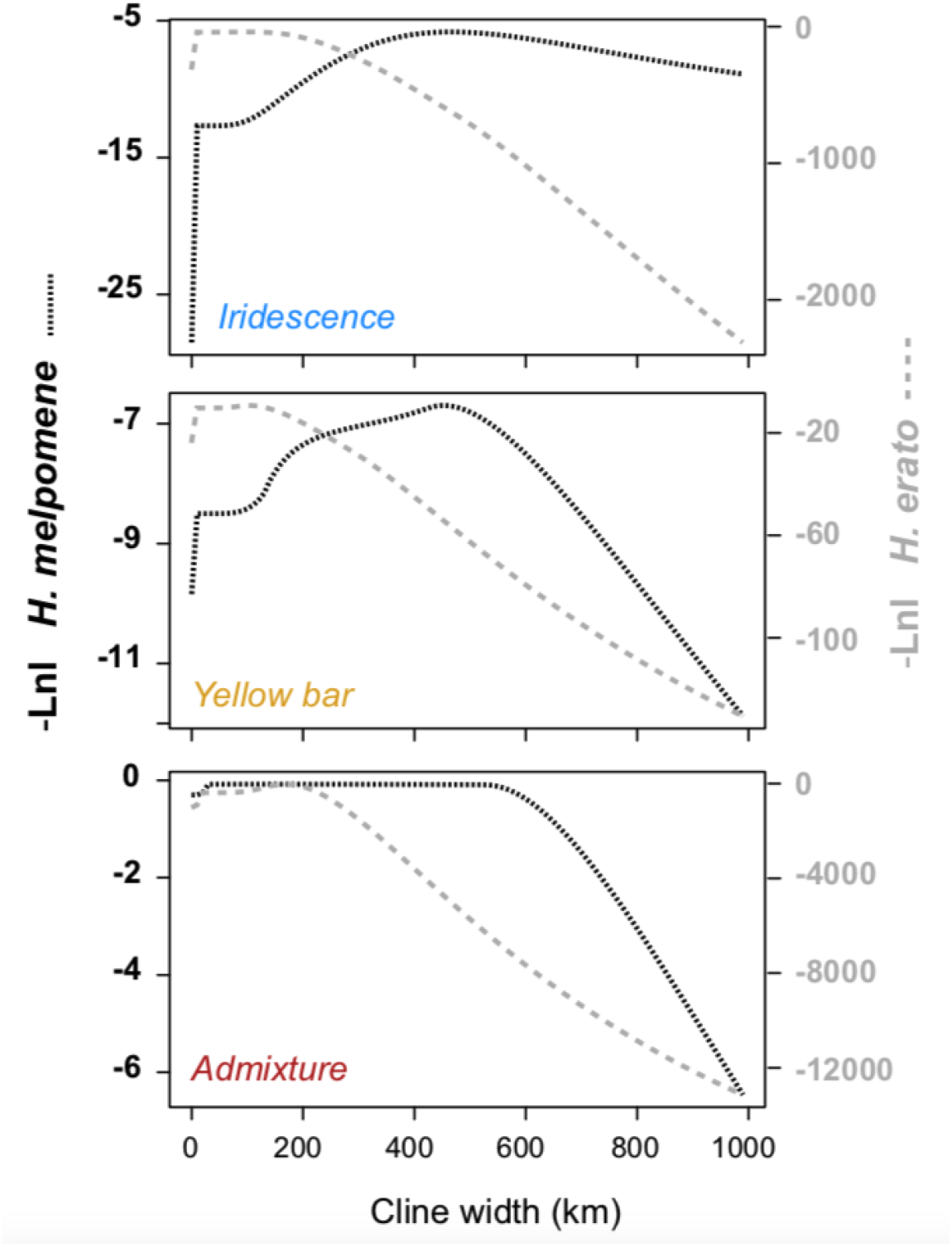
Likelihood profiles for the cline width for mean iridescence, frequency of yellow bar and mean admixture score for *H. melpomene* (narrow dashed line) and *H. erato* wide dashed line. Profiles were constructed using a step size of 10 km with all of the model parameters free to vary.

In contrast with the difference in cline widths, clines for each trait tended to have highly similar centres, indicating that they were positioned in roughly the same geographic area (Figure 4, Table 2). This is clearly observed in the likelihood profiles for each trait, as the -Lnl tended to peak over a relatively broad area, between 400 and 600 km along the transect (SI Figure S5). For all three traits the difference in the location of the peak likelihoods was not significantly difference (*P* ranging from 0.22 - 0.62)

Given the differences in cline width between the species, we estimated the effective selection (*s*\*) on y_wc_ across the hybrid zone in both species using the ML estimates and support limits of cline width (Table 1) and the dispersal estimates from Mallet et al. (1990) of 2.6 km for *H. erato* and 3.7 km for *H. melpomene*. Selection estimates were 0.00203 (0.00102–0.00427) for *H. erato*, and 0.000213 (0.000165– 0.000303) for *H. melpomene*. Blum (2002) estimates higher dispersal for *H. erato*, 10 km, which increased the value of *s*\* to 0.0300 (0.0151–0.0632). Given that the widths of the yellow bar and iridescence clines were not different within each species, similar estimates were found for iridescence. For *H. melpomene*, *s*\*=0.000200 (0.000135–0.000300), for *H. erato*, *s*\*= 0.00208 (0.00167–0.00267) if the dispersal distance is 2.6 km, and 0.0307 (0.0247–0.0395) if the dispersal distance is 10 km. However, it should be noted that in the case of iridescence, *s*\* is the average strength of selection acting across loci controlling iridescence.

## Discussion

Our analysis of parallel hybrid zones in the co-mimics *H. erato* and *H. melpomene* has revealed similarities, as well as striking differences in colour trait divergence between the species. Consistent with the predictions of the mimicry hypothesis, the clines in yellow bar and iridescence are highly coincident within and between species, suggesting that they are maintained by the same selective pressure. In contrast, the width of the clines in both colour traits vary substantially between the species, being far wider in *H. melpomene*. The difference in cline widths is probably due, at least in part, to differences in the strength of direct selection acting on colour variation between *H. melpomene*, and *H. erato*. However, differences in population structure and levels of genomic differentiation indicate that species-specific factors, such as different population histories, dispersal rates, and strength of reproductive isolation between the subspecies may also contribute to the different cline widths between the species.

### Comparing clines within species

Our geographic cline analysis revealed that clines in mean iridescence and the y_wc_ allele frequency had highly similar centres and widths within both of the species. While this is predicted to result from direct selection on a warning colour pattern, similar clines could also arise as a correlated response to selection if traits have a shared genetic basis, or if the loci that underlie them are physically linked (Price & Langen, 1992). We are able to rule out these explanations for the similarity of the clines in *H. erato*, as these colour traits segregate independently in F_2_ crosses made between iridescent and non-iridescent races (Brien et al., 2018). As the colour pattern traits studied here very different genetic architectures, with the yellow bar being controlled by a single major-effect locus (Joron et al., 2006; Mallet, 1986; Nadeau et al., 2016), and iridescence being controlled by multiple genes (Brien et al., 2018), it is highly unlikely that the clines in one of the colour traits could arise as a correlated response to selection acting on the other.

Another alternative explanation for the similar clines within the species, aside from direct selection acting on each trait, is that the clines are maintained by a permeable, but genome-wide barrier to gene flow between the subspecies. When reproductive isolation involves a large number of loci, or is very strong, selection against unfit hybrid offspring can generate a barrier to gene flow that can impact the spread of even neutral alleles across a hybrid zone (Barton & Gale, 1993). For a trait that is also under direct selection, the importance of the overall barrier in shaping a cline depends on the strength of direct selection acting on the trait relative to the strength of indirect selection resulting from selection at other barrier loci. For example, if the strength of indirect selection acting on a trait is much greater than direct selection, then the cline shape will be more informative about the overall barrier strength, and tell us nothing about the strength of direct selection. In situations where this is the case, clines in the trait should show a “stepped”, rather than sigmoid shape. In *H. erato*, the two colour pattern clines are coincident, and the best fitting cline model for variation in iridescence is stepped, which would indicate that indirect selection plays some role in shaping the cline (Kruuk et al., 1999). In contrast, the simple sigmoidal cline fits best for the yellow bar. Finally, while we do see stepped clines in *H. erato*, they are asymmetrical, with a left tail closely resembling that of the sigmoidal cline, and a much steeper right tail. It is possible that these tails reflect genuine asymmetry, due to hybrid zone movement, which has been predicted and documented in these species (Blum, 2002; Mallet, 1986; Thurman, Szejner‐Sigal, & McMillan, 2019). Although it is difficult to determine the overall importance and indirect selection in shaping these clines, it is unlikely that all of our results can be explained purely in terms of indirect selection.

Given the abundant evidence for the role of direct selection in shaping colour pattern variation across the genus *Heliconius*, it is likely that it plays at least some role in shaping variation in these species. Under a scenario where the colour pattern clines are maintained by a balance between migration and divergent ecological selection, similar cline centres arise when both traits experience the same source of selection, or when different ecological gradients change in approximately the same location (Barton & Hewitt, 1985). In *Heliconius*, local warning colour patterns are maintained by predator-mediated positive frequency-dependent selection, with rare colour morphs experiencing increased predation (Benson, 1972; Dell’aglio, Stevens, & Jiggins, 2016; Langham, 2004; Mallet & Barton, 1989). The centre of colour pattern clines could represent the location where the most effective warning pattern shifts to that of a neighbouring subspecies. The coincidence of cline centres for iridescence and y_wc_, which is observed in both *H. melpomene* and *H. erato*, suggests that both traits contribute to the warning signal.

In *H. melpomene*, the width and centre of the admixture proportion cline was not significantly different to the colour pattern clines. However, variation in admixture proportions had a poor fit to any of the cline models (Figures 2, 3, S5), illustrated by the large confidence intervals (Table 1). This is in part due to coarse sampling, but can also be explained by the less defined population structure in this species. Clear phenotypic intermediates in the hybrid zone are not of mixed ancestry (Figures 3, 4). This suggests that divergence in iridescence in this species is not tightly coupled with genome-wide differentiation. The broad phenotypic clines that we see are not characteristic of the steep, stepped clines which result from strong LD between selected loci and indirect selection (e.g. Szymura & Barton, 1991), and are more likely due to weak selection and/or isolation-by-distance.

### Comparing clines between species

Although patterns of clinal variation are very similar within species, we observed substantial differences between *H. melpomene* and *H. erato*. This is in contrast to what is expected, based on the strong existing evidence that colour pattern convergence in this pair of co-mimics is driven by Müllerian mimicry - a common positive frequency-dependent selection pressure based on predator learning. The coincidence of cline centres in colour pattern traits between the co-mimics is consistent with this hypothesis, as it suggests that variation in both species is structured by the same agent of selection. While the clines in yellow bar and iridescence are coincident between species, they are four times wider in *H. melpomene*. This difference could result from (*i*) variation in the strength of direct or indirect selection between the species, (*ii*) species-specific differences in the dispersal rate, (*iii*) different demographic histories between the species, or a combination of these explanations.

First, the wider clines in *H. melpomene* could be a result of having a greater dispersal capability. Direct estimates of dispersal are difficult in *Heliconius* butterflies due to most dispersal occurring soon after adult eclosion (Mallet, 1986a). The most reliable estimates are thought to be those made using cline theory and patterns of linkage disequilibrium (Blum, 2002; Mallet et al., 1990), including the only direct comparison of *H. erato* and *H. melpomene* (Mallet et al., 1990). This study reports higher dispersal distances in *H. melpomene*. However, our estimates of the selection coefficient *s*\* (Barton & Gale, 1993) show that even if this higher dispersal rate is taken into account, colour pattern traits in *H. melpomene* appear to be under much weaker selection. Other studies on parallel hybrid zones between neighbouring races in this species pair show that *H. melpomene* tend to have wider clines than *H. erato*, but not to the degree seen in the present study (Mallet et al., 1990; Salazar, 2012). *H. melpomene* displays less vivid iridescence than its co-mimic, and the colour difference between iridescent and non-iridescent *H. melpomene* is less pronounced than the colour difference between *H. erato* races (Parnell et al., 2018, Figure 1). Hybrid phenotypes are therefore less distinct from the parental populations in *H. melpomene* which could result in weaker selection against hybrid offspring.

The difference in divergence of one of the colour traits, namely iridescence, could also explain why clines differ in shape between the co-mimics. The main predators of *Heliconius* butterflies are thought to be birds of the tyrant flycatcher (Tyrannidae) and jacamar (Galbulidae) families (Jiggins, 2017), hence bird predation is expected to be the main driver of mimicry and phenotypic convergence between species. Previous work modelling bird visual systems has shown that birds can discriminate between the iridescent blue in *H. erato* and *H. melpomene* (Parnell et al., 2018). However, iridescence in *H. melpomene* is not as bright as in *H. erato*, which means that the visibility of the trait to selection may also vary between the species. This may weaken the overall mimetic signal in *H. melpomene*, which may also influence the strength of selection acting on the yellow bar.

Another possible factor that may explain the differences in cline widths between the species is that they may have experienced very different demographic histories. The inclusion of genomic data in our study revealed a striking difference in the level of population structure across these parallel hybrid zones. Specifically, we found strong divergence across the *H. erato* hybrid zone in contrast with the very weak structure across the *H. melpomene* hybrid zone. The defined population structure in *H. erato* is typically associated with populations that have diverged in allopatry, followed by secondary contact. This scenario can lead to genetic discontinuity and coincidence of clines in multiple traits (Barton, 1983), along with strong genome-wide reproductive isolation, meaning that indirect selection can play a greater role in the maintenance of geographic clines. It is also possible that strong selection acting on a quantitative trait in *H. erato* could be responsible for the formation of a genome-wide barrier to gene flow (Feder et al., 2012), although conservative estimates suggest iridescence is not polygenic enough to act as such a barrier (Brien et al., 2018). In contrast, the relatively low genetic structure in *H. melpomene* is usually associated with a primary intergradation scenario, where hybrid zones form due to divergent selection acting across a strong environmental gradient. Because primary hybrid zones form in the face of continuous gene flow, other barriers cannot evolve in isolation, meaning that direct selection on phenotypic traits must alone overcome migration. This makes it much harder for sharp clines to become established. Future studies of the historical demography of these species may shed more light on the role of history in shaping phenotypic traits associated with mimicry.

## Conclusions

Examples of parallel evolution are celebrated as some of the best evidence for the power of natural selection in driving local adaptation. Most studies of parallel evolution tend to focus on understanding the similarities between populations and species subject to the same selective pressures. Despite the striking parallelism at the level of the phenotype, including simple and complex colour traits, the Müllerian co-mimics *H. erato* and *H. melpomene* show striking differences in how trait variation is structured across geography, which would not be apparent without detailed sampling and analysis across their distribution. Although mimicry has almost certainly been the primary driver of parallel evolution in this system, other factors are needed to explain patterns of phenotypic variation, both within and between species. More focus on phenotypic differences may provide new insight into the processes underlying parallel evolution, and may help us to understand the factors that limit adaptation in general.

## Acknowledgements

Thanks to the governments of Panama and Colombia (ANLA-Permit 0530) for giving permission to collect butterfly specimens. Thanks to the McMillan and Jiggins labs for providing access to samples. Thanks also to Patricio Salazar, Juan Enciso, Juan Camilo Dumar, Melanie Brien, Carlos Arias, Agata Surma and others in Panama and Colombia, in particular the residents of Jaqué, Darién, for help with logistics and collecting in the field. Thanks to Roger Butlin for valuable comments on this manuscript. This work was funded by the UK Natural Environment Research Council (NERC) through an Independent Research Fellowship (NE/K008498/1) to NJN, and by The Royal Society through an International Exchange Scheme grant. EVC was funded by the NERC doctoral training partnership, ACCE. CS and CP were funded by COLCIENCIAS (Grant FP44842-5-2017).

## Data Accessibility Statement

Sequence data have been deposited in the European Nucleotide Archive with the project number PREJEB32848. Code for implementing the regression-based tests for concordance can be found at https://github.com/seanstankowski/Heliconius_MS. Phenotype measurements deposited at Dryad: XXX

## Author Contributions

EVC and NJN conceived and designed the study. EVC, NJN, CPD, CAS and ML carried out field work. EVC generated and analysed the data with the help of SS. EVC wrote the manuscript, and all co-authors revised the manuscript.

