## Supplementary Information for "Müllerian mimicry of a quantitative trait despite contrasting levels of genomic divergence and selection"

### Parallel clines in a quantitative trait in butterfly co-mimics despite different levels of genomic divergence and selection

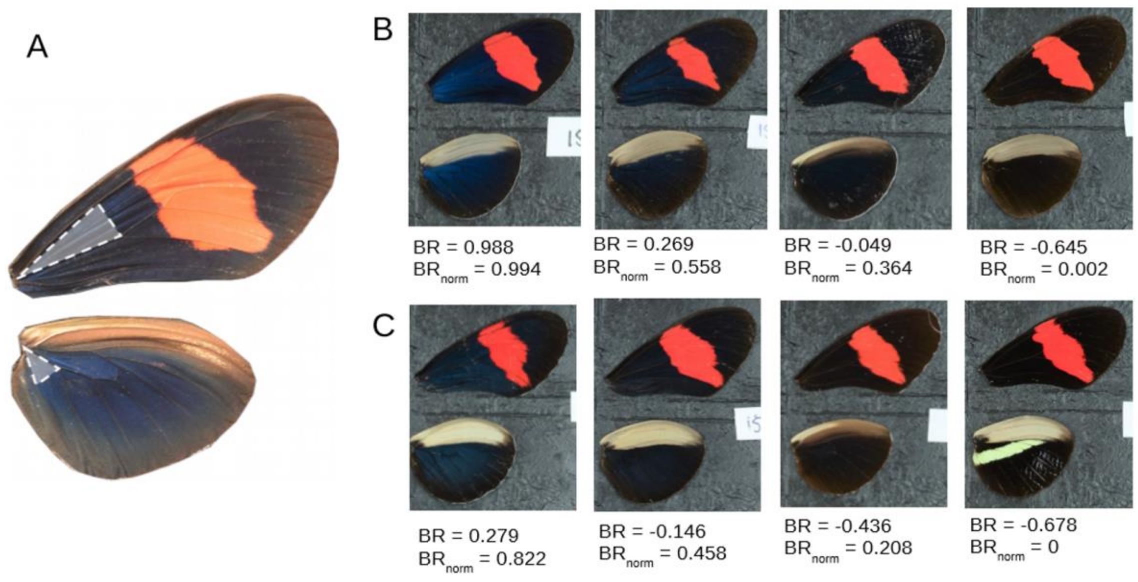


**Figure S1** – A) The wing regions highlighted were used to gather mean RGB scores for each individual. B) A selection of *Heliconius erato* wings with a range of BR (blue-red ratio) and $BR_{norm}$ (normalised blue-red ratio) values, spanning the range of iridescence. C) A selection of *H. melpomene* wings with a range of BR and $BR_{norm}$ values.

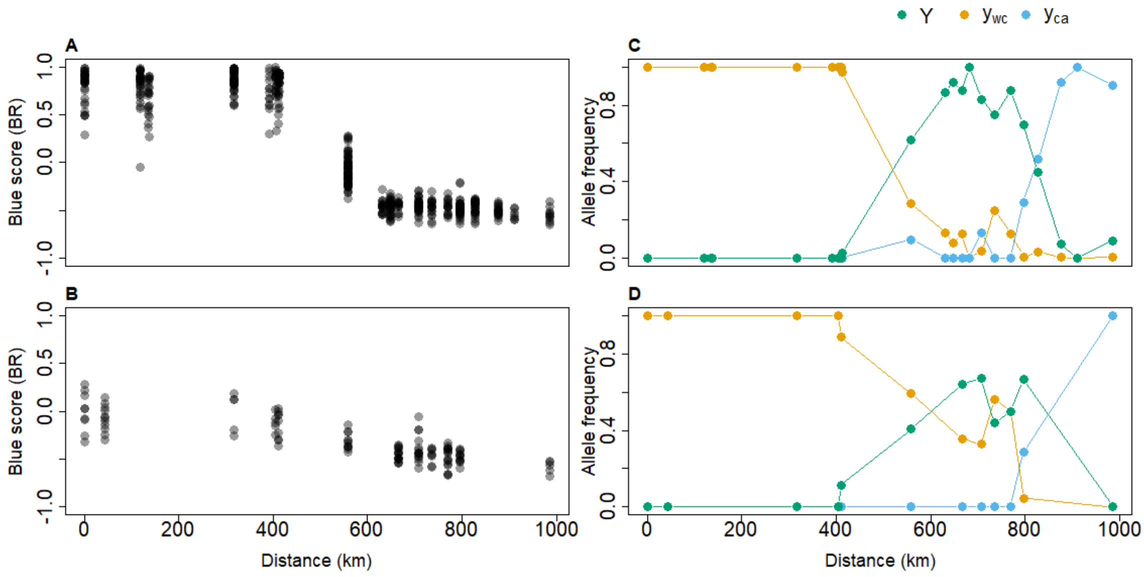


**Figure S2.** Variation in wing pattern phenotypes across the transect, from 0 km at Queremal (Qu), Colombia, to 987.85 km at El Valle, Panama. Blue score (BR) values of each individual for *H. erato* (**A**) and *H. melpomene* (**B**). Frequency of the North Colombian (Y, green), West Colombian ($y_{wc,}$ orange) and Central American ($y_{ca,}$ blue) yellow bar allele at each site with 5 or more samples for *H. erato* (**C**) and *H. melpomene* (**D**).

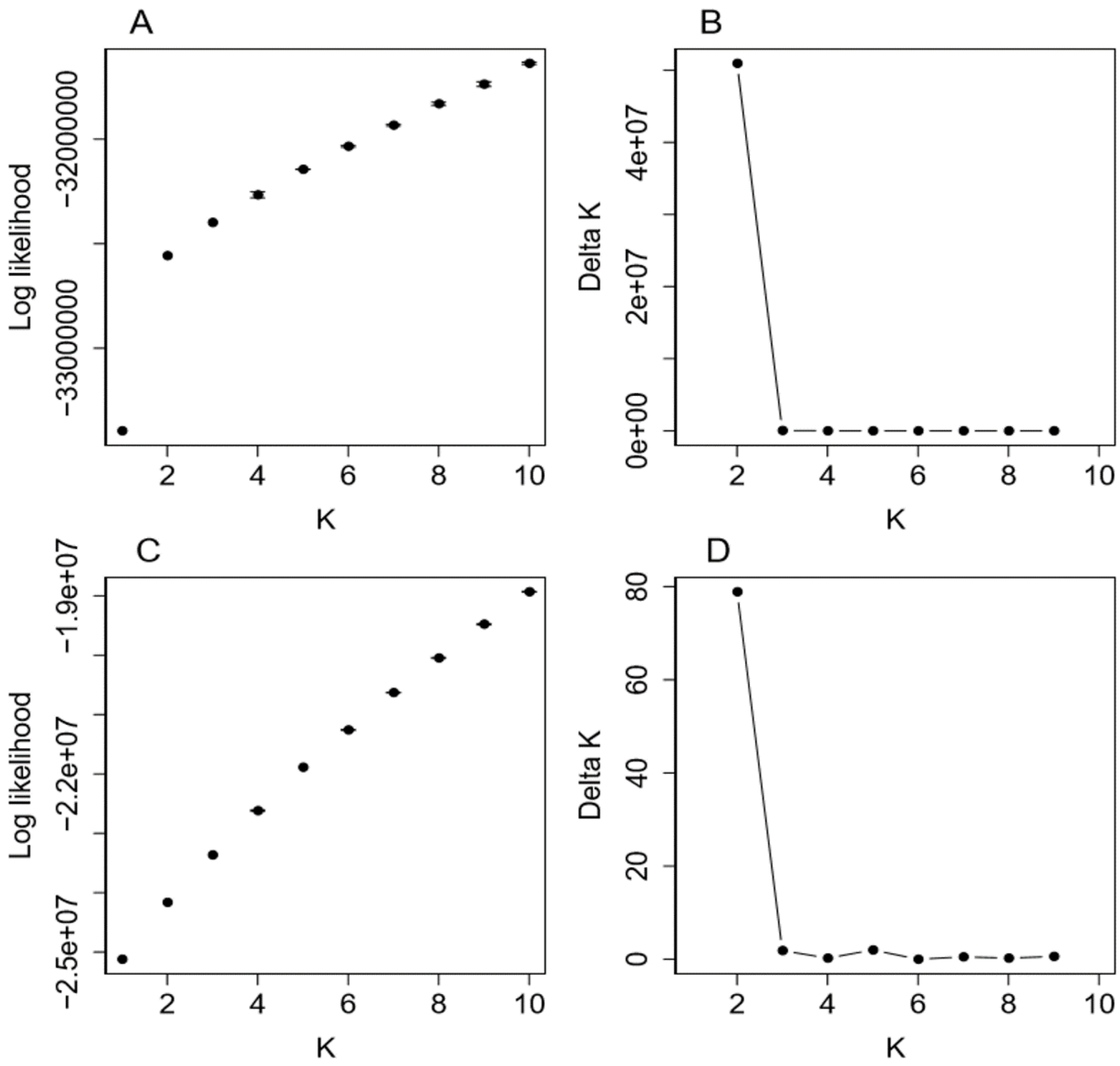


**Figure S3** – The log likelihood scores for increasing values of *k*, as estimated in the software NGSadmix for (A) *H. erato*, and (C) *H. melpomene.* The delta (Δ) *k* score is also given for (B) *H. erato* and (D) *H. melpomene.*

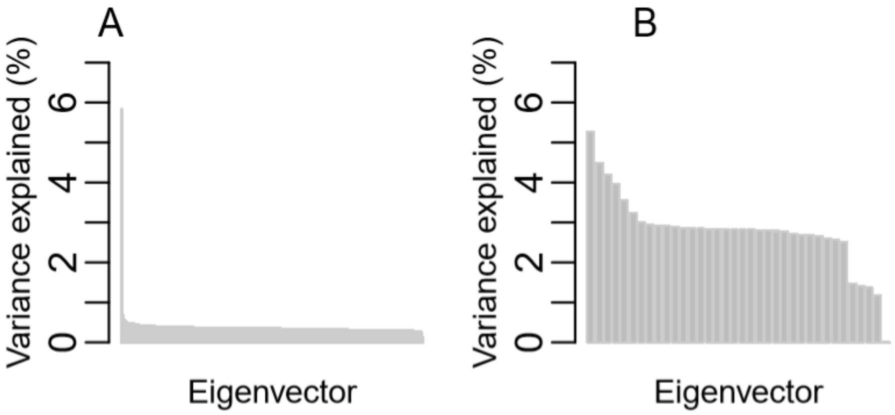


**Figure S4 –** Percent of genetic variance explained by each eigenvector from the principal components analysis for (A) *H. erato* and (B) *H. melpomene*

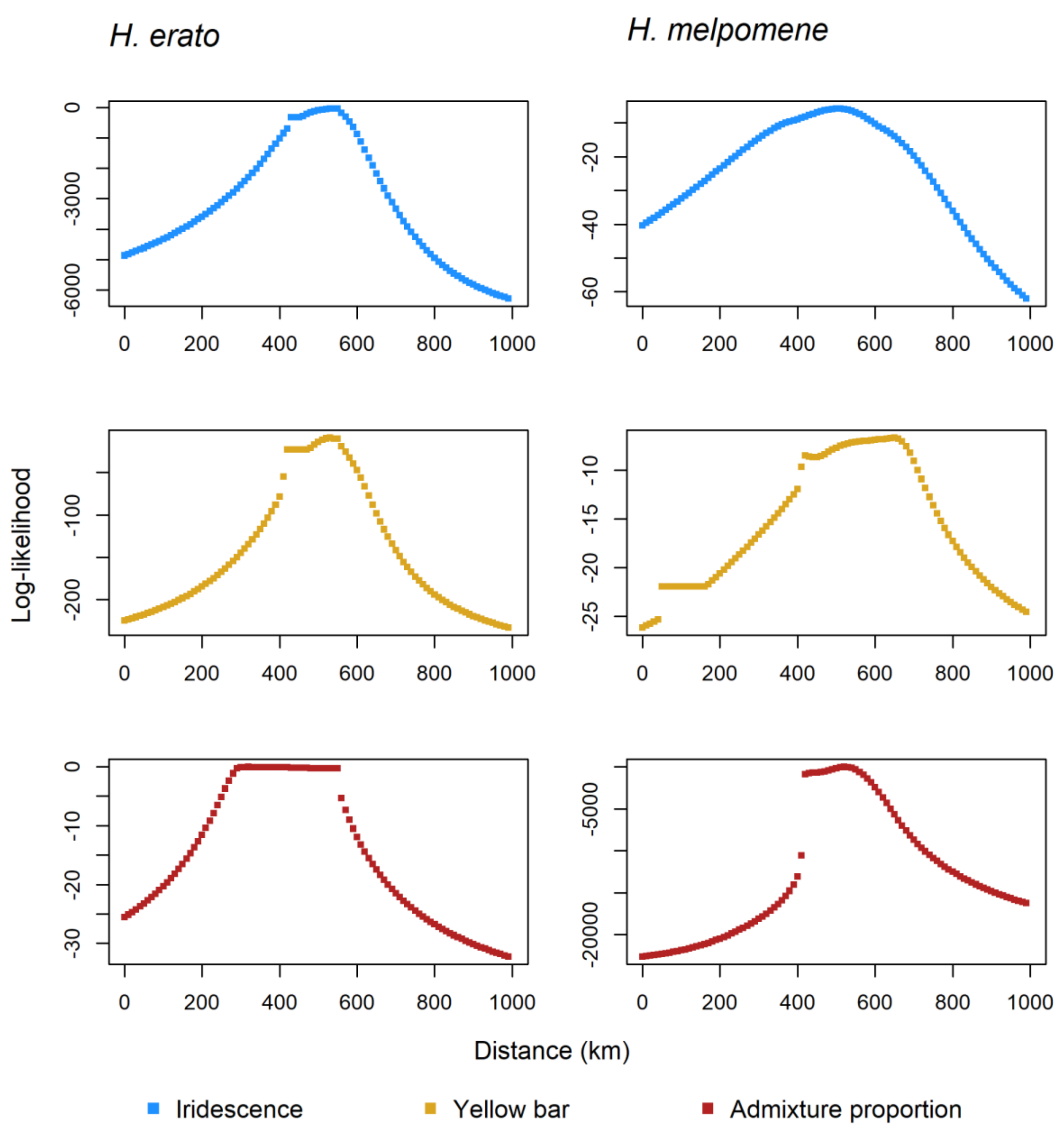


**Figure S5** – Log likelihoods for a range of values (0 and 1000 km at increments of 10 km) of cline centre for iridescence (blue), $y_{wc}$ (yellow), and admixture scores (red), for *H. erato* (left) and *H. melpomene* (right).

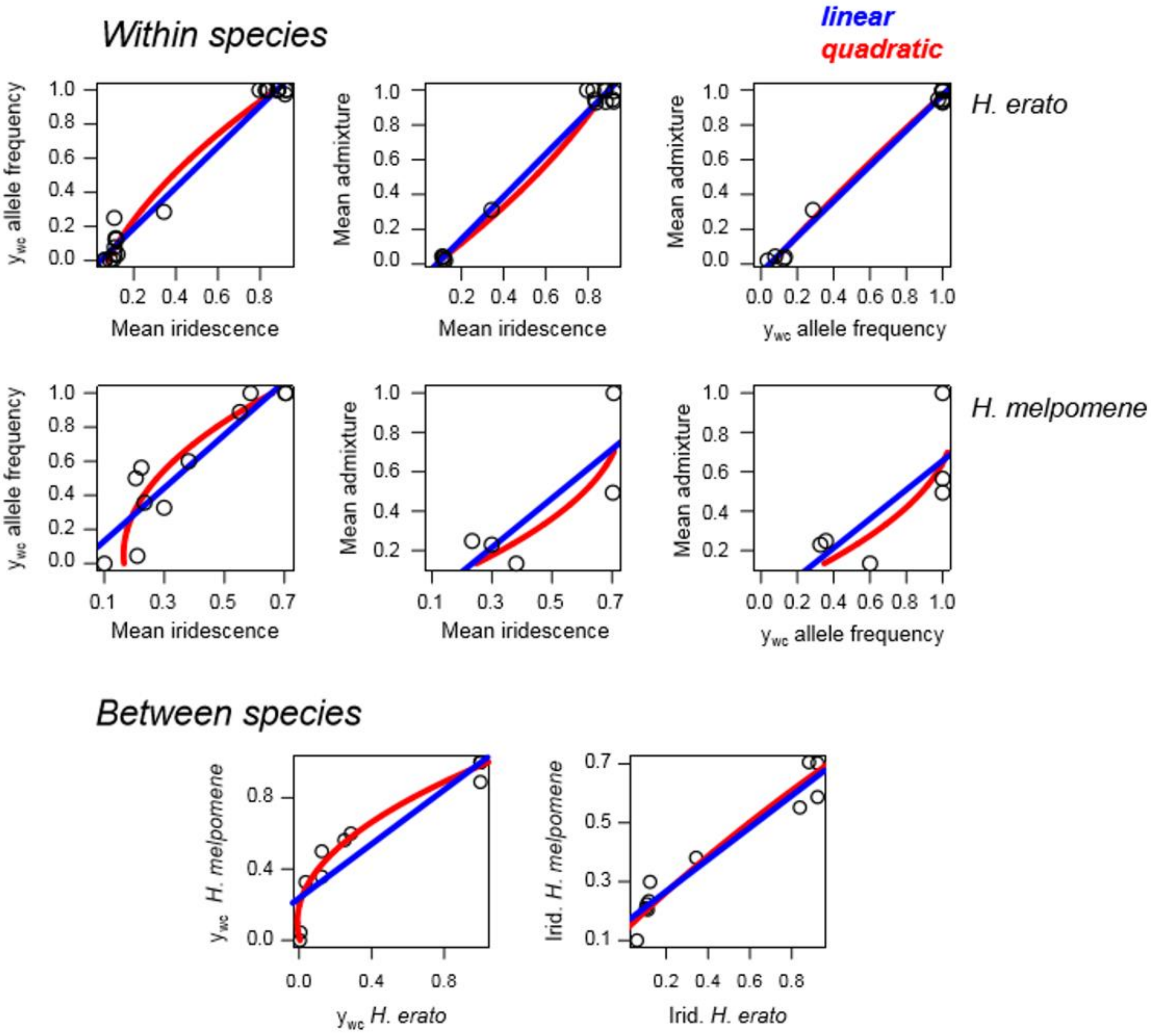


**Figure S6 -** Plots of the linear and quadratic polynomial models fit to pairs of traits within species and each trait between species. Note that this analysis could not be used to compare the admixture clines between the species because many sample locations only included genetic data for one of the species.

**Table S1** – Sampling locations and number of *H. erato* and *H. melpomene* individuals phenotyped (and genotyped)

| Locality (Site, Province, Country) | | Lat., Lon. | *Heliconius erato* | *Heliconius melpomene* | Collection | Year collected |
|---|---|---|---|---|---|---|
| (Qu) | El Queremal, Valle del Cauca, Colombia | 3.53199, -76.75461 | 43(24) | 9(8) | Sheffield (Nadeau) | 2015 |
| (RB) | Río Bravo, Valle del Cauca, Colombia | 3.88274, -76.57716 | 0 | 11 (2) | Sheffield (Nadeau) | 2015 |
| (SP) | San Pedro, Valle del Cauca, Colombia | 3.83691, -77.25734 | 41(24) | 0 | Sheffield (Nadeau) | 2015 |
| (La) | Ladrilleros, Valle del Cauca, Colombia | 3.93944, -77.36889 | 14(15) | 3 | Rosario (Salazar) | 2003, 2004 2007, 2008 2010, 2012 |
| (Ba) | La Barra, Valle del Cauca | 3.958333, -77.37333 | 9 | 0 | Rosario (Salazar) | 2003, 2004 2007, 2008 2010, 2012 |
| (Am) | Amargal, Chocó, Colombia | 5.57187, -77.50211 | 68(39) | 5(4) | Sheffield (Nadeau) | 2015 |
| (PP) | Playa Parra, Chocó, Colombia | 6.24066, -77.39544 | 18(6) | 0 | Sheffield (Nadeau) | 2016 |

| | | | | | | |
|---|---|---|---|---|---|---|
| (BS) | Bahía Solano, Chocó, Colombia | 6.38765, -77.38557 | 73(42) | 17 | Sheffield (Nadeau) | 2016 |
| (Ja) | Jaqué, Daríen, Panama | 7.48767, -78.12733 | 110(68) | 11(10) | Sheffield (Nadeau) | 2015 |
| (RF) | Rancho Frío, Daríen, Panama | 8.01974, -77.73252 | 19(16) | 1 | Cambridge (Jiggins) | 2011 |
| (Ya) | Yaviza, Daríen, Panama | 8.15602, -77.69308 | 40(4) | 1(1) | Cambridge (Jiggins) | 2011 |
| (SL) | Santa Librada, Daríen, Panama | 8.27971, -77.80982 | 19(11) | 23(3) | Cambridge (Jiggins) | 2011 |
| (LB) | Puerto Lajas Blancas, Daríen, Panama | 8.39141, -77.84473 | 1 | 2 | Cambridge (Jiggins) | 2011 |
| (PL) | Puerto Lara, Daríen, Panama | 8.61355, -78.13982 | 58(18) | 28(7) | Cambridge (Jiggins) | 2011 |
| (AF) | Agua Fría, Daríen, Panama | 8.85887, -78.22508 | 18 | 8 | STRI (McMillan) | 2015 |
| (Ip) | Ipeti, Panama, Panama | 8.97292, -78.51060 | 16 | 17 | STRI (McMillan) | 2015 |

| | | | | | | |
|---|---|---|---|---|---|---|
| (Ma) | Mangowichi, Panama, Panama | 9.13767, -78.68952 | 35 | 12 | STRI (McMillan) | 2015 |
| (El) | El llano, Panama, Panama | 9.23753, -78.95347 | 30 | 1 | STRI (McMillan) | 2015 |
| (To) | Tocumen, Panama, Panama | 9.20042, -79.39525 | 20 | 3 | STRI (McMillan) | 2015 |
| (Ga) | Gamboa, Colón, Panama | 9.11605, -79.69837 | 7 | 0 | STRI (McMillan) | 2015 |
| (EV) | El Valle, Coclé, Panama | 8.59323, -80.14108 | 11 | 5 | STRI (McMillan) | 2015 |

**Table S2.** Sequencing coverage, phenotype, and location information per individual for RAD sequenced *Heliconius erato.* Sequencing coverage information was calculated after duplicates were removed, and before quality filtering. The bar category refers to the yellow hindwing bar category, defined by Mallet (1986). BR refers to the blue score, a ratio of the blue-red content of photographed wings, described in the Methods.

| ID | species | race | Country | Province | Locality | Lat | Lon | Depth of coverage | % bases sequenced | bar category | BR | Accession (ENA) |
|---|---|---|---|---|---|---|---|---|---|---|---|---|
| 3127 | *H. erato* | *venus* | Colombia | Valle del Cauca | Queremal | 3.53199 | -76.75461 | 3.51 | 8.3 | C | 0.85 | ERS3546500 |
| 3131 | *H. erato* | *venus* | Colombia | Valle del Cauca | Queremal | 3.53199 | -76.75461 | 6.06 | 11.94 | C | 0.87 | ERS3546501 |
| 3355 | *H. erato* | *venus* | Colombia | Valle del Cauca | Queremal | 3.53199 | -76.75461 | 7.43 | 13 | C | 0.96 | ERS3546502 |
| 3358 | *H. erato* | *venus* | Colombia | Valle del Cauca | Queremal | 3.53199 | -76.75461 | 3.71 | 8.93 | C | 0.63 | ERS3546503 |
| 15N100 | *H. erato* | *venus* | Colombia | Valle del Cauca | Queremal | 3.53199 | -76.75461 | 4.14 | 9.66 | C | 0.60 | ERS3546504 |
| 15N103 | *H. erato* | *venus* | Colombia | Valle del Cauca | Queremal | 3.53199 | -76.75461 | 8.81 | 14.49 | C | 0.94 | ERS3546505 |
| 15N105 | *H. erato* | *venus* | Colombia | Valle del Cauca | Queremal | 3.53199 | -76.75461 | 6.33 | 12.82 | C | 0.99 | ERS3546506 |

| | | | | | | | | | | | | |
|---|---|---|---|---|---|---|---|---|---|---|---|---|
| 15N106 | *H. erato* | *venus* | Colombia | Valle del Cauca | Queremal | 3.53199 | -76.75461 | 5.19 | 11.59 | NA | NA | ERS3546507 |
| 15N107 | *H. erato* | *venus* | Colombia | Valle del Cauca | Queremal | 3.53199 | -76.75461 | 6.19 | 11.6 | C | 0.76 | ERS3546508 |
| 15N108 | *H. erato* | *venus* | Colombia | Valle del Cauca | Queremal | 3.53199 | -76.75461 | 5.51 | 11.91 | C | 0.92 | ERS3546509 |
| 15N109 | *H. erato* | *venus* | Colombia | Valle del Cauca | Queremal | 3.53199 | -76.75461 | 7.07 | 13.11 | C | 0.84 | ERS3546510 |
| 15N114 | *H. erato* | *venus* | Colombia | Valle del Cauca | Queremal | 3.53199 | -76.75461 | 5.56 | 11.95 | C | 0.85 | ERS3546511 |
| 15N122 | *H. erato* | *venus* | Colombia | Valle del Cauca | Queremal | 3.53199 | -76.75461 | 7.17 | 13.26 | C | 0.90 | ERS3546512 |
| 15N124 | *H. erato* | *venus* | Colombia | Valle del Cauca | Queremal | 3.53199 | -76.75461 | 5.38 | 11.73 | C | 0.87 | ERS3546513 |
| 15N147 | *H. erato* | *venus* | Colombia | Valle del Cauca | Queremal | 3.53199 | -76.75461 | 5.03 | 11.46 | C | 0.95 | ERS3546514 |
| 15N148 | *H. erato* | *venus* | Colombia | Valle del Cauca | Queremal | 3.53199 | -76.75461 | 3.98 | 9.83 | C | 0.50 | ERS3546515 |
| 15N155 | *H. erato* | *venus* | Colombia | Valle del Cauca | Queremal | 3.53199 | -76.75461 | 6.92 | 12.45 | C | 0.90 | ERS3546516 |
| 15N158 | *H. erato* | *venus* | Colombia | Valle del Cauca | La Elsa - Rio Dagua | 3.567778 | -76.776111 | 6.43 | 12.72 | C | 0.92 | ERS3546517 |
| 15N159 | *H. erato* | *venus* | Colombia | Valle del Cauca | La Elsa - Rio Dagua | 3.567778 | -76.776111 | 4.36 | 10.47 | C | 0.99 | ERS3546518 |
| 15N163 | *H. erato* | *venus* | Colombia | Valle del Cauca | La Elsa - Rio Dagua | 3.567778 | -76.776111 | 5.66 | 0.06 | C | 0.85 | ERS3546519 |

| | | | | | | | | | | | | |
|---|---|---|---|---|---|---|---|---|---|---|---|---|
| 15N164 | *H. erato* | *venus* | Colombia | Valle del Cauca | La Elsa - Rio Dagua | 3.567778 | -76.776111 | 7.22 | 13.36 | C | 0.82 | ERS3546520 |
| 15N165 | *H. erato* | *venus* | Colombia | Valle del Cauca | La Elsa - Rio Dagua | 3.567778 | -76.776111 | 5.55 | 11.96 | C | 0.91 | ERS3546521 |
| 15N166 | *H. erato* | *venus* | Colombia | Valle del Cauca | La Elsa - Rio Dagua | 3.567778 | -76.776111 | 3.86 | 9.71 | C | 0.92 | ERS3546522 |
| 15N167 | *H. erato* | *venus* | Colombia | Valle del Cauca | La Elsa - Rio Dagua | 3.567778 | -76.776111 | 6.69 | 12.86 | C | 0.78 | ERS3546523 |
| 15N201 | *H. erato* | *venus* | Colombia | Valle del Cauca | San Pedro | 3.83691 | -77.25734 | 6.85 | 12.94 | C | 0.95 | ERS3546524 |
| 15N203 | *H. erato* | *venus* | Colombia | Valle del Cauca | San Pedro | 3.83691 | -77.25734 | 4.61 | 10.8 | C | 0.58 | ERS3546525 |
| 15N204 | *H. erato* | *venus* | Colombia | Valle del Cauca | San Pedro | 3.83691 | -77.25734 | 12.53 | 16.4 | C | 0.99 | ERS3546526 |
| 15N205 | *H. erato* | *venus* | Colombia | Valle del Cauca | San Pedro | 3.83691 | -77.25734 | 4.99 | 11.16 | C | 0.71 | ERS3546527 |
| 15N206 | *H. erato* | *venus* | Colombia | Valle del Cauca | San Pedro | 3.83691 | -77.25734 | 4.79 | 10.68 | C | 0.91 | ERS3546528 |
| 15N207 | *H. erato* | *venus* | Colombia | Valle del Cauca | San Pedro | 3.83691 | -77.25734 | 4.94 | 10.97 | C | 0.96 | ERS3546529 |
| 15N208 | *H. erato* | *venus* | Colombia | Valle del Cauca | San Pedro | 3.83691 | -77.25734 | 3.82 | 9.34 | C | 0.74 | ERS3546530 |
| 15N209 | *H. erato* | *venus* | Colombia | Valle del Cauca | San Pedro | 3.83691 | -77.25734 | 2.84 | 7.41 | C | 0.91 | ERS3546531 |
| 15N214 | *H. erato* | *venus* | Colombia | Valle del Cauca | San Pedro | 3.83691 | -77.25734 | 4.48 | 10.33 | C | 0.98 | ERS3546532 |

| | | | | | | | | | | | | |
|---|---|---|---|---|---|---|---|---|---|---|---|---|
| 15N215 | *H. erato* | *venus* | Colombia | Valle del Cauca | San Pedro | 3.83691 | -77.25734 | 4.82 | 10.37 | C | 0.88 | ERS3546533 |
| 15N216 | *H. erato* | *venus* | Colombia | Valle del Cauca | San Pedro | 3.83691 | -77.25734 | 6.84 | 12.94 | C | 0.78 | ERS3546534 |
| 15N218 | *H. erato* | *venus* | Colombia | Valle del Cauca | San Pedro | 3.83691 | -77.25734 | 5.94 | 12.02 | C | 0.71 | ERS3546535 |
| 15N220 | *H. erato* | *venus* | Colombia | Valle del Cauca | San Pedro | 3.83691 | -77.25734 | 6.05 | 12.31 | C | 0.83 | ERS3546536 |
| 15N221 | *H. erato* | *venus* | Colombia | Valle del Cauca | San Pedro | 3.83691 | -77.25734 | 5.63 | 11.76 | C | 0.63 | ERS3546537 |
| 15N222 | *H. erato* | *venus* | Colombia | Valle del Cauca | San Pedro | 3.83691 | -77.25734 | 9.16 | 14.86 | C | 0.92 | ERS3546538 |
| 15N224 | *H. erato* | *venus* | Colombia | Valle del Cauca | San Pedro | 3.83691 | -77.25734 | 7.12 | 13.29 | C | 0.81 | ERS3546539 |
| 15N225 | *H. erato* | *venus* | Colombia | Valle del Cauca | San Pedro | 3.83691 | -77.25734 | 5.74 | 11.93 | C | 0.85 | ERS3546540 |
| 15N226 | *H. erato* | *venus* | Colombia | Valle del Cauca | San Pedro | 3.83691 | -77.25734 | 9.68 | 14.99 | C | 0.86 | ERS3546541 |
| 15N227 | *H. erato* | *venus* | Colombia | Valle del Cauca | San Pedro | 3.83691 | -77.25734 | 7.5 | 13.42 | C | 0.97 | ERS3546542 |
| 15N228 | *H. erato* | *venus* | Colombia | Valle del Cauca | San Pedro | 3.83691 | -77.25734 | 3.52 | 8.97 | C | 0.89 | ERS3546543 |
| 15N231 | *H. erato* | *venus* | Colombia | Valle del Cauca | San Pedro | 3.83691 | -77.25734 | 7.45 | 13.34 | C | 0.89 | ERS3546544 |
| 15N235 | *H. erato* | *venus* | Colombia | Valle del Cauca | San Pedro | 3.83691 | -77.25734 | 14.52 | 17.38 | C | 0.80 | ERS3546545 |

| | | | | | | | | | | | | |
|---|---|---|---|---|---|---|---|---|---|---|---|---|
| 15N236 | *H. erato* | *venus* | Colombia | Valle del Cauca | San Pedro | 3.83691 | -77.25734 | 3.28 | 8.38 | C | 0.82 | ERS3546546 |
| 15N237 | *H. erato* | *venus* | Colombia | Valle del Cauca | San Pedro | 3.83691 | -77.25734 | 6.98 | 12.96 | C | 0.56 | ERS3546547 |
| 275 | *H. erato* | *venus* | Colombia | Valle del Cauca | Ladrilleros | 3.939444 | -77.36889 | 3.06 | 8.05 | C | 0.78 | ERS3546548 |
| 278 | *H. erato* | *venus* | Colombia | Valle del Cauca | Ladrilleros | 3.939444 | -77.36889 | 2.27 | 5.64 | C | 0.37 | ERS3546549 |
| 282 | *H. erato* | *venus* | Colombia | Valle del Cauca | Ladrilleros | 3.939444 | -77.36889 | 5.12 | 10.93 | C | 0.70 | ERS3546550 |
| 1039 | *H. erato* | *venus* | Colombia | Valle del Cauca | Ladrilleros | 3.939444 | -77.36889 | 8.03 | 13.56 | C | 0.75 | ERS3546551 |
| 2471 | *H. erato* | *venus* | Colombia | Valle del Cauca | Ladrilleros | 3.939444 | -77.36889 | 9.53 | 14.57 | C | 0.83 | ERS3546552 |
| 251 | *H. erato* | *venus* | Colombia | Valle del Cauca | La Barra | 3.958333 | -77.37333 | 4.39 | 10.02 | C | 0.73 | ERS3546553 |
| 283 | *H. erato* | *venus* | Colombia | Valle del Cauca | La Barra | 3.958333 | -77.37333 | 7.21 | 13 | C | 0.90 | ERS3546554 |
| 286 | *H. erato* | *venus* | Colombia | Valle del Cauca | La Barra | 3.958333 | -77.37333 | 4 | 9.53 | C | 0.59 | ERS3546555 |
| 2626 | *H. erato* | *venus* | Colombia | Valle del Cauca | La Barra | 3.958333 | -77.37333 | 7.73 | 13.39 | C | 0.90 | ERS3546556 |
| 2631 | *H. erato* | *venus* | Colombia | Valle del Cauca | La Barra | 3.958333 | -77.37333 | 5.59 | 11.42 | C | 0.88 | ERS3546557 |
| 2633 | *H. erato* | *venus* | Colombia | Valle del Cauca | La Barra | 3.958333 | -77.37333 | 5.05 | 10.86 | C | 0.47 | ERS3546558 |

| | | | | | | | | | | | | |
|---|---|---|---|---|---|---|---|---|---|---|---|---|
| 2635 | *H. erato* | *venus* | Colombia | Valle del Cauca | La Barra | 3.958333 | -77.37333 | 4.22 | 9.86 | C | 0.27 | ERS3546559 |
| 2636 | *H. erato* | *venus* | Colombia | Valle del Cauca | La Barra | 3.958333 | -77.37333 | 3.19 | 8.11 | C | 0.78 | ERS3546560 |
| 2637 | *H. erato* | *venus* | Colombia | Valle del Cauca | La Barra | 3.958333 | -77.37333 | 3.61 | 8.83 | C | 0.87 | ERS3546561 |
| 15N242 | *H. erato* | *venus* | Colombia | Choco | Amargal | 5.57187 | -77.50211 | 4.86 | 10.98 | C | 0.87 | ERS3546562 |
| 15N244 | *H. erato* | *venus* | Colombia | Choco | Amargal | 5.57187 | -77.50211 | 2.93 | 6.78 | C | 0.82 | ERS3546563 |
| 15N245 | *H. erato* | *venus* | Colombia | Choco | Amargal | 5.57187 | -77.50211 | 3.89 | 8.35 | C | 0.82 | ERS3546564 |
| 15N248 | *H. erato* | *venus* | Colombia | Choco | Amargal | 5.57187 | -77.50211 | 8.55 | 12.82 | C | 0.61 | ERS3546565 |
| 15N251 | *H. erato* | *venus* | Colombia | Choco | Amargal | 5.57187 | -77.50211 | 3.94 | 8.45 | C | 0.87 | ERS3546566 |
| 15N252 | *H. erato* | *venus* | Colombia | Choco | Amargal | 5.57187 | -77.50211 | 7.22 | 12.25 | C | 0.82 | ERS3546567 |
| 15N253 | *H. erato* | *venus* | Colombia | Choco | Amargal | 5.57187 | -77.50211 | 4 | 8.49 | C | 0.93 | ERS3546568 |
| 15N255 | *H. erato* | *venus* | Colombia | Choco | Amargal | 5.57187 | -77.50211 | 5.63 | 10.61 | C | 0.94 | ERS3546569 |
| 15N256 | *H. erato* | *venus* | Colombia | Choco | Amargal | 5.57187 | -77.50211 | 4.86 | 9.95 | C | 0.74 | ERS3546570 |
| 15N257 | *H. erato* | *venus* | Colombia | Choco | Amargal | 5.57187 | -77.50211 | 5.95 | 10.73 | C | 0.89 | ERS3546571 |
| 15N258 | *H. erato* | *venus* | Colombia | Choco | Amargal | 5.57187 | -77.50211 | 7.23 | 12.04 | C | 0.86 | ERS3546572 |
| 15N262 | *H. erato* | *venus* | Colombia | Choco | Amargal | 5.57187 | -77.50211 | 5 | 9.99 | C | 0.97 | ERS3546573 |
| 15N263 | *H. erato* | *venus* | Colombia | Choco | Amargal | 5.57187 | -77.50211 | 13.31 | 15.49 | C | 0.97 | ERS3546574 |
| 15N264 | *H. erato* | *venus* | Colombia | Choco | Amargal | 5.57187 | -77.50211 | 4.46 | 9.42 | C | 0.87 | ERS3546575 |
| 15N265 | *H. erato* | *venus* | Colombia | Choco | Amargal | 5.57187 | -77.50211 | 5.51 | 10.4 | C | 0.96 | ERS3546576 |
| 15N266 | *H. erato* | *venus* | Colombia | Choco | Amargal | 5.57187 | -77.50211 | 4.17 | 8.92 | C | 0.94 | ERS3546577 |

| | | | | | | | | | | | | |
|---|---|---|---|---|---|---|---|---|---|---|---|---|
| 15N269 | *H. erato* | *venus* | Colombia | Choco | Amargal | 5.57187 | -77.50211 | 5.08 | 10.11 | C | 0.98 | ERS3546578 |
| 15N271 | *H. erato* | *venus* | Colombia | Choco | Amargal | 5.57187 | -77.50211 | 3.47 | 7.85 | C | 0.99 | ERS3546579 |
| 15N276 | *H. erato* | *venus* | Colombia | Choco | Amargal | 5.57187 | -77.50211 | 14.45 | 15.77 | C | 0.86 | ERS3546580 |
| 15N277 | *H. erato* | *venus* | Colombia | Choco | Amargal | 5.57187 | -77.50211 | 3.47 | 7.81 | C | 0.95 | ERS3546581 |
| 15N278 | *H. erato* | *venus* | Colombia | Choco | Amargal | 5.57187 | -77.50211 | 5.01 | 10.01 | C | 0.99 | ERS3546582 |
| 15N280 | *H. erato* | *venus* | Colombia | Choco | Amargal | 5.57187 | -77.50211 | 6.53 | 11.42 | C | 0.80 | ERS3546583 |
| 15N285 | *H. erato* | *venus* | Colombia | Choco | Amargal | 5.57187 | -77.50211 | 6.4 | 12.3 | C | 0.99 | ERS3546584 |
| 15N287 | *H. erato* | *venus* | Colombia | Choco | Amargal | 5.57187 | -77.50211 | 2.6 | 6.69 | C | 0.94 | ERS3546585 |
| 15N289 | *H. erato* | *venus* | Colombia | Choco | Amargal | 5.57187 | -77.50211 | 6.1 | 11.95 | C | 0.92 | ERS3546586 |
| 15N290 | *H. erato* | *venus* | Colombia | Choco | Amargal | 5.57187 | -77.50211 | 3.27 | 8.26 | C | 0.90 | ERS3546587 |
| 15N291 | *H. erato* | *venus* | Colombia | Choco | Amargal | 5.57187 | -77.50211 | 5.31 | 11.21 | C | 0.96 | ERS3546588 |
| 15N294 | *H. erato* | *venus* | Colombia | Choco | Amargal | 5.57187 | -77.50211 | 5.96 | 11.91 | C | 0.69 | ERS3546589 |
| 15N295 | *H. erato* | *venus* | Colombia | Choco | Amargal | 5.57187 | -77.50211 | 5.92 | 11.87 | C | 0.98 | ERS3546590 |
| 15N296 | *H. erato* | *venus* | Colombia | Choco | Amargal | 5.57187 | -77.50211 | 6.1 | 12.09 | C | 0.97 | ERS3546591 |
| 15N301 | *H. erato* | *venus* | Colombia | Choco | Amargal | 5.57187 | -77.50211 | 3.5 | 8.61 | C | 0.83 | ERS3546592 |
| 15N302 | *H. erato* | *venus* | Colombia | Choco | Amargal | 5.57187 | -77.50211 | 3.16 | 8.09 | C | 0.97 | ERS3546593 |
| 15N304 | *H. erato* | *venus* | Colombia | Choco | Amargal | 5.57187 | -77.50211 | 2.4 | 6.14 | C | 0.85 | ERS3546594 |
| 15N307 | *H. erato* | *venus* | Colombia | Choco | Amargal | 5.57187 | -77.50211 | 4.06 | 9.71 | C | 0.85 | ERS3546595 |
| 15N308 | *H. erato* | *venus* | Colombia | Choco | Amargal | 5.57187 | -77.50211 | 4.35 | 9.97 | C | 0.84 | ERS3546596 |
| 15N310 | *H. erato* | *venus* | Colombia | Choco | Amargal | 5.57187 | -77.50211 | 3.32 | 8.26 | C | 0.79 | ERS3546597 |
| 15N315 | *H. erato* | *venus* | Colombia | Choco | Amargal | 5.57187 | -77.50211 | 6.9 | 12.6 | C | 0.87 | ERS3546598 |

| | | | | | | | | | | | | |
|---|---|---|---|---|---|---|---|---|---|---|---|---|
| 15N316 | *H. erato* | *venus* | Colombia | Choco | Amargal | 5.57187 | -77.50211 | 18.53 | 18.17 | C | 0.97 | ERS3546599 |
| 15N317 | *H. erato* | *venus* | Colombia | Choco | Amargal | 5.57187 | -77.50211 | 4.6 | 10.37 | C | 0.88 | ERS3546600 |
| 16N008 | *H. erato* | *venus* | Colombia | Choco | Playa Parra | 6.24066 | -77.39544 | 8.61 | 15.17 | C | 0.30 | ERS3546601 |
| 16N010 | *H. erato* | *venus* | Colombia | Choco | Playa Parra | 6.24066 | -77.39544 | 3.59 | 8.66 | C | 0.77 | ERS3546602 |
| 16N014 | *H. erato* | *venus* | Colombia | Choco | Playa Parra | 6.24066 | -77.39544 | 3.52 | 8.64 | C | 0.59 | ERS3546603 |
| 16N015 | *H. erato* | *venus* | Colombia | Choco | Playa Parra | 6.24066 | -77.39544 | 5.48 | 11.5 | C | 0.90 | ERS3546604 |
| 16N017 | *H. erato* | *venus* | Colombia | Choco | Playa Parra | 6.24066 | -77.39544 | 3.99 | 9.5 | C | 0.95 | ERS3546605 |
| 16N019 | *H. erato* | *venus* | Colombia | Choco | Playa Parra | 6.24066 | -77.39544 | 7.28 | 13.26 | C | 0.99 | ERS3546606 |
| 16N096 | *H. erato* | *venus* | Colombia | Choco | Potes | 6.3495 | -77.36697 | 4.48 | 10.25 | C | 0.75 | ERS3546607 |
| 16N098 | *H. erato* | *venus* | Colombia | Choco | Potes | 6.3495 | -77.36697 | 6.07 | 12.2 | C | 0.97 | ERS3546608 |
| 16N101 | *H. erato* | *venus* | Colombia | Choco | Potes | 6.3495 | -77.36697 | 4.18 | 9.98 | C | 0.70 | ERS3546609 |
| 16N103 | *H. erato* | *venus* | Colombia | Choco | Potes | 6.3495 | -77.36697 | 5.98 | 12.5 | C | 0.90 | ERS3546610 |
| 16N104 | *H. erato* | *venus* | Colombia | Choco | Potes | 6.3495 | -77.36697 | 6.19 | 12.33 | C | 0.90 | ERS3546611 |
| 16N105 | *H. erato* | *venus* | Colombia | Choco | Potes | 6.3495 | -77.36697 | 4.93 | 10.87 | C | 0.89 | ERS3546612 |
| 16N106 | *H. erato* | *venus* | Colombia | Choco | Potes | 6.3495 | -77.36697 | 10.59 | 15.62 | C | 0.95 | ERS3546613 |
| 16N107 | *H. erato* | *venus* | Colombia | Choco | Potes | 6.3495 | -77.36697 | 4.73 | 10.41 | C | 1.00 | ERS3546614 |
| 16N109 | *H. erato* | *venus* | Colombia | Choco | Potes | 6.3495 | -77.36697 | 9.35 | 14.85 | C | 0.89 | ERS3546615 |
| 16N112 | *H. erato* | *venus* | Colombia | Choco | Potes | 6.3495 | -77.36697 | 8.31 | 13.94 | C | 0.80 | ERS3546616 |
| 16N113 | *H. erato* | *venus* | Colombia | Choco | Potes | 6.3495 | -77.36697 | 6.09 | 12.19 | C | 0.96 | ERS3546617 |
| 16N114 | *H. erato* | *venus* | Colombia | Choco | Potes | 6.3495 | -77.36697 | 3.45 | 8.57 | C | 0.73 | ERS3546618 |
| 16N115 | *H. erato* | *venus* | Colombia | Choco | Potes | 6.3495 | -77.36697 | 4.81 | 10.63 | C | 0.92 | ERS3546619 |

| | | | | | | | | | | | | |
|---|---|---|---|---|---|---|---|---|---|---|---|---|
| 16N116 | *H. erato* | *venus* | Colombia | Choco | Potes | 6.3495 | -77.36697 | 4.42 | 10.22 | C | 0.93 | ERS3546620 |
| 16N118 | *H. erato* | *venus* | Colombia | Choco | Potes | 6.3495 | -77.36697 | 11.4 | 15.57 | C | 0.95 | ERS3546621 |
| 16N119 | *H. erato* | *venus* | Colombia | Choco | Potes | 6.3495 | -77.36697 | 5.92 | 13.89 | C | 0.86 | ERS3546622 |
| 16N121 | *H. erato* | *venus* | Colombia | Choco | Potes | 6.3495 | -77.36697 | 4.15 | 9.86 | C | 0.81 | ERS3546623 |
| 16N125 | *H. erato* | *venus* | Colombia | Choco | Potes | 6.3495 | -77.36697 | 5.29 | 11.17 | C | 0.80 | ERS3546624 |
| 16N050 | *H. erato* | *venus* | Colombia | Choco | Cocalito | 6.38504 | -77.40251 | 3.77 | 9.07 | C | 0.75 | ERS3546625 |
| 16N053 | *H. erato* | *venus* | Colombia | Choco | Cocalito | 6.38504 | -77.40251 | 5.24 | 11.24 | C | 0.88 | ERS3546626 |
| 16N054 | *H. erato* | *venus* | Colombia | Choco | Cocalito | 6.38504 | -77.40251 | 5.79 | 11.83 | C | 0.83 | ERS3546627 |
| 16N055 | *H. erato* | *venus* | Colombia | Choco | Cocalito | 6.38504 | -77.40251 | 3.27 | 8.06 | C | 0.57 | ERS3546628 |
| 16N060 | *H. erato* | *venus* | Colombia | Choco | Cocalito | 6.38504 | -77.40251 | 5.08 | 11 | C | 0.62 | ERS3546629 |
| 16N061 | *H. erato* | *venus* | Colombia | Choco | Cocalito | 6.38504 | -77.40251 | 6.43 | 12.4 | C | 0.50 | ERS3546630 |
| 16N062 | *H. erato* | *venus* | Colombia | Choco | Cocalito | 6.38504 | -77.40251 | 5.95 | 12.06 | C | 0.85 | ERS3546631 |
| 16N065 | *H. erato* | *venus* | Colombia | Choco | Cocalito | 6.38504 | -77.40251 | 6.66 | 12.76 | C | 0.69 | ERS3546632 |
| 16N067 | *H. erato* | *venus* | Colombia | Choco | Cocalito | 6.38504 | -77.40251 | 8.61 | 14.3 | C | 0.87 | ERS3546633 |
| 16N069 | *H. erato* | *venus* | Colombia | Choco | Cocalito | 6.38504 | -77.40251 | 6.45 | 12.67 | C | 0.41 | ERS3546634 |
| 16N070 | *H. erato* | *venus* | Colombia | Choco | Cocalito | 6.38504 | -77.40251 | 3.85 | 9.41 | C | 0.70 | ERS3546635 |
| 16N076 | *H. erato* | *venus* | Colombia | Choco | Cocalito | 6.38504 | -77.40251 | 5.81 | 11.9 | C | 0.94 | ERS3546636 |
| 16N079 | *H. erato* | *venus* | Colombia | Choco | Cocalito | 6.38504 | -77.40251 | 4.93 | 10.93 | C | 0.72 | ERS3546637 |
| 16N082 | *H. erato* | *venus* | Colombia | Choco | Cocalito | 6.38504 | -77.40251 | 5.95 | 12.08 | C | 0.79 | ERS3546638 |
| 16N021 | *H. erato* | *venus* | Colombia | Choco | Playa Flores | 6.38765 | -77.38557 | 4.23 | 9.84 | C | 0.74 | ERS3546639 |
| 16N024 | *H. erato* | *venus* | Colombia | Choco | Playa Flores | 6.38765 | -77.38557 | 4.03 | 9.6 | C | 0.89 | ERS3546640 |

| | | | | | | | | | | | | |
|---|---|---|---|---|---|---|---|---|---|---|---|---|
| 16N025 | *H. erato* | *venus* | Colombia | Choco | Playa Flores | 6.38765 | -77.38557 | 3.21 | 7.82 | C | 0.90 | ERS3546641 |
| 16N026 | *H. erato* | *venus* | Colombia | Choco | Playa Flores | 6.38765 | -77.38557 | 5.43 | 11.34 | C | 0.57 | ERS3546642 |
| 16N028 | *H. erato* | *venus* | Colombia | Choco | Playa Flores | 6.38765 | -77.38557 | 3.37 | 8.26 | C | 0.93 | ERS3546643 |
| 16N039 | *H. erato* | *venus* | Colombia | Choco | Playa Flores | 6.38765 | -77.38557 | 3.59 | 8.8 | C | 0.93 | ERS3546644 |
| 16N041 | *H. erato* | *venus* | Colombia | Choco | Playa Flores | 6.38765 | -77.38557 | 6.89 | 12.87 | C | 0.91 | ERS3546645 |
| 16N044 | *H. erato* | *venus* | Colombia | Choco | Playa Flores | 6.38765 | -77.38557 | 4.36 | 10.15 | C | 0.83 | ERS3546646 |
| 16N045 | *H. erato* | *venus* | Colombia | Choco | Playa Flores | 6.38765 | -77.38557 | 7.49 | 13.33 | C | 0.92 | ERS3546647 |
| 16N046 | *H. erato* | *venus* | Colombia | Choco | Playa Flores | 6.38765 | -77.38557 | 6.38 | 12.53 | C | 0.93 | ERS3546648 |
| 15N324 | *H. erato* | *hydara* | Panama | Darien | Jaque | 7.48767 | -78.12733 | 6.57 | 12.96 | B | -0.25 | ERS3546649 |
| 15N330 | *H. erato* | *hydara* | Panama | Darien | Jaque | 7.48767 | -78.12733 | 9.38 | 14.94 | B | 0.12 | ERS3546650 |
| 15N331 | *H. erato* | *hydara* | Panama | Darien | Jaque | 7.48767 | -78.12733 | 12.86 | 16.74 | B | -0.24 | ERS3546651 |
| 15N332 | *H. erato* | *hydara* | Panama | Darien | Jaque | 7.48767 | -78.12733 | 7.68 | 13.86 | B | -0.25 | ERS3546652 |
| 15N336 | *H. erato* | *venus x hydara* | Panama | Darien | Jaque | 7.48767 | -78.12733 | 6.54 | 12.96 | B | -0.17 | ERS3546653 |
| 15N337 | *H. erato* | *hydara* | Panama | Darien | Jaque | 7.48767 | -78.12733 | 6.82 | 14.33 | A | 0.14 | ERS3546654 |
| 15N344 | *H. erato* | *hydara* | Panama | Darien | Jaque | 7.48767 | -78.12733 | 5.06 | 12.41 | A | -0.15 | ERS3546655 |
| 15N345 | *H. erato* | *hydara* | Panama | Darien | Jaque | 7.48767 | -78.12733 | 4.89 | 12.17 | B | -0.09 | ERS3546656 |
| 15N346 | *H. erato* | *hydara* | Panama | Darien | Jaque | 7.48767 | -78.12733 | 3.65 | 10.4 | A | 0.28 | ERS3546657 |
| 15N347 | *H. erato* | *hydara* | Panama | Darien | Jaque | 7.48767 | -78.12733 | 3.23 | 9.35 | B | -0.08 | ERS3546658 |
| 15N348 | *H. erato* | *hydara* | Panama | Darien | Jaque | 7.48767 | -78.12733 | 3.15 | 9.21 | B | -0.23 | ERS3546659 |
| 15N349 | *H. erato* | *hydara* | Panama | Darien | Jaque | 7.48767 | -78.12733 | 2.98 | 8.65 | A | 0.00 | ERS3546660 |

| | | | | | | | | | | | | |
|---|---|---|---|---|---|---|---|---|---|---|---|---|
| 15N357 | *H. erato* | *hydara* | Panama | Darien | Jaque | 7.48767 | -78.12733 | 3.24 | 9.42 | B | -0.12 | ERS3546661 |
| 15N358 | *H. erato* | *hydara* | Panama | Darien | Jaque | 7.48767 | -78.12733 | 11.57 | 17.58 | A | -0.04 | ERS3546662 |
| 15N359 | *H. erato* | *hydara* | Panama | Darien | Jaque | 7.48767 | -78.12733 | 5.15 | 12.44 | A | 0.12 | ERS3546663 |
| 15N360 | *H. erato* | *hydara* | Panama | Darien | Jaque | 7.48767 | -78.12733 | 12.47 | 17.91 | A | 0.00 | ERS3546664 |
| 15N361 | *H. erato* | *hydara* | Panama | Darien | Jaque | 7.48767 | -78.12733 | 5.15 | 12.57 | A | 0.06 | ERS3546665 |
| 15N362 | *H. erato* | *hydara* | Panama | Darien | Jaque | 7.48767 | -78.12733 | 5.29 | 12.69 | A | -0.10 | ERS3546666 |
| 15N363 | *H. erato* | *hydara* | Panama | Darien | Jaque | 7.48767 | -78.12733 | 1.97 | 5.51 | A | -0.04 | ERS3546667 |
| 15N364 | *H. erato* | *hydara* | Panama | Darien | Jaque | 7.48767 | -78.12733 | 3.85 | 10.74 | B | 0.12 | ERS3546668 |
| 15N365 | *H. erato* | *hydara* | Panama | Darien | Jaque | 7.48767 | -78.12733 | 3.4 | 9.93 | A | 0.09 | ERS3546669 |
| 15N368 | *H. erato* | *venus* | Panama | Darien | Jaque | 7.48767 | -78.12733 | 3.51 | 10.11 | C | 0.06 | ERS3546670 |
| 15N369 | *H. erato* | *hydara* | Panama | Darien | Jaque | 7.48767 | -78.12733 | 5.7 | 13.2 | B | -0.13 | ERS3546671 |
| 15N371 | *H. erato* | *demophoon* | Panama | Darien | Jaque | 7.48767 | -78.12733 | 5.98 | 13.37 | D | -0.18 | ERS3546672 |
| 15N374 | *H. erato* | *hydara* | Panama | Darien | Jaque | 7.48767 | -78.12733 | 3.27 | 9.53 | A | -0.07 | ERS3546673 |
| 15N375 | *H. erato* | *hydara* | Panama | Darien | Jaque | 7.48767 | -78.12733 | 6.52 | 14.26 | A | 0.04 | ERS3546674 |
| 15N378 | *H. erato* | *hydara* | Panama | Darien | Jaque | 7.48767 | -78.12733 | 6.45 | 13.9 | A | -0.11 | ERS3546675 |
| 15N381 | *H. erato* | *hydara* | Panama | Darien | Jaque | 7.48767 | -78.12733 | 3.31 | 9.12 | B | -0.17 | ERS3546676 |
| 15N385 | *H. erato* | *hydara* | Panama | Darien | Jaque | 7.48767 | -78.12733 | 5.55 | 12.52 | B | 0.01 | ERS3546677 |
| 15N386 | *H. erato* | *hydara* | Panama | Darien | Jaque | 7.48767 | -78.12733 | 7.04 | 14.1 | B | -0.06 | ERS3546678 |
| 15N388 | *H. erato* | *hydara* | Panama | Darien | Jaque | 7.48767 | -78.12733 | 4.88 | 11.71 | B | 0.26 | ERS3546679 |
| 15N392 | *H. erato* | *hydara* | Panama | Darien | Jaque | 7.48767 | -78.12733 | 8.06 | 14.95 | B | -0.20 | ERS3546680 |
| 15N393 | *H. erato* | *hydara* | Panama | Darien | Jaque | 7.48767 | -78.12733 | 3.42 | 9.35 | B | 0.00 | ERS3546681 |

| | | | | | | | | | | | | |
|---|---|---|---|---|---|---|---|---|---|---|---|---|
| 15N395 | *H. erato* | *venus* | Panama | Darien | Jaque | 7.48767 | -78.12733 | 3.33 | 9.35 | C | -0.17 | ERS3546682 |
| 15N396 | *H. erato* | *hydara* | Panama | Darien | Jaque | 7.48767 | -78.12733 | 5.13 | 12.14 | A | -0.23 | ERS3546683 |
| 15N400 | *H. erato* | *hydara* | Panama | Darien | Jaque | 7.48767 | -78.12733 | 4.12 | 10.62 | A | -0.05 | ERS3546684 |
| 15N401 | *H. erato* | *hydara* | Panama | Darien | Jaque | 7.48767 | -78.12733 | 5.24 | 12.3 | A | -0.15 | ERS3546685 |
| 15N402 | *H. erato* | *hydara* | Panama | Darien | Jaque | 7.48767 | -78.12733 | 5.13 | 12.27 | B | -0.14 | ERS3546686 |
| 15N405 | *H. erato* | *hydara* | Panama | Darien | Jaque | 7.48767 | -78.12733 | 5.42 | 12.5 | B | -0.15 | ERS3546687 |
| 15N406 | *H. erato* | *hydara* | Panama | Darien | Jaque | 7.48767 | -78.12733 | 4.29 | 10.94 | B | -0.01 | ERS3546688 |
| 15N407 | *H. erato* | *hydara* | Panama | Darien | Jaque | 7.48767 | -78.12733 | 3.15 | 8.82 | B | 0.27 | ERS3546689 |
| 15N412 | *H. erato* | *hydara* | Panama | Darien | Jaque | 7.48767 | -78.12733 | 4.29 | 11.15 | A | 0.22 | ERS3546690 |
| 15N413 | *H. erato* | *hydara* | Panama | Darien | Jaque | 7.48767 | -78.12733 | 4.47 | 11.21 | B | -0.16 | ERS3546691 |
| 15N414 | *H. erato* | *venus* | Panama | Darien | Jaque | 7.48767 | -78.12733 | 5.59 | 12.76 | C | -0.06 | ERS3546692 |
| 15N416 | *H. erato* | *hydara* | Panama | Darien | Jaque | 7.48767 | -78.12733 | 5.56 | 12.5 | A | -0.24 | ERS3546693 |
| 15N418 | *H. erato* | *hydara* | Panama | Darien | Jaque | 7.48767 | -78.12733 | 4.1 | 10.56 | B | -0.26 | ERS3546694 |
| 15N419 | *H. erato* | *hydara* | Panama | Darien | Jaque | 7.48767 | -78.12733 | 8.3 | 14.97 | A | -0.02 | ERS3546695 |
| 15N422 | *H. erato* | *hydara* | Panama | Darien | Jaque | 7.48767 | -78.12733 | 4.83 | 11.84 | B | -0.20 | ERS3546696 |
| 15N424 | *H. erato* | *hydara* | Panama | Darien | Jaque | 7.48767 | -78.12733 | 7.39 | 14.07 | B | 0.25 | ERS3546697 |
| 15N425 | *H. erato* | *hydara* | Panama | Darien | Jaque | 7.48767 | -78.12733 | 3.83 | 10.08 | A | -0.05 | ERS3546698 |
| 15N428 | *H. erato* | *hydara* | Panama | Darien | Jaque | 7.48767 | -78.12733 | 7.17 | 13.25 | B | -0.12 | ERS3546699 |
| 15N430 | *H. erato* | *hydara* | Panama | Darien | Jaque | 7.48767 | -78.12733 | 5.67 | 12.58 | A | -0.02 | ERS3546700 |
| 15N431 | *H. erato* | *venus* | Panama | Darien | Jaque | 7.48767 | -78.12733 | 3.08 | 8.71 | C | -0.15 | ERS3546701 |
| 15N433 | *H. erato* | *hydara* | Panama | Darien | Jaque | 7.48767 | -78.12733 | 3.77 | 9.86 | A | -0.29 | ERS3546702 |

| | | | | | | | | | | | | |
|---|---|---|---|---|---|---|---|---|---|---|---|---|
| 15N434 | *H. erato* | *hydara* | Panama | Darien | Jaque | 7.48767 | -78.12733 | 5.61 | 12.32 | A | -0.16 | ERS3546703 |
| 15N436 | *H. erato* | *hydara* | Panama | Darien | Jaque | 7.48767 | -78.12733 | 7.26 | 14.09 | B | 0.03 | ERS3546704 |
| 15N441 | *H. erato* | *hydara* | Panama | Darien | Jaque | 7.48767 | -78.12733 | 4.59 | 11.14 | A | -0.38 | ERS3546705 |
| 15N442 | *H. erato* | *hydara* | Panama | Darien | Jaque | 7.48767 | -78.12733 | 5.1 | 11.97 | B | -0.10 | ERS3546706 |
| 15N444 | *H. erato* | *hydara* | Panama | Darien | Jaque | 7.48767 | -78.12733 | 5.01 | 11.85 | A | -0.31 | ERS3546707 |
| 15N445 | *H. erato* | *venus* | Panama | Darien | Jaque | 7.48767 | -78.12733 | 5.3 | 12.38 | C | -0.22 | ERS3546708 |
| 15N446 | *H. erato* | *hydara* | Panama | Darien | Jaque | 7.48767 | -78.12733 | 6.48 | 13.4 | B | -0.12 | ERS3546709 |
| 15N447 | *H. erato* | *hydara* | Panama | Darien | Jaque | 7.48767 | -78.12733 | 6.17 | 13.11 | A | 0.07 | ERS3546710 |
| 15N448 | *H. erato* | *hydara* | Panama | Darien | Jaque | 7.48767 | -78.12733 | 9.23 | 15.44 | A | -0.15 | ERS3546711 |
| 15N449 | *H. erato* | *venus* | Panama | Darien | Jaque | 7.48767 | -78.12733 | 6.17 | 13.09 | C | -0.22 | ERS3546712 |
| 15N450 | *H. erato* | *venus* | Panama | Darien | Jaque | 7.48767 | -78.12733 | 6.11 | 12.94 | C | -0.22 | ERS3546713 |
| 15N452 | *H. erato* | *venus* | Panama | Darien | Jaque | 7.48767 | -78.12733 | 5.59 | 12.53 | C | -0.06 | ERS3546714 |
| 15N453 | *H. erato* | *hydara* | Panama | Darien | Jaque | 7.48767 | -78.12733 | 4.9 | 11.74 | A | -0.06 | ERS3546715 |
| 15N454 | *H. erato* | *hydara* | Panama | Darien | Jaque | 7.48767 | -78.12733 | 2.87 | 8.02 | A | -0.10 | ERS3546716 |
| 18088 | *H. erato* | *hydara x demophoon* | Panama | Darien | Rancho Frio | 8.01974 | -77.73252 | 4.13 | 11.36 | B | -0.52 | ERS3546717 |
| 18089 | *H. erato* | *hydara x demophoon* | Panama | Darien | Rancho Frio | 8.01974 | -77.73252 | 6.33 | 13.35 | B | -0.44 | ERS3546718 |
| 18090 | *H. erato* | *hydara* | Panama | Darien | Rancho Frio | 8.01974 | -77.73252 | 3.76 | 9.8 | A | -0.54 | ERS3546719 |
| 18103 | *H. erato* | *hydara* | Panama | Darien | Rancho Frio | 8.01974 | -77.73252 | 6.72 | 13.64 | A | -0.44 | ERS3546720 |
| 18104 | *H. erato* | *hydara* | Panama | Darien | Rancho Frio | 8.01974 | -77.73252 | 2.99 | 8.26 | A | -0.44 | ERS3546721 |
| 18106 | *H. erato* | *hydara* | Panama | Darien | Rancho Frio | 8.01974 | -77.73252 | 6.09 | 13.07 | A | -0.47 | ERS3546722 |

| | | | | | | | | | | | | |
|---|---|---|---|---|---|---|---|---|---|---|---|---|
| 18107 | *H. erato* | *hydara* | Panama | Darien | Rancho Frio | 8.01974 | -77.73252 | 4.92 | 11.75 | A | -0.52 | ERS3546723 |
| 18108 | *H. erato* | *hydara* | Panama | Darien | Rancho Frio | 8.01974 | -77.73252 | 6.26 | 13.27 | A | -0.40 | ERS3546724 |
| 18109 | *H. erato* | *hydara* | Panama | Darien | Rancho Frio | 8.01974 | -77.73252 | 5.05 | 12.09 | A | -0.40 | ERS3546725 |
| 18110 | *H. erato* | *hydara* | Panama | Darien | Rancho Frio | 8.01974 | -77.73252 | 5.9 | 12.68 | A | -0.53 | ERS3546726 |
| 18111 | *H. erato* | *hydara* | Panama | Darien | Rancho Frio | 8.01974 | -77.73252 | 3.06 | 8.39 | A | -0.28 | ERS3546727 |
| 18113 | *H. erato* | *hydara* | Panama | Darien | Rancho Frio | 8.01974 | -77.73252 | 3.96 | 10.32 | A | -0.55 | ERS3546728 |
| 18114 | *H. erato* | *hydara* | Panama | Darien | Rancho Frio | 8.01974 | -77.73252 | 4.05 | 10.39 | A | -0.45 | ERS3546729 |
| 18116 | *H. erato* | *hydara* | Panama | Darien | Rancho Frio | 8.01974 | -77.73252 | 8.31 | 14.88 | A | -0.47 | ERS3546730 |
| 18118 | *H. erato* | *hydara x demophoon* | Panama | Darien | Rancho Frio | 8.01974 | -77.73252 | 1.93 | 4.88 | B | -0.47 | ERS3546731 |
| 18123 | *H. erato* | *hydara* | Panama | Darien | Yaviza | 8.15602 | -77.69308 | 3.32 | 8.77 | A | -0.36 | ERS3546732 |
| 18124 | *H. erato* | *hydara* | Panama | Darien | Yaviza | 8.15602 | -77.69308 | 5.14 | 11.42 | A | -0.49 | ERS3546733 |
| 18127 | *H. erato* | *hydara* | Panama | Darien | Yaviza | 8.15602 | -77.69308 | 5.75 | 12.06 | A | -0.32 | ERS3546734 |
| 18128 | *H. erato* | *hydara* | Panama | Darien | Yaviza | 8.15602 | -77.69308 | 3.43 | 8.84 | A | -0.45 | ERS3546735 |
| 18091 | *H. erato* | *hydara* | Panama | Darien | Santa Librada | 8.27971 | -77.80982 | 2.78 | 7.98 | A | -0.42 | ERS3546736 |
| 18095 | *H. erato* | *hydara* | Panama | Darien | Santa Librada | 8.27971 | -77.80982 | 5.2 | 12.01 | A | -0.43 | ERS3546737 |
| 18096 | *H. erato* | *hydara* | Panama | Darien | Santa Librada | 8.27971 | -77.80982 | 10.56 | 16.41 | B | -0.54 | ERS3546738 |
| 18146 | *H. erato* | *hydara* | Panama | Darien | Santa Librada | 8.27971 | -77.80982 | 4.69 | 11.55 | A | -0.46 | ERS3546739 |

| | | | | | | | | | | | | |
|---|---|---|---|---|---|---|---|---|---|---|---|---|
| 18151 | *H. erato* | *hydara* | Panama | Darien | Santa Librada | 8.27971 | -77.80982 | 7.56 | 14.37 | A | -0.43 | ERS3546740 |
| 18153 | *H. erato* | *hydara* | Panama | Darien | Santa Librada | 8.27971 | -77.80982 | 9.75 | 15.96 | A | -0.56 | ERS3546741 |
| 18164 | *H. erato* | *hydara x demophoon* | Panama | Darien | Santa Librada | 8.27971 | -77.80982 | 5.24 | 12.14 | B | -0.42 | ERS3546742 |
| 18166 | *H. erato* | *hydara* | Panama | Darien | Santa Librada | 8.27971 | -77.80982 | 6 | 13.02 | A | -0.45 | ERS3546743 |
| 18167 | *H. erato* | *hydara* | Panama | Darien | Santa Librada | 8.27971 | -77.80982 | 3.6 | 9.89 | A | -0.54 | ERS3546744 |
| 18170 | *H. erato* | *hydara* | Panama | Darien | Santa Librada | 8.27971 | -77.80982 | 3.18 | 9.02 | A | -0.37 | ERS3546745 |
| 18172 | *H. erato* | *hydara* | Panama | Darien | Santa Librada | 8.27971 | -77.80982 | 3.46 | 9.58 | A | -0.46 | ERS3546746 |
| 18006 | *H. erato* | *hydara x demophoon* | Panama | Darien | Puerto Lara | 8.61355 | -78.13982 | 3.68 | 10.01 | B | -0.53 | ERS3546747 |
| 18007 | *H. erato* | *hydara* | Panama | Darien | Puerto Lara | 8.61355 | -78.13982 | 3.4 | 9.4 | NA | NA | ERS3546748 |
| 18010 | *H. erato* | *hydara* | Panama | Darien | Puerto Lara | 8.61355 | -78.13982 | 4.22 | 10.86 | B | -0.42 | ERS3546749 |
| 18012 | *H. erato* | *hydara* | Panama | Darien | Puerto Lara | 8.61355 | -78.13982 | 2.93 | 8.2 | A | -0.29 | ERS3546750 |
| 18015 | *H. erato* | *hydara* | Panama | Darien | Puerto Lara | 8.61355 | -78.13982 | 3.56 | 9.69 | A | -0.45 | ERS3546751 |
| 18018 | *H. erato* | *hydara* | Panama | Darien | Puerto Lara | 8.61355 | -78.13982 | 5.86 | 13.03 | A | -0.50 | ERS3546752 |
| 18026 | *H. erato* | *hydara x demophoon* | Panama | Darien | Puerto Lara | 8.61355 | -78.13982 | 5.56 | 12.78 | B | -0.48 | ERS3546753 |
| 18028 | *H. erato* | *hydara* | Panama | Darien | Puerto Lara | 8.61355 | -78.13982 | 5.89 | 13.24 | A | -0.43 | ERS3546754 |

| | | | | | | | | | | | | |
|---|---|---|---|---|---|---|---|---|---|---|---|---|
| 18032 | *H. erato* | *hydara* | Panama | Darien | Puerto Lara | 8.61355 | -78.13982 | 8.75 | 15.74 | A | -0.40 | ERS3546755 |
| 18033 | *H. erato* | *hydara* | Panama | Darien | Puerto Lara | 8.61355 | -78.13982 | 9.35 | 16.18 | A | -0.45 | ERS3546756 |
| 18037 | *H. erato* | *hydara* | Panama | Darien | Puerto Lara | 8.61355 | -78.13982 | 6.52 | 13.76 | A | -0.56 | ERS3546757 |
| 18046 | *H. erato* | *hydara x demophoon* | Panama | Darien | Puerto Lara | 8.61355 | -78.13982 | 4.78 | 11.68 | B | -0.42 | ERS3546758 |
| 18050 | *H. erato* | *hydara x demophoon* | Panama | Darien | Puerto Lara | 8.61355 | -78.13982 | 4.07 | 10.71 | B | -0.51 | ERS3546759 |
| 18056 | *H. erato* | *hydara* | Panama | Darien | Puerto Lara | 8.61355 | -78.13982 | 3.73 | 10.27 | A | -0.46 | ERS3546760 |
| 18057 | *H. erato* | *hydara x demophoon* | Panama | Darien | Puerto Lara | 8.61355 | -78.13982 | 3.6 | 9.93 | B | -0.49 | ERS3546761 |
| 18063 | *H. erato* | *hydara* | Panama | Darien | Puerto Lara | 8.61355 | -78.13982 | 5.25 | 12.28 | A | -0.38 | ERS3546762 |
| 18077 | *H. erato* | *hydara* | Panama | Darien | Puerto Lara | 8.61355 | -78.13982 | 8.7 | 15.73 | A | -0.50 | ERS3546763 |

**Table S3.** Sequencing coverage, phenotype, and location information per individual for whole genome re-sequenced *Heliconius melpomene.* Sequencing coverage information was calculated after duplicates were removed, and before quality filtering. The bar category refers to the yellow hindwing bar category, defined by Mallet (1986). BR refers to the blue score, a ratio of the blue-red content of photographed wings, described in the Methods.

| ID | species | Race | Country | Province | Locality | Lat | Lon | Depth of coverage | % bases sequenced | bar category | BR | Accession (ENA) |
|---|---|---|---|---|---|---|---|---|---|---|---|---|
| 18003 | *H. melpomene* | *melpomene* | Panama | Darien | Puerto Lara | 8.61355 | -78.1398 | 26.43 | 97.38 | B | -0.44 | ERS3546765 |
| 18005 | *H. melpomene* | *melpomene* | Panama | Darien | Puerto Lara | 8.61355 | -78.1398 | 25.38 | 97.54 | A | -0.06 | ERS3546766 |
| 18014 | *H. melpomene* | *melpomene* | Panama | Darien | Puerto Lara | 8.61355 | -78.1398 | 21.31 | 97.45 | A | -0.47 | ERS3546767 |
| 18055 | *H. melpomene* | *melpomene* | Panama | Darien | Puerto Lara | 8.61355 | -78.1398 | 24.95 | 97.53 | NA | NA | ERS3546768 |
| 18085 | *H. melpomene* | *melpomene* | Panama | Darien | Puerto Lara | 8.61355 | -78.1398 | 25.13 | 97.47 | A | -0.2 | ERS3546769 |
| 18093 | *H. melpomene* | *melpomene* | Panama | Darien | Santa Librada | 8.27971 | -77.8098 | 25.63 | 97.51 | A | -0.36 | ERS3546770 |
| 18102 | *H. melpomene* | *melpomene* | Panama | Darien | Rancho Frio | 8.01974 | -77.7325 | 25.47 | 97.53 | B | -0.55 | ERS3546771 |
| 18120 | *H. melpomene* | *melpomene* | Panama | Darien | Yaviza | 8.15602 | -77.6931 | 25.32 | 97.67 | B | -0.55 | ERS3546772 |
| 18160 | *H. melpomene* | *melpomene* | Panama | Darien | Santa Librada | 8.27971 | -77.8098 | 25.92 | 97.55 | A | -0.49 | ERS3546773 |
| 18191 | *H. melpomene* | *melpomene* | Panama | Darien | Puerto Lajas Blancas | 8.41323 | -77.8092 | 26.7 | 97.54 | B | -0.66 | ERS3546774 |
| 18192 | *H. melpomene* | *melpomene* | Panama | Darien | Puerto Lajas Blancas | 8.41323 | -77.8092 | 28.14 | 97.54 | B | -0.59 | ERS3546775 |

| | | | | | | | | | | | | |
|---|---|---|---|---|---|---|---|---|---|---|---|---|
| 18193 | *H. melpomene* | *melpomene* | Panama | Darien | Santa Librada | 8.27971 | -77.8098 | 25.19 | 97.52 | A | -0.54 | ERS3546776 |
| 15N110 | *H. melpomene* | *vulcanus* | Colombia | Valle del Cauca | Queremal | 3.53199 | -76.7546 | 23.69 | 97.23 | C | -0.32 | ERS3546777 |
| 15N123 | *H. melpomene* | *vulcanus* | Colombia | Valle del Cauca | Queremal | 3.53199 | -76.7546 | 23.34 | 97.35 | C | -0.08 | ERS3546778 |
| 15N133 | *H. melpomene* | *vulcanus* | Colombia | Valle del Cauca | Rio Bravo | 3.53199 | -76.7546 | 26.46 | 97.33 | C | 0.15 | ERS3546779 |
| 15N143 | *H. melpomene* | *vulcanus* | Colombia | Valle del Cauca | Queremal | 3.53199 | -76.7546 | 24.46 | 98.36 | C | 0.03 | ERS3546780 |
| 15N144 | *H. melpomene* | *vulcanus* | Colombia | Valle del Cauca | Queremal | 3.53199 | -76.7546 | 24.53 | 98.25 | C | 0.03 | ERS3546781 |
| 15N151 | *H. melpomene* | *vulcanus* | Colombia | Valle del Cauca | Queremal | 3.53199 | -76.7546 | 22.24 | 97.24 | C | -0.09 | ERS3546782 |
| 15N152 | *H. melpomene* | *vulcanus* | Colombia | Valle del Cauca | Queremal | 3.53199 | -76.7546 | 23.72 | 97.35 | C | -0.26 | ERS3546783 |
| 15N153 | *H. melpomene* | *vulcanus* | Colombia | Valle del Cauca | Queremal | 3.53199 | -76.7546 | 26.5 | 97.37 | C | 0.16 | ERS3546784 |
| 15N154 | *H. melpomene* | *vulcanus* | Colombia | Valle del Cauca | Queremal | 3.53199 | -76.7546 | 24.54 | 97.31 | C | 0.28 | ERS3546785 |
| 15N172 | *H. melpomene* | *vulcanus* | Colombia | Valle del Cauca | Rio Bravo | 8.01974 | -77.7325 | 23.21 | 97.19 | C | -0.3 | ERS3546786 |
| 15N240 | *H. melpomene* | *vulcanus* | Colombia | Choco | Amargal | 5.57187 | -77.5021 | 22.87 | 97.41 | C | -0.26 | ERS3546787 |
| 15N260 | *H. melpomene* | *vulcanus* | Colombia | Choco | Amargal | 5.57187 | -77.5021 | 22.37 | 97.4 | C | 0.12 | ERS3546788 |
| 15N281 | *H. melpomene* | *vulcanus* | Colombia | Choco | Amargal | 5.57187 | -77.5021 | 22.07 | 97.31 | C | -0.2 | ERS3546789 |
| 15N299 | *H. melpomene* | *vulcanus* | Colombia | Choco | Amargal | 5.57187 | -77.5021 | 22.53 | 97.44 | C | 0.13 | ERS3546790 |
| 15N328 | *H. melpomene* | *melpomene* | Panama | Darien | Jaque | 7.48767 | -78.1273 | 24.51 | 97.53 | B | -0.33 | ERS3546791 |
| 15N338 | *H. melpomene* | *melpomene* | Panama | Darien | Jaque | 7.48767 | -78.1273 | 23.35 | 97.43 | B | -0.21 | ERS3546792 |

| | | | | | | | | | | | | |
|---|---|---|---|---|---|---|---|---|---|---|---|---|
| 15N353 | *H. melpomene* | *melpomene* | Panama | Darien | Jaque | 7.48767 | -78.1273 | 24.14 | 97.5 | A | -0.3 | ERS3546793 |
| 15N354 | *H. melpomene* | *vulcanus x rosina* | Panama | Darien | Jaque | 7.48767 | -78.1273 | 23.29 | 97.36 | C | -0.22 | ERS3546794 |
| 15N355 | *H. melpomene* | *melpomene* | Panama | Darien | Jaque | 7.48767 | -78.1273 | 23.93 | 97.53 | B | -0.38 | ERS3546795 |
| 15N387 | *H. melpomene* | *melpomene* | Panama | Darien | Jaque | 7.48767 | -78.1273 | 20.29 | 97.43 | B | -0.15 | ERS3546796 |
| 15N420 | *H. melpomene* | *melpomene* | Panama | Darien | Jaque | 7.48767 | -78.1273 | 21.41 | 97.47 | B | -0.31 | ERS3546797 |
| 15N421 | *H. melpomene* | *melpomene* | Panama | Darien | Jaque | 7.48767 | -78.1273 | 24.42 | 97.48 | B | -0.37 | ERS3546798 |
| 15N435 | *H. melpomene* | *vulcanus* | Panama | Darien | Jaque | 7.48767 | -78.1273 | 23.78 | 97.49 | C | -0.39 | ERS3546799 |
| 15N443 | *H. melpomene* | *vulcanus* | Panama | Darien | Jaque | 7.48767 | -78.1273 | 22.83 | 97.48 | C | -0.36 | ERS3546800 |

**Table S4.** Phenotype and location information for *Heliconius melpomene* and *Heliconius erato* individuals with no sequence data. The bar category refers to the yellow hindwing bar category, defined by Mallet (1986). BR refers to the blue score, a ratio of the blue-red content of photographed wings, described in the Methods.

| ID | species | race | Country | Province | Locality | Lat | Lon | bar category | BR |
|---|---|---|---|---|---|---|---|---|---|
| 3654 | *H. erato* | *venus* | Colombia | Valle del Cauca | Queremal | 3.53199 | -76.75461 | C | 0.49 |
| 3656 | *H. erato* | *venus* | Colombia | Valle del Cauca | Queremal | 3.53199 | -76.75461 | C | 0.50 |
| 3658 | *H. erato* | *venus* | Colombia | Valle del Cauca | Queremal | 3.53199 | -76.75461 | C | 0.54 |
| 15N104 | *H. erato* | *venus x chestertonii* | Colombia | Valle del Cauca | Queremal | 3.53199 | -76.75461 | C | 0.87 |
| 15N111 | *H. erato* | *venus* | Colombia | Valle del Cauca | Queremal | 3.53199 | -76.75461 | C | 0.83 |
| 15N112 | *H. erato* | *venus* | Colombia | Valle del Cauca | Queremal | 3.53199 | -76.75461 | C | 0.97 |
| 15N113 | *H. erato* | *venus* | Colombia | Valle del Cauca | Queremal | 3.53199 | -76.75461 | C | 0.67 |
| 15N120 | *H. erato* | *venus* | Colombia | Valle del Cauca | Queremal | 3.53199 | -76.75461 | C | 0.29 |
| 15N121 | *H. erato* | *venus* | Colombia | Valle del Cauca | Queremal | 3.53199 | -76.75461 | C | 0.92 |
| 15N160 | *H. erato* | *venus* | Colombia | Valle del Cauca | La Elsa - Rio Dagua | 3.567778 | -76.77611 | C | 0.89 |
| 15N192 | *H. erato* | *venus* | Colombia | Valle del Cauca | San Pedro | 3.83691 | -77.25734 | C | -0.04 |
| 15N193 | *H. erato* | *venus* | Colombia | Valle del Cauca | San Pedro | 3.83691 | -77.25734 | C | 0.89 |

| | | | | | | | | | |
|---|---|---|---|---|---|---|---|---|---|
| 15N194 | *H. erato* | *venus* | Colombia | Valle del Cauca | San Pedro | 3.83691 | -77.25734 | C | 0.68 |
| 15N195 | *H. erato* | *venus* | Colombia | Valle del Cauca | San Pedro | 3.83691 | -77.25734 | C | 0.87 |
| 15N196 | *H. erato* | *venus* | Colombia | Valle del Cauca | San Pedro | 3.83691 | -77.25734 | C | 0.96 |
| 15N197 | *H. erato* | *venus* | Colombia | Valle del Cauca | San Pedro | 3.83691 | -77.25734 | C | 0.85 |
| 288 | *H. erato* | *venus* | Colombia | Valle del Cauca | Ladrilleros | 3.93944 | -77.36889 | C | 0.73 |
| 631 | *H. erato* | *venus* | Colombia | Valle del Cauca | Ladrilleros | 3.93944 | -77.36889 | C | 0.55 |
| 635 | *H. erato* | *venus* | Colombia | Valle del Cauca | Ladrilleros | 3.93944 | -77.36889 | C | 0.50 |
| 641 | *H. erato* | *venus* | Colombia | Valle del Cauca | Ladrilleros | 3.93944 | -77.36889 | C | 0.40 |
| 643 | *H. erato* | *venus* | Colombia | Valle del Cauca | Ladrilleros | 3.93944 | -77.36889 | C | 0.73 |
| 644 | *H. erato* | *venus* | Colombia | Valle del Cauca | Ladrilleros | 3.93944 | -77.36889 | C | 0.89 |
| 2106 | *H. erato* | *venus* | Colombia | Valle del Cauca | Ladrilleros | 3.93944 | -77.36889 | C | 0.58 |
| 2279 | *H. erato* | *venus* | Colombia | Valle del Cauca | Ladrilleros | 3.93944 | -77.36889 | C | 0.58 |
| 2467 | *H. erato* | *venus* | Colombia | Valle del Cauca | Ladrilleros | 3.93944 | -77.36889 | C | 0.86 |
| 15N238 | *H. erato* | *venus* | Colombia | Chocó | Amargal | 5.57187 | -77.50211 | C | 0.83 |
| 15N239 | *H. erato* | *venus* | Colombia | Chocó | Amargal | 5.57187 | -77.50211 | C | 0.76 |
| 15N241 | *H. erato* | *venus* | Colombia | Chocó | Amargal | 5.57187 | -77.50211 | C | 0.77 |
| 15N246 | *H. erato* | *venus* | Colombia | Chocó | Amargal | 5.57187 | -77.50211 | C | 0.60 |
| 15N249 | *H. erato* | *venus* | Colombia | Chocó | Amargal | 5.57187 | -77.50211 | C | 0.83 |
| 15N250 | *H. erato* | *venus* | Colombia | Chocó | Amargal | 5.57187 | -77.50211 | C | 0.85 |
| 15N261 | *H. erato* | *venus* | Colombia | Chocó | Amargal | 5.57187 | -77.50211 | C | 0.69 |

| | | | | | | | | | |
|---|---|---|---|---|---|---|---|---|---|
| 15N279 | *H. erato* | *venus* | Colombia | Chocó | Amargal | 5.57187 | -77.50211 | C | 0.96 |
| 15N297 | *H. erato* | *venus* | Colombia | Chocó | Amargal | 5.57187 | -77.50211 | C | 0.96 |
| 15N303 | *H. erato* | *venus* | Colombia | Chocó | Amargal | 5.57187 | -77.50211 | C | 0.83 |
| 15N309 | *H. erato* | *venus* | Colombia | Chocó | Amargal | 5.57187 | -77.50211 | C | 0.98 |
| 15N311 | *H. erato* | *venus* | Colombia | Chocó | Amargal | 5.57187 | -77.50211 | C | 0.61 |
| 16N000 | *H. erato* | *venus* | Colombia | Chocó | Playa Parra | 6.24066 | -77.39544 | C | 0.57 |
| 16N001 | *H. erato* | *venus* | Colombia | Chocó | Playa Parra | 6.24066 | -77.39544 | C | 0.67 |
| 16N002 | *H. erato* | *venus* | Colombia | Chocó | Playa Parra | 6.24066 | -77.39544 | C | 0.61 |
| 16N006 | *H. erato* | *venus* | Colombia | Chocó | Playa Parra | 6.24066 | -77.39544 | C | 0.67 |
| 16N012 | *H. erato* | *venus* | Colombia | Chocó | Playa Parra | 6.24066 | -77.39544 | C | 0.78 |
| 16N013 | *H. erato* | *venus* | Colombia | Chocó | Playa Parra | 6.24066 | -77.39544 | C | 0.73 |
| 16N016 | *H. erato* | *venus* | Colombia | Chocó | Playa Parra | 6.24066 | -77.39544 | C | 0.86 |
| 16N100 | *H. erato* | *venus* | Colombia | Choco | Potes | 6.3495 | -77.36697 | C | 0.75 |
| 16N124 | *H. erato* | *venus* | Colombia | Choco | Potes | 6.3495 | -77.36697 | C | 0.33 |
| 16N064 | *H. erato* | *venus* | Colombia | Choco | Cocalito | 6.38504 | -77.40251 | C | 0.86 |
| 16N027 | *H. erato* | *venus* | Colombia | Chocó | Playa Flores | 6.38765 | -77.38557 | C | 0.89 |
| 16N029 | *H. erato* | *venus* | Colombia | Chocó | Playa Flores | 6.38765 | -77.38557 | C | 0.89 |
| 16N037 | *H. erato* | *venus* | Colombia | Chocó | Playa Flores | 6.38765 | -77.38557 | C | 0.82 |
| 16N038 | *H. erato* | *venus* | Colombia | Chocó | Playa Flores | 6.38765 | -77.38557 | C | 0.93 |
| 16N042 | *H. erato* | *venus* | Colombia | Chocó | Playa Flores | 6.38765 | -77.38557 | C | 0.84 |
| 15N320 | *H. erato* | *venus* | Panama | Daríen | Jaque | 7.48767 | -78.12733 | C | 0.09 |
| 15N321 | *H. erato* | *hydara* | Panama | Daríen | Jaque | 7.48767 | -78.12733 | B | -0.04 |
| 15N322 | *H. erato* | *hydara* | Panama | Daríen | Jaque | 7.48767 | -78.12733 | B | -0.03 |
| 15N323 | *H. erato* | *venus* | Panama | Daríen | Jaque | 7.48767 | -78.12733 | C | -0.15 |
| 15N340 | *H. erato* | *venus* | Panama | Daríen | Jaque | 7.48767 | -78.12733 | C | -0.22 |
| 15N376 | *H. erato* | *hydara* | Panama | Daríen | Jaque | 7.48767 | -78.12733 | B | -0.11 |

| | | | | | | | | | |
|---|---|---|---|---|---|---|---|---|---|
| 15N382 | *H. erato* | *hydara* | Panama | Daríen | Jaque | 7.48767 | -78.12733 | B | -0.17 |
| 15N384 | *H. erato* | *hydara* | Panama | Daríen | Jaque | 7.48767 | -78.12733 | B | -0.16 |
| 15N389 | *H. erato* | *venus* | Panama | Daríen | Jaque | 7.48767 | -78.12733 | C | -0.23 |
| 15N411 | *H. erato* | *hydara* | Panama | Daríen | Jaque | 7.48767 | -78.12733 | A | -0.31 |
| 15N437 | *H. erato* | *hydara* | Panama | Daríen | Jaque | 7.48767 | -78.12733 | B | -0.18 |
| 18087 | *H. erato* | *hydara* | Panama | Daríen | Rancho Frio | 8.01974 | -77.73252 | A | -0.42 |
| 18112 | *H. erato* | *hydara x demophoon* | Panama | Daríen | Rancho Frio | 8.01974 | -77.73252 | B | -0.46 |
| 18119 | *H. erato* | *hydara* | Panama | Daríen | Yaviza | 8.15602 | -77.69308 | A | -0.50 |
| 18121 | *H. erato* | *hydara* | Panama | Daríen | Yaviza | 8.15602 | -77.69308 | A | -0.37 |
| 18125 | *H. erato* | *hydara* | Panama | Daríen | Yaviza | 8.15602 | -77.69308 | A | -0.38 |
| 18129 | *H. erato* | *hydara* | Panama | Daríen | Yaviza | 8.15602 | -77.69308 | A | -0.39 |
| 18130 | *H. erato* | *hydara* | Panama | Daríen | Yaviza | 8.15602 | -77.69308 | A | -0.43 |
| 18131 | *H. erato* | *hydara* | Panama | Daríen | Yaviza | 8.15602 | -77.69308 | A | -0.44 |
| 18132 | *H. erato* | *hydara* | Panama | Daríen | Yaviza | 8.15602 | -77.69308 | A | -0.39 |
| 18133 | *H. erato* | *hydara* | Panama | Daríen | Yaviza | 8.15602 | -77.69308 | A | -0.52 |
| 18134 | *H. erato* | *hydara* | Panama | Daríen | Yaviza | 8.15602 | -77.69308 | A | -0.44 |
| 18135 | *H. erato* | *hydara* | Panama | Daríen | Yaviza | 8.15602 | -77.69308 | A | -0.62 |
| 18136 | *H. erato* | *hydara x demophoon* | Panama | Daríen | Yaviza | 8.15602 | -77.69308 | B | -0.56 |
| 18137 | *H. erato* | *hydara* | Panama | Daríen | Yaviza | 8.15602 | -77.69308 | A | -0.57 |
| 18138 | *H. erato* | *hydara* | Panama | Daríen | Yaviza | 8.15602 | -77.69308 | A | -0.46 |
| 18139 | *H. erato* | *hydara* | Panama | Daríen | Yaviza | 8.15602 | -77.69308 | A | -0.40 |
| 18140 | *H. erato* | *hydara x demophoon* | Panama | Daríen | Yaviza | 8.15602 | -77.69308 | B | -0.60 |
| 18141 | *H. erato* | *hydara* | Panama | Daríen | Yaviza | 8.15602 | -77.69308 | A | -0.41 |
| 18142 | *H. erato* | *hydara* | Panama | Daríen | Yaviza | 8.15602 | -77.69308 | A | -0.45 |

| | | | | | | | | | |
|---|---|---|---|---|---|---|---|---|---|
| 18144 | *H. erato* | *hydara x demophoon* | Panama | Daríen | Yaviza | 8.15602 | -77.69308 | B | -0.42 |
| 18177 | *H. erato* | *hydara* | Panama | Daríen | Yaviza | 8.15602 | -77.69308 | A | -0.54 |
| 18178 | *H. erato* | *hydara* | Panama | Daríen | Yaviza | 8.15602 | -77.69308 | A | -0.56 |
| 18179 | *H. erato* | *hydara* | Panama | Daríen | Yaviza | 8.15602 | -77.69308 | B | -0.57 |
| 18180 | *H. erato* | *hydara* | Panama | Daríen | Yaviza | 8.15602 | -77.69308 | A | -0.49 |
| 18181 | *H. erato* | *hydara* | Panama | Daríen | Yaviza | 8.15602 | -77.69308 | A | -0.48 |
| 18182 | *H. erato* | *hydara* | Panama | Daríen | Yaviza | 8.15602 | -77.69308 | A | -0.41 |
| 18183 | *H. erato* | *hydara* | Panama | Daríen | Yaviza | 8.15602 | -77.69308 | A | -0.48 |
| 18184 | *H. erato* | *hydara* | Panama | Daríen | Yaviza | 8.15602 | -77.69308 | A | -0.42 |
| 18185 | *H. erato* | *hydara* | Panama | Daríen | Yaviza | 8.15602 | -77.69308 | A | -0.45 |
| 18188 | *H. erato* | *hydara* | Panama | Daríen | Yaviza | 8.15602 | -77.69308 | A | -0.50 |
| 18190 | *H. erato* | *hydara x demophoon* | Panama | Daríen | Yaviza | 8.15602 | -77.69308 | B | -0.60 |
| 18158 | *H. erato* | *hydara* | Panama | Daríen | Santa Librada | 8.27971 | -77.80982 | A | -0.41 |
| 18168 | *H. erato* | *hydara* | Panama | Daríen | Santa Librada | 8.27971 | -77.80982 | A | -0.43 |
| 18086 | *H. erato* | *hydara* | Panama | Daríen | Meteti | 8.406767 | -77.99898 | A | -0.38 |
| WOM7762 | *H. erato* | *demophoon* | Panama | Coclé | El Valle | 8.59323 | -80.14108 | D | -0.46 |
| WOM7763 | *H. erato* | *hydara* | Panama | Coclé | El Valle | 8.59323 | -80.14108 | B | -0.61 |
| WOM7765 | *H. erato* | *demophoon* | Panama | Coclé | El Valle | 8.59323 | -80.14108 | D | -0.54 |
| WOM7766 | *H. erato* | *demophoon* | Panama | Coclé | El Valle | 8.59323 | -80.14108 | D | -0.40 |
| WOM7768 | *H. erato* | *demophoon* | Panama | Coclé | El Valle | 8.59323 | -80.14108 | D | -0.56 |
| WOM7770 | *H. erato* | *demophoon* | Panama | Coclé | El Valle | 8.59323 | -80.14108 | D | -0.58 |
| WOM7771 | *H. erato* | *demophoon* | Panama | Coclé | El Valle | 8.59323 | -80.14108 | D | -0.63 |
| WOM7772 | *H. erato* | *demophoon* | Panama | Coclé | El Valle | 8.59323 | -80.14108 | D | -0.51 |
| WOM7776 | *H. erato* | *demophoon* | Panama | Coclé | El Valle | 8.59323 | -80.14108 | D | -0.54 |

| | | | | | | | | | |
|---|---|---|---|---|---|---|---|---|---|
| WOM7777 | *H. erato* | *hydara* | Panama | Coclé | El Valle | 8.59323 | -80.14108 | B | -0.65 |
| WOM7796 | *H. erato* | *demophoon* | Panama | Coclé | El Valle | 8.59323 | -80.14108 | D | -0.56 |
| 18002 | *H. erato* | *hydara* | Panama | Daríen | Puerto Lara | 8.61355 | -78.13982 | A | -0.44 |
| 18008 | *H. erato* | *hydara* | Panama | Daríen | Puerto Lara | 8.61355 | -78.13982 | A | -0.57 |
| 18009 | *H. erato* | *hydara* | Panama | Daríen | Puerto Lara | 8.61355 | -78.13982 | A | -0.47 |
| 18022 | *H. erato* | *hydara x demophoon* | Panama | Daríen | Puerto Lara | 8.61355 | -78.13982 | B | -0.44 |
| 18027 | *H. erato* | *hydara* | Panama | Daríen | Puerto Lara | 8.61355 | -78.13982 | A | -0.48 |
| 18029 | *H. erato* | *hydara* | Panama | Daríen | Puerto Lara | 8.61355 | -78.13982 | A | -0.40 |
| 18035 | *H. erato* | *hydara x demophoon* | Panama | Daríen | Puerto Lara | 8.61355 | -78.13982 | B | -0.30 |
| 18036 | *H. erato* | *hydara* | Panama | Daríen | Puerto Lara | 8.61355 | -78.13982 | A | -0.36 |
| 18044 | *H. erato* | *hydara x demophoon* | Panama | Daríen | Puerto Lara | 8.61355 | -78.13982 | B | -0.63 |
| 18047 | *H. erato* | *hydara* | Panama | Daríen | Puerto Lara | 8.61355 | -78.13982 | A | -0.35 |
| 18049 | *H. erato* | *hydara x demophoon* | Panama | Daríen | Puerto Lara | 8.61355 | -78.13982 | B | -0.42 |
| 18054 | *H. erato* | *demophoon* | Panama | Daríen | Puerto Lara | 8.61355 | -78.13982 | D | -0.42 |
| 18059 | *H. erato* | *hydara x demophoon* | Panama | Daríen | Puerto Lara | 8.61355 | -78.13982 | A | -0.43 |
| 18060 | *H. erato* | *hydara* | Panama | Daríen | Puerto Lara | 8.61355 | -78.13982 | A | -0.48 |
| 18064 | *H. erato* | *hydara x demophoon* | Panama | Daríen | Puerto Lara | 8.61355 | -78.13982 | B | -0.43 |
| 18072 | *H. erato* | *hydara* | Panama | Daríen | Puerto Lara | 8.61355 | -78.13982 | A | -0.46 |
| 18073 | *H. erato* | *hydara* | Panama | Daríen | Puerto Lara | 8.61355 | -78.13982 | B | -0.44 |
| 18074 | *H. erato* | *hydara* | Panama | Daríen | Puerto Lara | 8.61355 | -78.13982 | A | -0.45 |
| 18080 | *H. erato* | *hydara* | Panama | Daríen | Puerto Lara | 8.61355 | -78.13982 | A | -0.46 |
| 18081 | *H. erato* | *hydara* | Panama | Daríen | Puerto Lara | 8.61355 | -78.13982 | A | -0.43 |
| 18082 | *H. erato* | *hydara* | Panama | Daríen | Puerto Lara | 8.61355 | -78.13982 | A | -0.46 |

| | | | | | | | | | |
|---|---|---|---|---|---|---|---|---|---|
| 18083 | *H. erato* | *hydara* | Panama | Daríen | Puerto Lara | 8.61355 | -78.13982 | A | -0.36 |
| WOM7120 | *H. erato* | *hydara* | Panama | Daríen | Agua Fria | 8.858867 | -78.22508 | A | -0.46 |
| WOM7122 | *H. erato* | *hydara* | Panama | Daríen | Agua Fria | 8.858867 | -78.22508 | A | -0.31 |
| WOM7123 | *H. erato* | *hydara* | Panama | Daríen | Agua Fria | 8.858867 | -78.22508 | A | -0.46 |
| WOM7125 | *H. erato* | *hydara* | Panama | Daríen | Agua Fria | 8.858867 | -78.22508 | A | -0.50 |
| WOM8862 | *H. erato* | *hydara* | Panama | Daríen | Agua Fria | 8.858867 | -78.22508 | A | -0.39 |
| WOM9000 | *H. erato* | *hydara* | Panama | Daríen | Agua Fria | 8.858867 | -78.22508 | B | -0.55 |
| WOM9001 | *H. erato* | *hydara* | Panama | Daríen | Agua Fria | 8.858867 | -78.22508 | B | -0.43 |
| WOM9002 | *H. erato* | *demophoon* | Panama | Daríen | Agua Fria | 8.858867 | -78.22508 | C | -0.40 |
| WOM9003 | *H. erato* | *hydara* | Panama | Daríen | Agua Fria | 8.858867 | -78.22508 | A | -0.41 |
| WOM9145 | *H. erato* | *hydara* | Panama | Daríen | Agua Fria | 8.858867 | -78.22508 | A | -0.64 |
| WOM9146 | *H. erato* | *hydara* | Panama | Daríen | Agua Fria | 8.858867 | -78.22508 | A | -0.52 |
| WOM9148 | *H. erato* | *hydara* | Panama | Daríen | Agua Fria | 8.858867 | -78.22508 | A | -0.42 |
| WOM9150 | *H. erato* | *hydara* | Panama | Daríen | Agua Fria | 8.858867 | -78.22508 | A | -0.40 |
| WOM9152 | *H. erato* | *hydara* | Panama | Daríen | Agua Fria | 8.858867 | -78.22508 | B | -0.48 |
| WOM9153 | *H. erato* | *hydara* | Panama | Daríen | Agua Fria | 8.858867 | -78.22508 | B | -0.56 |
| WOM9154 | *H. erato* | *hydara* | Panama | Daríen | Agua Fria | 8.858867 | -78.22508 | B | -0.47 |
| WOM9155 | *H. erato* | *hydara* | Panama | Daríen | Agua Fria | 8.858867 | -78.22508 | B | -0.62 |
| WOM9169 | *H. erato* | *hydara* | Panama | Daríen | Agua Fria | 8.858867 | -78.22508 | B | -0.45 |
| WOM8888 | *H. erato* | *hydara* | Panama | Panama | Ipeti | 8.972917 | -78.5106 | A | -0.38 |
| WOM8889 | *H. erato* | *hydara* | Panama | Panama | Ipeti | 8.972917 | -78.5106 | A | -0.42 |
| WOM8890 | *H. erato* | *hydara* | Panama | Panama | Ipeti | 8.972917 | -78.5106 | A | -0.55 |
| WOM8891 | *H. erato* | *hydara* | Panama | Panama | Ipeti | 8.972917 | -78.5106 | A | -0.46 |
| WOM8892 | *H. erato* | *hydara* | Panama | Panama | Ipeti | 8.972917 | -78.5106 | B | -0.45 |
| WOM8893 | *H. erato* | *hydara* | Panama | Panama | Ipeti | 8.972917 | -78.5106 | A | -0.33 |
| WOM8947 | *H. erato* | *hydara* | Panama | Panama | Ipeti | 8.972917 | -78.5106 | A | -0.47 |

| | | | | | | | | | |
|---|---|---|---|---|---|---|---|---|---|
| WOM8948 | *H. erato* | *hydara* | Panama | Panama | Ipeti | 8.972917 | -78.5106 | A | -0.40 |
| WOM8950 | *H. erato* | *hydara* | Panama | Panama | Ipeti | 8.972917 | -78.5106 | A | -0.56 |
| WOM8951 | *H. erato* | *hydara* | Panama | Panama | Ipeti | 8.972917 | -78.5106 | A | -0.42 |
| WOM8952 | *H. erato* | *hydara* | Panama | Panama | Ipeti | 8.972917 | -78.5106 | B | -0.55 |
| WOM8953 | *H. erato* | *hydara* | Panama | Panama | Ipeti | 8.972917 | -78.5106 | B | -0.58 |
| WOM8954 | *H. erato* | *hydara* | Panama | Panama | Ipeti | 8.972917 | -78.5106 | A | -0.47 |
| WOM8956 | *H. erato* | *hydara* | Panama | Panama | Ipeti | 8.972917 | -78.5106 | B | -0.53 |
| WOM8967 | *H. erato* | *hydara* | Panama | Panama | Ipeti | 8.972917 | -78.5106 | A | -0.39 |
| WOM8968 | *H. erato* | *hydara* | Panama | Panama | Ipeti | 8.972917 | -78.5106 | A | -0.46 |
| WOM8845 | *H. erato* | *demophoon* | Panama | Colón | Gamboa | 9.11605 | -79.69837 | D | -0.59 |
| WOM8846 | *H. erato* | *demophoon* | Panama | Colón | Gamboa | 9.11605 | -79.69837 | D | -0.53 |
| WOM8848 | *H. erato* | *demophoon* | Panama | Colón | Gamboa | 9.11605 | -79.69837 | D | -0.47 |
| WOM8849 | *H. erato* | *demophoon* | Panama | Colón | Gamboa | 9.11605 | -79.69837 | D | -0.60 |
| WOM8850 | *H. erato* | *demophoon* | Panama | Colón | Gamboa | 9.11605 | -79.69837 | D | -0.56 |
| WOM8851 | *H. erato* | *demophoon* | Panama | Colón | Gamboa | 9.11605 | -79.69837 | D | -0.58 |
| WOM8852 | *H. erato* | *demophoon* | Panama | Colón | Gamboa | 9.11605 | -79.69837 | D | -0.49 |
| WOM7577 | *H. erato* | *hydara* | Panama | Panama | Mangowichi | 9.137667 | -78.68952 | A | -0.48 |
| WOM7578 | *H. erato* | *hydara* | Panama | Panama | Mangowichi | 9.137667 | -78.68952 | A | -0.47 |
| WOM7579 | *H. erato* | *hydara* | Panama | Panama | Mangowichi | 9.137667 | -78.68952 | A | -0.59 |
| WOM7580 | *H. erato* | *venus* | Panama | Panama | Mangowichi | 9.137667 | -78.68952 | C | -0.21 |
| WOM7583 | *H. erato* | *hydara* | Panama | Panama | Mangowichi | 9.137667 | -78.68952 | B | -0.53 |
| WOM7584 | *H. erato* | *hydara* | Panama | Panama | Mangowichi | 9.137667 | -78.68952 | A | -0.53 |
| WOM7585 | *H. erato* | *hydara* | Panama | Panama | Mangowichi | 9.137667 | -78.68952 | B | -0.38 |
| WOM7587 | *H. erato* | *hydara* | Panama | Panama | Mangowichi | 9.137667 | -78.68952 | A | -0.53 |
| WOM7589 | *H. erato* | *hydara* | Panama | Panama | Mangowichi | 9.137667 | -78.68952 | A | -0.43 |
| WOM7591 | *H. erato* | *hydara* | Panama | Panama | Mangowichi | 9.137667 | -78.68952 | B | -0.48 |

| | | | | | | | | | |
|---|---|---|---|---|---|---|---|---|---|
| WOM7592 | *H. erato* | *hydara* | Panama | Panama | Mangowichi | 9.137667 | -78.68952 | B | -0.54 |
| WOM7593 | *H. erato* | *demophoon* | Panama | Panama | Mangowichi | 9.137667 | -78.68952 | D | -0.56 |
| WOM7594 | *H. erato* | *demophoon* | Panama | Panama | Mangowichi | 9.137667 | -78.68952 | D | -0.52 |
| WOM7595 | *H. erato* | *hydara* | Panama | Panama | Mangowichi | 9.137667 | -78.68952 | A | -0.61 |
| WOM7596 | *H. erato* | *hydara* | Panama | Panama | Mangowichi | 9.137667 | -78.68952 | B | -0.51 |
| WOM7597 | *H. erato* | *hydara* | Panama | Panama | Mangowichi | 9.137667 | -78.68952 | A | -0.22 |
| WOM7598 | *H. erato* | *hydara* | Panama | Panama | Mangowichi | 9.137667 | -78.68952 | A | -0.43 |
| WOM7600 | *H. erato* | *hydara* | Panama | Panama | Mangowichi | 9.137667 | -78.68952 | B | -0.51 |
| WOM7601 | *H. erato* | *hydara* | Panama | Panama | Mangowichi | 9.137667 | -78.68952 | B | -0.65 |
| WOM7602 | *H. erato* | *hydara* | Panama | Panama | Mangowichi | 9.137667 | -78.68952 | B | -0.46 |
| WOM7603 | *H. erato* | *hydara* | Panama | Panama | Mangowichi | 9.137667 | -78.68952 | A | -0.50 |
| WOM7641 | *H. erato* | *hydara* | Panama | Panama | Mangowichi | 9.137667 | -78.68952 | A | -0.39 |
| WOM7643 | *H. erato* | *hydara* | Panama | Panama | Mangowichi | 9.137667 | -78.68952 | B | -0.42 |
| WOM7644 | *H. erato* | *hydara* | Panama | Panama | Mangowichi | 9.137667 | -78.68952 | A | -0.48 |
| WOM7648 | *H. erato* | *hydara* | Panama | Panama | Mangowichi | 9.137667 | -78.68952 | A | -0.45 |
| WOM7650 | *H. erato* | *hydara* | Panama | Panama | Mangowichi | 9.137667 | -78.68952 | A | -0.52 |
| WOM7653 | *H. erato* | *hydara* | Panama | Panama | Mangowichi | 9.137667 | -78.68952 | B | -0.50 |
| WOM7654 | *H. erato* | *hydara* | Panama | Panama | Mangowichi | 9.137667 | -78.68952 | A | -0.44 |
| WOM7655 | *H. erato* | *hydara* | Panama | Panama | Mangowichi | 9.137667 | -78.68952 | A | -0.46 |
| WOM7656 | *H. erato* | *hydara* | Panama | Panama | Mangowichi | 9.137667 | -78.68952 | B | -0.46 |
| WOM7658 | *H. erato* | *hydara* | Panama | Panama | Mangowichi | 9.137667 | -78.68952 | B | -0.49 |
| WOM7659 | *H. erato* | *hydara* | Panama | Panama | Mangowichi | 9.137667 | -78.68952 | A | -0.46 |
| WOM7660 | *H. erato* | *hydara* | Panama | Panama | Mangowichi | 9.137667 | -78.68952 | B | -0.52 |
| WOM7661 | *H. erato* | *hydara* | Panama | Panama | Mangowichi | 9.137667 | -78.68952 | A | -0.56 |
| WOM7716 | *H. erato* | *demophoon* | Panama | Panama | Mangowichi | 9.137667 | -78.68952 | D | -0.43 |
| WOM7787 | *H. erato* | *demophoon* | Panama | Panama | Tocumen | 9.200417 | -79.39525 | D | -0.54 |

| | | | | | | | | | |
|---|---|---|---|---|---|---|---|---|---|
| WOM7790 | *H. erato* | *demophoon* | Panama | Panama | Tocumen | 9.200417 | -79.39525 | D | -0.50 |
| WOM7801 | *H. erato* | *demophoon* | Panama | Panama | Tocumen | 9.200417 | -79.39525 | D | -0.50 |
| WOM7804 | *H. erato* | *demophoon* | Panama | Panama | Tocumen | 9.200417 | -79.39525 | D | -0.55 |
| WOM7806 | *H. erato* | *demophoon* | Panama | Panama | Tocumen | 9.200417 | -79.39525 | D | -0.45 |
| WOM7807 | *H. erato* | *demophoon* | Panama | Panama | Tocumen | 9.200417 | -79.39525 | D | -0.55 |
| WOM7808 | *H. erato* | *demophoon* | Panama | Panama | Tocumen | 9.200417 | -79.39525 | D | -0.46 |
| WOM7810 | *H. erato* | *hydara* | Panama | Panama | Tocumen | 9.200417 | -79.39525 | B | -0.49 |
| WOM7812 | *H. erato* | *hydara* | Panama | Panama | Tocumen | 9.200417 | -79.39525 | B | -0.53 |
| WOM7813 | *H. erato* | *hydara* | Panama | Panama | Tocumen | 9.200417 | -79.39525 | B | -0.43 |
| WOM7822 | *H. erato* | *demophoon* | Panama | Panama | Tocumen | 9.200417 | -79.39525 | D | -0.51 |
| WOM7823 | *H. erato* | *demophoon* | Panama | Panama | Tocumen | 9.200417 | -79.39525 | D | -0.50 |
| WOM7824 | *H. erato* | *demophoon* | Panama | Panama | Tocumen | 9.200417 | -79.39525 | D | -0.53 |
| WOM7828 | *H. erato* | *demophoon* | Panama | Panama | Tocumen | 9.200417 | -79.39525 | D | -0.43 |
| WOM7829 | *H. erato* | *demophoon* | Panama | Panama | Tocumen | 9.200417 | -79.39525 | D | -0.51 |
| WOM7831 | *H. erato* | *demophoon* | Panama | Panama | Tocumen | 9.200417 | -79.39525 | D | -0.52 |
| WOM7832 | *H. erato* | *demophoon* | Panama | Panama | Tocumen | 9.200417 | -79.39525 | D | -0.61 |
| WOM7833 | *H. erato* | *demophoon* | Panama | Panama | Tocumen | 9.200417 | -79.39525 | D | -0.41 |
| WOM7834 | *H. erato* | *demophoon* | Panama | Panama | Tocumen | 9.200417 | -79.39525 | D | -0.47 |
| WOM7836 | *H. erato* | *demophoon* | Panama | Panama | Tocumen | 9.200417 | -79.39525 | D | -0.58 |
| WOM7697 | *H. erato* | *hydara* | Panama | Panama | El llano | 9.237533 | -78.95347 | B | -0.53 |
| WOM7698 | *H. erato* | *hydara* | Panama | Panama | El llano | 9.237533 | -78.95347 | B | -0.47 |
| WOM7699 | *H. erato* | *hydara* | Panama | Panama | El llano | 9.237533 | -78.95347 | B | -0.52 |
| WOM7700 | *H. erato* | *demophoon* | Panama | Panama | El llano | 9.237533 | -78.95347 | D | -0.43 |
| WOM7701 | *H. erato* | *hydara* | Panama | Panama | El llano | 9.237533 | -78.95347 | B | -0.41 |
| WOM7702 | *H. erato* | *demophoon* | Panama | Panama | El llano | 9.237533 | -78.95347 | D | -0.47 |
| WOM7703 | *H. erato* | *demophoon* | Panama | Panama | El llano | 9.237533 | -78.95347 | D | -0.44 |

| | | | | | | | | | |
|---|---|---|---|---|---|---|---|---|---|
| WOM7704 | *H. erato* | *hydara* | Panama | Panama | El llano | 9.237533 | -78.95347 | B | -0.41 |
| WOM7706 | *H. erato* | *demophoon* | Panama | Panama | El llano | 9.237533 | -78.95347 | D | -0.42 |
| WOM7707 | *H. erato* | *venus* | Panama | Panama | El llano | 9.237533 | -78.95347 | C | -0.47 |
| WOM7708 | *H. erato* | *hydara* | Panama | Panama | El llano | 9.237533 | -78.95347 | B | -0.44 |
| WOM7709 | *H. erato* | *hydara* | Panama | Panama | El llano | 9.237533 | -78.95347 | B | -0.39 |
| WOM7710 | *H. erato* | *hydara* | Panama | Panama | El llano | 9.237533 | -78.95347 | A | -0.50 |
| WOM7711 | *H. erato* | *hydara* | Panama | Panama | El llano | 9.237533 | -78.95347 | A | -0.59 |
| WOM7712 | *H. erato* | *hydara* | Panama | Panama | El llano | 9.237533 | -78.95347 | B | -0.52 |
| WOM7713 | *H. erato* | *demophoon* | Panama | Panama | El llano | 9.237533 | -78.95347 | D | -0.41 |
| WOM7859 | *H. erato* | *hydara* | Panama | Panama | El llano | 9.237533 | -78.95347 | B | -0.64 |
| WOM7860 | *H. erato* | *hydara* | Panama | Panama | El llano | 9.237533 | -78.95347 | A | -0.40 |
| WOM7861 | *H. erato* | *hydara* | Panama | Panama | El llano | 9.237533 | -78.95347 | B | -0.37 |
| WOM7862 | *H. erato* | *hydara* | Panama | Panama | El llano | 9.237533 | -78.95347 | B | -0.44 |
| WOM7863 | *H. erato* | *hydara* | Panama | Panama | El llano | 9.237533 | -78.95347 | A | -0.43 |
| WOM7865 | *H. erato* | *hydara* | Panama | Panama | El llano | 9.237533 | -78.95347 | B | -0.40 |
| WOM7960 | *H. erato* | *hydara* | Panama | Panama | El llano | 9.237533 | -78.95347 | B | -0.55 |
| WOM7961 | *H. erato* | *demophoon* | Panama | Panama | El llano | 9.237533 | -78.95347 | D | -0.42 |
| WOM8858 | *H. erato* | *hydara* | Panama | Panama | El llano | 9.237533 | -78.95347 | B | -0.62 |
| WOM8859 | *H. erato* | *demophoon* | Panama | Panama | El llano | 9.237533 | -78.95347 | D | -0.52 |
| WOM8861 | *H. erato* | *hydara* | Panama | Panama | El llano | 9.237533 | -78.95347 | A | -0.42 |
| WOM8864 | *H. erato* | *hydara* | Panama | Panama | El llano | 9.237533 | -78.95347 | A | -0.54 |
| WOM9067 | *H. erato* | *demophoon* | Panama | Panama | El llano | 9.237533 | -78.95347 | D | -0.50 |
| WOM9068 | *H. erato* | *hydara* | Panama | Panama | El llano | 9.237533 | -78.95347 | B | -0.39 |
| 15N156 | *H. melpomene* | *vulcanus* | Colombia | Valle del Cauca | Queremal | 3.53199 | -76.75461 | C | 0.21 |
| 574 | *H. melpomene* | *vulcanus* | Colombia | Valle del Cauca | Rio Bravo | 3.88274 | -76.57716 | C | -0.18 |

| | | | | | | | | | |
|---|---|---|---|---|---|---|---|---|---|
| 710 | *H. melpomene* | *vulcanus* | Colombia | Valle del Cauca | Rio Bravo | 3.88274 | -76.57716 | C | -0.05 |
| 711 | *H. melpomene* | *vulcanus* | Colombia | Valle del Cauca | Rio Bravo | 3.88274 | -76.57716 | C | -0.24 |
| 713 | *H. melpomene* | *vulcanus* | Colombia | Valle del Cauca | Rio Bravo | 3.88274 | -76.57716 | C | 0.00 |
| 714 | *H. melpomene* | *vulcanus* | Colombia | Valle del Cauca | Rio Bravo | 3.88274 | -76.57716 | C | -0.06 |
| 747 | *H. melpomene* | *vulcanus* | Colombia | Valle del Cauca | Rio Bravo | 3.88274 | -76.57716 | C | 0.08 |
| 749 | *H. melpomene* | *vulcanus* | Colombia | Valle del Cauca | Rio Bravo | 3.88274 | -76.57716 | C | -0.14 |
| 750 | *H. melpomene* | *vulcanus* | Colombia | Valle del Cauca | Rio Bravo | 3.88274 | -76.57716 | C | 0.04 |
| 1001 | *H. melpomene* | *vulcanus* | Colombia | Valle del Cauca | Rio Bravo | 3.88274 | -76.57716 | C | -0.10 |
| 457 | *H. melpomene* | *vulcanus* | Colombia | Valle del Cauca | Ladrilleros | 3.93944 | -77.36889 | C | -0.31 |
| 636 | *H. melpomene* | *vulcanus* | Colombia | Valle del Cauca | Ladrilleros | 3.93944 | -77.36889 | C | -0.22 |
| 647 | *H. melpomene* | *vulcanus* | Colombia | Valle del Cauca | Ladrilleros | 3.93944 | -77.36889 | C | -0.17 |
| 15N306 | *H. melpomene* | *vulcanus* | Colombia | Chocó | Amargal | 5.57187 | -77.50211 | C | 0.18 |
| 16N091 | *H. melpomene* | *vulcanus* | Colombia | Chocó | Potes | 6.3495 | -77.36697 | C | -0.08 |
| 16N092 | *H. melpomene* | *vulcanus* | Colombia | Chocó | Potes | 6.3495 | -77.36697 | C | -0.25 |
| 16N093 | *H. melpomene* | *vulcanus* | Colombia | Chocó | Potes | 6.3495 | -77.36697 | C | 0.02 |
| 16N094 | *H. melpomene* | *vulcanus* | Colombia | Chocó | Potes | 6.3495 | -77.36697 | C | -0.15 |
| 16N095 | *H. melpomene* | *vulcanus* | Colombia | Chocó | Potes | 6.3495 | -77.36697 | C | -0.12 |

| | | | | | | | | | |
|---|---|---|---|---|---|---|---|---|---|
| 16N059 | *H. melpomene* | *vulcanus* | Colombia | Chocó | Cocalito | 6.38504 | -77.40251 | C | -0.10 |
| 16N063 | *H. melpomene* | *vulcanus* | Colombia | Chocó | Cocalito | 6.38504 | -77.40251 | C | -0.23 |
| 16N066 | *H. melpomene* | *vulcanus* | Colombia | Chocó | Cocalito | 6.38504 | -77.40251 | C | -0.37 |
| 16N068 | *H. melpomene* | *vulcanus* | Colombia | Chocó | Cocalito | 6.38504 | -77.40251 | C | -0.04 |
| 16N072 | *H. melpomene* | *vulcanus x melpomene* | Colombia | Chocó | Cocalito | 6.38504 | -77.40251 | B | -0.01 |
| 16N073 | *H. melpomene* | *vulcanus x melpomene* | Colombia | Chocó | Cocalito | 6.38504 | -77.40251 | B | -0.30 |
| 16N077 | *H. melpomene* | *vulcanus* | Colombia | Chocó | Cocalito | 6.38504 | -77.40251 | C | 0.03 |
| 16N078 | *H. melpomene* | *vulcanus* | Colombia | Chocó | Cocalito | 6.38504 | -77.40251 | C | -0.30 |
| 16N080 | *H. melpomene* | *vulcanus* | Colombia | Chocó | Cocalito | 6.38504 | -77.40251 | C | -0.03 |
| 16N020 | *H. melpomene* | *vulcanus* | Colombia | Chocó | Playa Flores | 6.38765 | -77.38557 | C | -0.19 |
| 16N047 | *H. melpomene* | *vulcanus* | Colombia | Chocó | Playa Flores | 6.38765 | -77.38557 | C | 0.01 |
| 16N048 | *H. melpomene* | *vulcanus* | Colombia | Chocó | Playa Flores | 6.38765 | -77.38557 | C | 0.02 |
| 15N451 | *H. melpomene* | *melpomene* | Panama | Daríen | Jaque | 7.48767 | -78.12733 | B | -0.42 |
| 18092 | *H. melpomene* | *melpomene x rosina* | Panama | Daríen | Santa Librada | 8.27971 | -77.80982 | B | -0.51 |
| 18098 | *H. melpomene* | *melpomene x rosina* | Panama | Daríen | Santa Librada | 8.27971 | -77.80982 | B | -0.37 |
| 18147 | *H. melpomene* | *melpomene x rosina* | Panama | Daríen | Santa Librada | 8.27971 | -77.80982 | B | -0.44 |
| 18149 | *H. melpomene* | *melpomene x rosina* | Panama | Daríen | Santa Librada | 8.27971 | -77.80982 | B | -0.38 |

| | | | | | | | | | |
|---|---|---|---|---|---|---|---|---|---|
| 18156 | *H. melpomene* | *melpomene x rosina* | Panama | Daríen | Santa Librada | 8.27971 | -77.80982 | B | -0.49 |
| 18157 | *H. melpomene* | *melpomene x rosina* | Panama | Daríen | Santa Librada | 8.27971 | -77.80982 | B | -0.53 |
| 18159 | *H. melpomene* | *melpomene x rosina* | Panama | Daríen | Santa Librada | 8.27971 | -77.80982 | B | -0.49 |
| 18161 | *H. melpomene* | *melpomene* | Panama | Daríen | Santa Librada | 8.27971 | -77.80982 | A | -0.36 |
| 18169 | *H. melpomene* | *melpomene x rosina* | Panama | Daríen | Santa Librada | 8.27971 | -77.80982 | B | -0.44 |
| 18173 | *H. melpomene* | *melpomene* | Panama | Daríen | Santa Librada | 8.27971 | -77.80982 | A | -0.43 |
| 18174 | *H. melpomene* | *melpomene x rosina* | Panama | Daríen | Santa Librada | 8.27971 | -77.80982 | B | -0.44 |
| 18175 | *H. melpomene* | *melpomene* | Panama | Daríen | Santa Librada | 8.27971 | -77.80982 | A | -0.40 |
| 18194 | *H. melpomene* | *melpomene x rosina* | Panama | Daríen | Santa Librada | 8.27971 | -77.80982 | B | -0.50 |
| 18195 | *H. melpomene* | *melpomene x rosina* | Panama | Daríen | Santa Librada | 8.27971 | -77.80982 | B | -0.55 |
| WOM9170 | *H. melpomene* | *rosina* | Panama | Coclé | El Valle | 8.59323 | -80.14108 | D | -0.61 |
| WOM9196 | *H. melpomene* | *rosina* | Panama | Coclé | El Valle | 8.59323 | -80.14108 | D | -0.56 |
| WOM9199 | *H. melpomene* | *rosina* | Panama | Coclé | El Valle | 8.59323 | -80.14108 | D | -0.52 |
| WOM9209 | *H. melpomene* | *rosina* | Panama | Coclé | El Valle | 8.59323 | -80.14108 | D | -0.53 |
| WOM9210 | *H. melpomene* | *rosina* | Panama | Coclé | El Valle | 8.59323 | -80.14108 | D | -0.68 |
| 18017 | *H. melpomene* | *vulcanus x rosina* | Panama | Daríen | Puerto Lara | 8.61355 | -78.13982 | C | -0.53 |
| 18019 | *H. melpomene* | *vulcanus* | Panama | Daríen | Puerto Lara | 8.61355 | -78.13982 | C | -0.44 |

| | | | | | | | | | |
|---|---|---|---|---|---|---|---|---|---|
| 18023 | *H. melpomene* | *melpomene x rosina* | Panama | Daríen | Puerto Lara | 8.61355 | -78.13982 | B | -0.31 |
| 18030 | *H. melpomene* | *melpomene* | Panama | Daríen | Puerto Lara | 8.61355 | -78.13982 | A | -0.19 |
| 18040 | *H. melpomene* | *melpomene x rosina* | Panama | Daríen | Puerto Lara | 8.61355 | -78.13982 | B | -0.48 |
| 18043 | *H. melpomene* | *melpomene x rosina* | Panama | Daríen | Puerto Lara | 8.61355 | -78.13982 | C | -0.60 |
| 18048 | *H. melpomene* | *melpomene x rosina* | Panama | Daríen | Puerto Lara | 8.61355 | -78.13982 | B | -0.44 |
| 18051 | *H. melpomene* | *melpomene x rosina* | Panama | Daríen | Puerto Lara | 8.61355 | -78.13982 | B | -0.38 |
| 18053 | *H. melpomene* | *melpomene x rosina* | Panama | Daríen | Puerto Lara | 8.61355 | -78.13982 | B | -0.45 |
| 18065 | *H. melpomene* | *melpomene x rosina* | Panama | Daríen | Puerto Lara | 8.61355 | -78.13982 | B | -0.41 |
| 18066 | *H. melpomene* | *melpomene x rosina* | Panama | Daríen | Puerto Lara | 8.61355 | -78.13982 | B | -0.50 |
| WOM9157 | *H. melpomene* | *melpomene* | Panama | Daríen | Agua Fria | 8.85887 | -78.22508 | B | -0.59 |
| WOM9158 | *H. melpomene* | *melpomene* | Panama | Daríen | Agua Fria | 8.85887 | -78.22508 | B | -0.39 |
| WOM9159 | *H. melpomene* | *melpomene* | Panama | Daríen | Agua Fria | 8.85887 | -78.22508 | B | -0.40 |
| WOM9160 | *H. melpomene* | *melpomene* | Panama | Daríen | Agua Fria | 8.85887 | -78.22508 | B | -0.58 |
| WOM9161 | *H. melpomene* | *rosina* | Panama | Daríen | Agua Fria | 8.85887 | -78.22508 | C | -0.46 |
| WOM9166 | *H. melpomene* | *melpomene* | Panama | Daríen | Agua Fria | 8.85887 | -78.22508 | B | -0.39 |
| WOM9167 | *H. melpomene* | *melpomene* | Panama | Daríen | Agua Fria | 8.85887 | -78.22508 | B | -0.45 |
| WOM9168 | *H. melpomene* | *melpomene* | Panama | Daríen | Agua Fria | 8.85887 | -78.22508 | B | -0.46 |

| | | | | | | | | | |
|---|---|---|---|---|---|---|---|---|---|
| WOM8895 | *H. melpomene* | *melpomene* | Panama | Panama | Ipeti | 8.97292 | -78.5106 | B | -0.51 |
| WOM8896 | *H. melpomene* | *melpomene* | Panama | Panama | Ipeti | 8.97292 | -78.5106 | B | -0.56 |
| WOM8897 | *H. melpomene* | *vulcanus* | Panama | Panama | Ipeti | 8.97292 | -78.5106 | C | -0.53 |
| WOM8898 | *H. melpomene* | *melpomene* | Panama | Panama | Ipeti | 8.97292 | -78.5106 | B | -0.66 |
| WOM8900 | *H. melpomene* | *melpomene* | Panama | Panama | Ipeti | 8.97292 | -78.5106 | B | -0.40 |
| WOM8901 | *H. melpomene* | *melpomene* | Panama | Panama | Ipeti | 8.97292 | -78.5106 | B | -0.37 |
| WOM8902 | *H. melpomene* | *melpomene* | Panama | Panama | Ipeti | 8.97292 | -78.5106 | B | -0.40 |
| WOM8903 | *H. melpomene* | *melpomene* | Panama | Panama | Ipeti | 8.97292 | -78.5106 | B | -0.44 |
| WOM8904 | *H. melpomene* | *vulcanus* | Panama | Panama | Ipeti | 8.97292 | -78.5106 | C | -0.42 |
| WOM8958 | *H. melpomene* | *melpomene* | Panama | Panama | Ipeti | 8.97292 | -78.5106 | A | -0.40 |
| WOM8959 | *H. melpomene* | *melpomene* | Panama | Panama | Ipeti | 8.97292 | -78.5106 | B | -0.33 |
| WOM8960 | *H. melpomene* | *melpomene* | Panama | Panama | Ipeti | 8.97292 | -78.5106 | B | -0.38 |
| WOM8961 | *H. melpomene* | *melpomene* | Panama | Panama | Ipeti | 8.97292 | -78.5106 | B | -0.43 |
| WOM8962 | *H. melpomene* | *melpomene* | Panama | Panama | Ipeti | 8.97292 | -78.5106 | B | -0.66 |
| WOM8963 | *H. melpomene* | *melpomene* | Panama | Panama | Ipeti | 8.97292 | -78.5106 | B | -0.56 |
| WOM8965 | *H. melpomene* | *melpomene* | Panama | Panama | Ipeti | 8.97292 | -78.5106 | A | -0.50 |
| WOM8971 | *H. melpomene* | *melpomene* | Panama | Panama | Ipeti | 8.97292 | -78.5106 | B | -0.67 |

| | | | | | | | | | |
|---|---|---|---|---|---|---|---|---|---|
| WOM7576 | *H. melpomene* | *melpomene* | Panama | Panama | Mangowichi | 9.13767 | -78.68952 | A | -0.46 |
| WOM7586 | *H. melpomene* | *melpomene* | Panama | Panama | Mangowichi | 9.13767 | -78.68952 | B | -0.45 |
| WOM7588 | *H. melpomene* | *melpomene* | Panama | Panama | Mangowichi | 9.13767 | -78.68952 | B | -0.40 |
| WOM7590 | *H. melpomene* | *melpomene* | Panama | Panama | Mangowichi | 9.13767 | -78.68952 | A | -0.40 |
| WOM7599 | *H. melpomene* | *melpomene* | Panama | Panama | Mangowichi | 9.13767 | -78.68952 | A | -0.51 |
| WOM7642 | *H. melpomene* | *melpomene* | Panama | Panama | Mangowichi | 9.13767 | -78.68952 | B | -0.59 |
| WOM7645 | *H. melpomene* | *melpomene* | Panama | Panama | Mangowichi | 9.13767 | -78.68952 | B | -0.52 |
| WOM7646 | *H. melpomene* | *melpomene* | Panama | Panama | Mangowichi | 9.13767 | -78.68952 | B | -0.53 |
| WOM7647 | *H. melpomene* | *melpomene* | Panama | Panama | Mangowichi | 9.13767 | -78.68952 | B | -0.48 |
| WOM7649 | *H. melpomene* | *melpomene* | Panama | Panama | Mangowichi | 9.13767 | -78.68952 | A | -0.51 |
| WOM7652 | *H. melpomene* | *melpomene* | Panama | Panama | Mangowichi | 9.13767 | -78.68952 | A | -0.41 |
| WOM7657 | *H. melpomene* | *rosina* | Panama | Panama | Mangowichi | 9.13767 | -78.68952 | D | -0.46 |
| WOM7815 | *H. melpomene* | *melpomene* | Panama | Panama | Tocumen | 9.20042 | -79.39525 | B | -0.49 |
| WOM7816 | *H. melpomene* | *rosina* | Panama | Panama | Tocumen | 9.20042 | -79.39525 | D | -0.45 |
| WOM7817 | *H. melpomene* | *rosina* | Panama | Panama | Tocumen | 9.20042 | -79.39525 | D | -0.56 |
| WOM8860 | *H. melpomene* | *melpomene* | Panama | Panama | El llano | 9.23753 | -78.95347 | B | -0.47 |

**Table S5.** Repeatability test for measures of red and blue colour channels from photographs.

| Measurement | d.f. | Mean square groups | Mean square error | F value | *P* | Repeatability |
|---|---|---|---|---|---|---|
| Wing region 1, blue mean | 25, 26 | 1298.8 | 9.3 | 139.3 | <0.001 | 0.986 |
| Wing region 2, blue mean | 25, 26 | 1763.6 | 23.8 | 74.13 | <0.001 | 0.973 |
| Wing region 1, red mean | 25, 26 | 310.59 | 2.05 | 151.7 | <0.001 | 0.987 |
| Wing region 2, red mean | 25, 26 | 396.6 | 2.3 | 171.3 | <0.001 | 0.988 |

**Table S6** – Likelihood ratio tests to compare three different cline models, sigmoidal (Sig), symmetrical stepped (Sstep) and asymmetrical stepped (Astep), for iridescence, the west Colombian yellow bar allele frequency ($y_{wc}$), and admixture proportions. The test statistic (2ΔML) and degrees of freedom (d.f.) are given.

| Species | Trait | Model comparison | 2ΔML | d.f. | *P* |
|---|---|---|---|---|---|
| *Heliconius erato* | Iridescence | Sig x Sstep | 9.004 | 2 | 0.01109 |
| | | Sig x Astep | 22.572 | 4 | 0.000154 |
| | | Sstep x Astep | 13.568 | 2 | 0.001132 |
| | $y_{wc}$ | Sig x Sstep | 0 | 2 | 1 |
| | | Sig x Astep | 5.388 | 4 | 0.2498 |
| | | Sstep x Astep | 5.388 | 2 | 0.06761 |
| | Admixture proportion | Sig x Sstep | 10.602 | 2 | 0.004987 |
| | | Sig x Astep | 24.396 | 4 | 6.65E-05 |
| | | Sstep x Astep | 13.794 | 2 | 0.001011 |
| *Heliconius melpomene* | Iridescence | Sig x Sstep | 4.39 | 2 | 0.1114 |
| | | Sig x Astep | 4.84 | 4 | 0.3041 |
| | | Sstep x Astep | 0.45 | 2 | 0.7985 |
| | $y_{wc}$ | Sig x Sstep | 0 | 2 | 1 |
| | | Sig x Astep | 1.302 | 4 | 0.861 |
| | | Sstep x Astep | 1.302 | 2 | 0.5215 |
| | Admixture proportion | Sig x Sstep | 0 | 2 | 1 |
| | | Sig x Astep | 0 | 4 | 1 |
| | | Sstep x Astep | 0 | 2 | 1 |

**Table S7 -** Summary of the linear and quadratic polynomial models fit to pairs of traits within species and each trait between species. Note that this analysis could not be used to compare the admixture clines between the species because many sample locations only included genetic data for one of the species.

| | | Linear | | | Quadratic | | |
|---|---|---|---|---|---|---|---|
| | Trait(s) | *F* | *P* | $r^2$ | *F* | *P* | $r^2$ |
| *H. erato* | Yellow bar ~ Iridescence | 1083.0 | 2.2e-16 | 0.981 | 608.5 | 2.2e-16 | 0.983 |
| | Iridescence ~ Admixture | 744.4 | 7.3e-13 | 0.981 | 376.1 | 1.4e-11 | 0.981 |
| | Admixture ~ Yellow bar | 1753.0 | 2.9e-15 | 0.992 | 827.0 | 1.4e-13 | 0.991 |
| *H. melpomene* | Yellow bar ~ Iridescence | 46.1 | 7.9e-05 | 0.819 | 44.12 | 4.7e-05 | 0.896 |
| | Iridescence ~ Admixture | 5.4 | 0.102 | 0.523 | 2.67 | 0.272 | 0.455 |
| | Admixture ~ Yellow bar | 5.5 | 0.079 | 0.472 | 3.51 | 0.163 | 0.501 |
| Between species | Yellow bar | 58.96 | 3.1e-05 | 0.853 | 144.1 | 5.3e-07 | 0.966 |
| | Iridescence | 127.9 | 1.3e-06 | 0.927 | 58.21 | 1.7e-05 | 0.936 |
